# Domain-general neural effects of associative learning and expectations on pain and hedonic taste perception

**DOI:** 10.1101/2025.04.10.648205

**Authors:** Yili Zhao, In-Seon Lee, Margaret Rose-McCandlish, Qingbao Yu, Dominik Mischkowski, Jason Avery, John E Ingeholm, Richard Reynolds, Gang Chen, Lauren Atlas

## Abstract

Predictive cues significantly influence perception through associative learning. However, it is unknown whether circuits are conserved across domains. We investigated how associative learning influences perceived intensity and valence of pain and hedonic taste and whether the mechanisms that support expectancy-based modulation vary as a function of aversiveness and modality. Sixty participants were randomly assigned to receive either painful heat, unpleasant liquid saline, or pleasant liquid sucrose during fMRI scanning. Following conditioning, cues that were initially associated with low or high intensity outcomes were intermittently followed by stimuli calibrated to elicit medium intensity ratings. Learned cues modulated expectations and subjective outcomes similarly across domains. Consistent with this, the orbitofrontal cortex exhibited domain-general anticipatory activation. The left anterior insula mediated domain-general cue effects on subjective intensity, whereas the thalamus mediated cue effects on subjective valence. Notably, these two regions were nearly identical to those previously implicated in mediating cue effects on pain (nearly identical to those previously implicated in cue effects on mediating pain (Atlas et al., 2010). Pain specificity was evident when we measured variations in stimulus intensity, whether we used univariate or multivariate approaches, but there was minimal evidence of specificity by modality or aversiveness when we examined cue effects on medium trials. These findings suggest that shared neural circuits mediate the effects of learned expectations on perception, linking pain with other areas of affective processing and perception across domains.

## Introduction

Expectations profoundly shape human perception. Humans, animals, and low-level organisms learn to associate predictive cues with outcomes through associative learning. Research on reward learning highlights the role of brain regions including the striatum and orbitofrontal cortex (OFC) in generating and updating predictions about primary and secondary reinforcers (O’Doherty, 2004; Sescousse et al., 2013; Rolls, 2015). The OFC and the broader ventromedial prefrontal cortex (VMPFC) is thought to provide a “common currency” of value (Levy & Glimcher, 2016), comparing different outcomes to maximize reward and avoid punishment. Consistent with this idea, meta-analyses illustrate a common neural representation of expected rewards within specific subregions of the OFC (Levy & Glimcher, 2012) . Other work highlights the crucial role of this region in modulating pain perception in the context of cue-based pain modulation and placebo analgesia (Roy et al., 2012; Atlas & Wager, 2014), and lesions of this region alter the effects of cues on expectations and pain unpleasantness (Motzkin et al., 2023). One study which re-analyzed functional neuroimaging studies of placebo analgesia and placebo anxiolysis found that that placebo effects on pain and emotional processing are both linked to responses in the lateral OFC (Petrovic et al., 2010). These findings may be consistent with a general role for the OFC in the formation of expectations during associative learning (Rudebeck & Murray, 2014). Likewise, other regions involved in valuation such as the striatum and amygdala have been implicated in learning about both appetitive and aversive outcomes, as well as pain (Seymour et al., 2005; Delgado, 2007; Atlas et al., 2016).

Once associations are established, the presence of a cue can prompt predictions which can in turn alter perceptions of stimuli themselves. We refer to such modulation as expectancy effects. Expectancy effects have been demonstrated in various contexts, including pain (Atlas & Wager, 2012; Fields, 2018), taste (Nitschke, Dixon, et al., 2006), visual perception (Balcetis & Dunning, 2006), and many other domains (Kirsch, 1997; Heinz et al., 2013). Only a few studies have directly measured whether the brain mechanisms that support expectancy effects vary across domains (Fazeli & Büchel, 2018; Sharvit et al., 2018; Lee et al., 2024). Using within-subjects designs, Sharvit et al. (2018) and Fazili and Buechel (2018) crossed within-modality predictive cues for pain with cues that predicted outcomes in one other aversive domain (i.e. disgusting odors and negative pictures, respectively). Both studies found that the anterior insula is more engaged in domain-general processing of expectancy and prediction error, while the posterior insula is more specific to pain processing (Sharvit et al., 2019). Thus the insula is a second region that is highly implicated in domain-general expectancy effects, in addition to the OFC. We note that most of these studies presented explicit cues and/or clear instructions to participants to induce expectations, raising the question of whether these findings hold when expectations arise purely from associative learning. In addition, as prior work has focused on aversive outcomes, it is unknown whether these findings extend to appetitive outcomes as well. For this reason, we measured both subjective intensity (regardless of valence) and valence to capture differences between appetitive and aversive outcomes.

We asked whether the brain mechanisms underlying expectancy effects on aversive and appetitive outcomes are domain-general or domain-specific. We compared pain with hedonic taste due to their shared evolutionary importance, neural activations, and chemosensory characteristics (Breslin, 2013; Riello et al., 2019; Aroke et al., 2020; Sandri et al., 2021). Pain and taste both depend heavily on the insula, which contains gustatory cortex (Legrain et al., 2011; Corradi-Dell’Acqua et al., 2016; Avery et al., 2020) and is involved in expectancy effects on both domains as discussed above. Machine learning has been used to determine neural representations supporting pain-specific activity, including the Neurologic Pain Signature (NPS; Wager et al., 2013), which is sensitive and specific to nociceptive pain, and the Stimulus-Intensity Independent Pain Signature (SIIPS; Woo et al., 2017), which predicts subjective pain independent of stimulus intensity. Recently, Ceko et al (2022) identified a domain-general pattern that predicts subjective unpleasantness across aversive stimuli (i.e., mechanical pain, thermal pain, noise, and negative pictures) and is not affected by positive affect. These signatures can be used to test whether associative learning and expectancy effects modulate domain-specific and domain-general processes. Furthermore, the ability of these patterns to distinguish between pain and taste modalities, and to classify experiences as aversive or appetitive, merits further examination.

In this study, we conducted a between-groups fMRI experiment in which predictive cues were paired with painful heat, aversive taste (saline solution), or appetitive taste (sucrose solution). Cues were presented without contingency instructions to measure the effects of associative learning on expectancy and perception. This design enabled us to investigate whether predictive cues modulated perception similarly across domains (pain vs.taste; appetitive vs. aversive outcomes). Specifically, we tested the preregistered hypotheses that our task would engage three dissociable neural circuits: 1) nociception-specific networks that are sensitive to changes in temperature but not cues or taste (posterior insula and secondary somatosensory cortex); 2) circuits that are sensitive to modality and expectancy (anterior insula); and 3) domain-general networks that support learning and expectancy across domains (OFC, VMPFC, amygdala, striatum, dorsolateral prefrontal cortex [DLPFC]). We also evaluated pain-specific and domain-general neural patterns and tested whether cue effects on intensity and valence were mediated through similar or distinct mechanisms.

## Results

### Behavioral outcomes: Stimulus level, expectation, and cue effects

#### Effects of stimulus type and magnitude on subjective perception

We focus on the preregistered model that controlled for Group differences (Heat vs. Salt; Salt vs. Sugar); behavioral results for models that replaced Group with Modality (Heat vs. Taste) or Aversiveness (Appetitive vs. Aversive) are provided in Table S2 for completeness and are acknowledged when they are inconsistent with our main model.

On the Calibration visit, we observed high correlations between temperature/concentration and subjective intensity in all conditions (see Figure 1A and Supplemental Results). Linear mixed models revealed strong associations between stimulus temperature/concentration (z-scored) and subjective intensity across all conditions (see Figure 1A), such that an increase of one SD temperature or concentration was associated with 2.2 units higher subjective intensity (B = 2.20*, p* < .001). We also observed a main effect of Condition (*F*(2, 4728.6) = 255.5, *p* < .001) and a Stimulus Temperature/concentration x Condition interaction (*F*(2,4728) = 34.68, *p* < .001), such that intensity ratings and slopes were steepest with saline solution (Intercept = 5.61, B = 2.42; all *p*’s < .001), followed by heat (Intercept = 4.68, B = 2.28; all *p*’s < .001), and then sucrose (Intercept = 4.18, B = 1.88; all *p*’s < .001). Pairwise differences between conditions were all significant (all *p*’s < .001). For additional details on psychophysics of each modality, including associations with unpleasantness and pleasantness, see Supplement Results.

**Figure 1.**
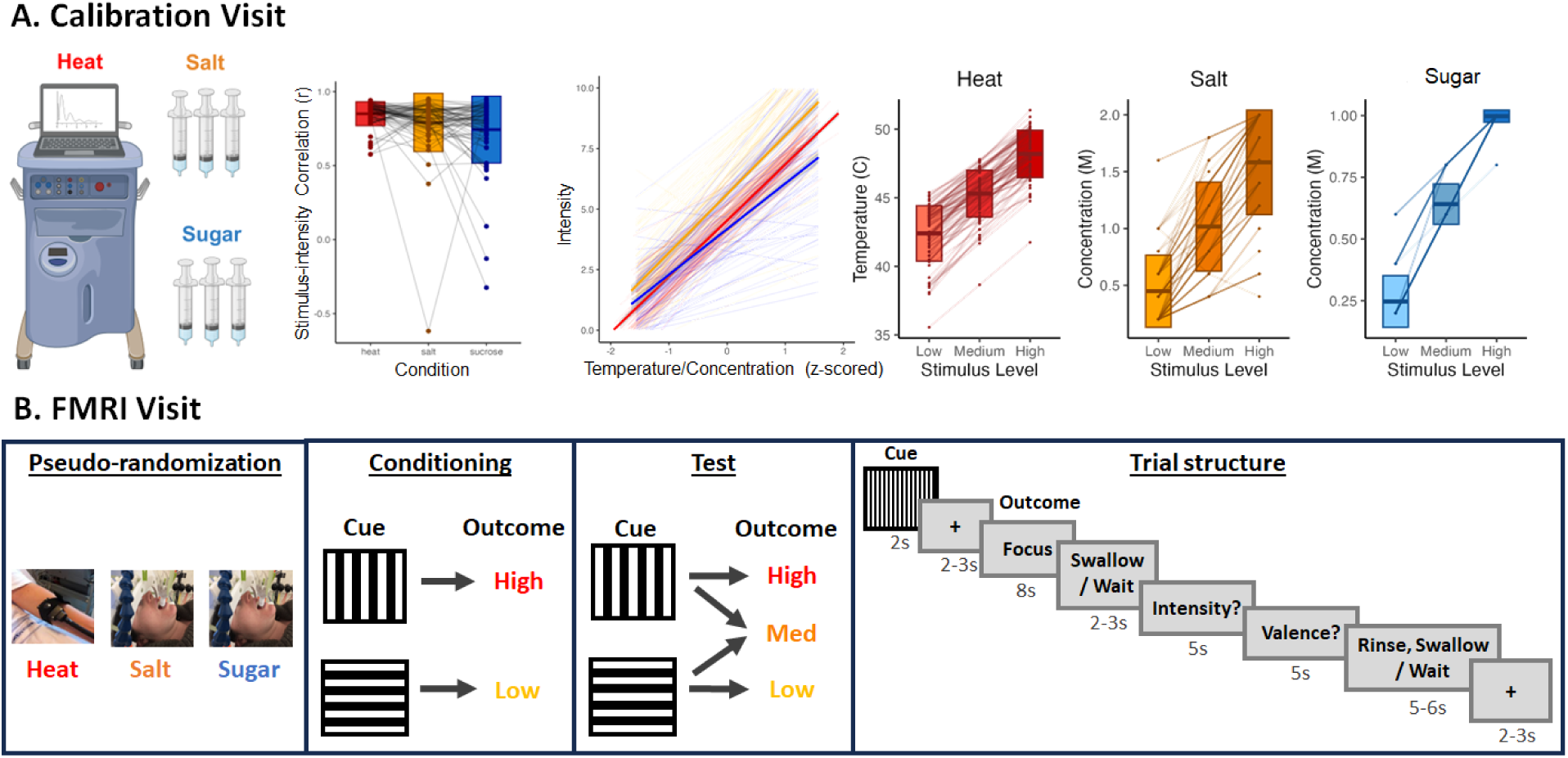
Calibration and fMRI Experiment Design. (A) Calibration design and results. During the Calibration Visit, all participants underwent thermal stimulation on their left volar forearms and self-administered salt and sucrose solutions at varying intensities and concentrations to evaluate psychophysical profiles for each modality. Stimulus magnitudes were correlated with subjective intensity in each modality (second-left panel). Linear mixed models (middle panel) revealed strong effects of stimulus magnitude (temperature/concentration) on subjective intensity across all modalities (B = 2.20, *p* <.001), with steepest slopes in the Salt Group. For each participant and each modality, temperatures or concentrations were selected corresponding to low, medium, or high intensity for use during the fMRI session (three right panels). (B) fMRI experiment design. After the Calibration Visit, participants were pseudo-randomly assigned to one of three groups: Heat, Salt, or Sugar (see Methods). They returned for an fMRI session, where they underwent a second calibration followed by the fMRI experiment. fMRI scanning began with a conditioning phase, during which High Cues were always followed by a stimulus calibrated to elicit ratings of high intensity, and Low Cues were followed by a stimulus calibrated to elicit ratings of low intensity. Following conditioning, High and Low Cues each had a 50% chance of being followed by a high or low stimulation and a 50% chance of being followed by a stimulus calibrated to elicit moderate intensity. After 8 seconds, participants in the Salt and Sugar groups were prompted to swallow the solution. Participants reported intensity and valence ratings on each trial. Taste group participants rinsed with a neutral solution between each trial to prevent carry over effects. Heat participants were prompted to wait during the swallow and rinse phases.

We next used linear mixed models to examine associations between changes in stimulus magnitude (low, medium, or high temperature or concentration) and subjective perception on the fMRI visit, when subjects were assigned to Group. We observed significant main effects of Stimulus Level on both intensity and valence, such that participants reported greater subjective intensity and subjective unpleasantness as a function of outcome magnitude across all groups (see Figure 2A; Intensity: B = 1.940, *p* < .001; Valence: B = 1.662, *p* < .001).

**Figure 2.**
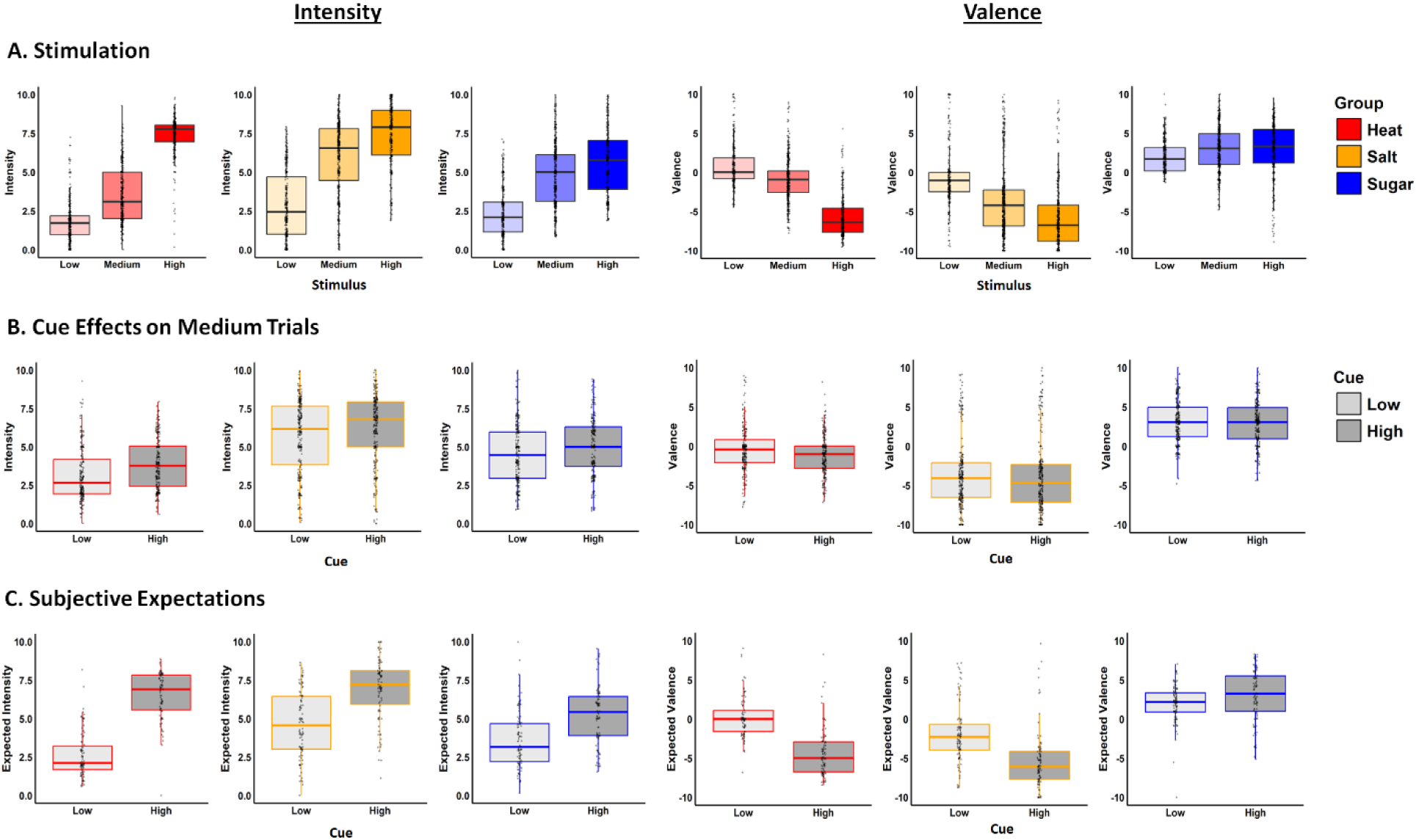
Behavioral Results during the fMRI Visit. (A) Effects of changes in stimulation on subjective intensity (left) and valence (right) for Heat (red), Salt (orange), and Sugar (blue) groups. Intensity ratings increased with stimulus intensity across all groups (main effect: B = 1.940, *p <* .001), while valence ratings became more negative (unpleasant) as stimulus intensity increased for Heat and Salt (Stimulus level * Group _Heat vs. Salt_: B = 0.355, *p* = .03), and more positive (pleasant) for Sugar (Stimulus level * Group _Salt vs. Sugar_: B = 0.575, *p* < .001, see Table S2). (B) Cue effects on subjective intensity (left) and valence (right). Across groups, participants reported higher intensity and less pleasantness in response to High Cues (dark) than Low Cues (light) (intensity: B = 0.305, *p* < .001; valence: B = -0.201, *p* = .0*04, see Table S3). There were no interactions with Group*. (C) Cue effects on subjective expectations. Across all groups, participants anticipated higher intensity (B = 1.312, *p* < .001) and higher unpleasantness (B = -1.187, *p* < .001) following High Cues compared to Low Cues. We observed significant interaction between Group and Cue for both intensity and valence expectations, driven by larger cue-induced difference for Heat relative to Salt and for Salt relative to Sugar (See Table S4).

We observed significant main effects of Group in both models, driven by higher intensity ratings in the Salt group than the Heat Group (B _Heat vs Salt_ = -1.043, *p* < .01; Heat: Mean = 4.161, SD = 2.643; Salt: Mean = 5.546, SD = 2.870) and by higher valence ratings in the Sugar group than the Salt Group in valence (B _Salt vs Sugar_ = -3.413, *p*< .001, Salt: Mean = -3.439, SD = 4.646; Sugar: Mean *=* 2.698, SD = 2.855). Consistent with these group differences, our additional models (see Table S2) confirmed that intensity ratings were impacted by Modality (B _Pain vs Tastes_ = -0.785, *p* < .001) but not Aversiveness (*p* > .50), while valence ratings were impacted by Aversiveness (B _Aversive vs Appetitive_ = -2.591, *p*< .001), but not Modality (*p* > .50). For complete results, see Table S2.

Main effects of Group and Stimulus Level were qualified by a significant Group x Stimulus Level interaction in all models (all *p’s* < .002; see Table S2 -1). There were stronger effects of Stimulus Level on intensity in the Salt group than the Heat group (B _Heat vs Salt_ = -1. 043, *p* = .0016), and there were positive effects of Stimulus Level on valence in the Sugar Group and negative effects in the Pain and Salt groups (B _Salt vs Sugar_ = -3.413, *p*< .001).

Finally, all models revealed a significant main effect of Cue, such that stimuli preceded by high cues were perceived as more intense (High: Mean = 6.169, SD = 2.283; Low: Mean = 3.332, SD *=* 2.312; all *p’s* < .001) and more unpleasant (High: Mean = -1.987, SD = 5.191; Low: Mean = 0.232, SD *=* 3.827; all *p’s* < .001) than stimuli preceded by low cues. We did not observe interactions between Group and Cue, or between Group, Cue, and Trial in any model (see Table S2). Importantly, we did observe significant Group x Stimulus x Cue interactions in the intensity model (B _Heat vs Salt_ = 1.055, *p* < .001; B _Salt vs Sugar_ = 0.507, *p* < .01) and a Stimulus x Cue interaction in valence models (B *=* -0.748, *p* < .01). We therefore conducted follow up models restricted to medium trials, which were equally crossed with cues. See all results in Table S2.

#### Learned cues modulate subjective perception in response to medium stimulation

We observed a significant main effect of Cue on Intensity (B = 0.301, *p* <.001) and Valence (B = -0.201, *p* = 0.004), such that medium stimuli preceded by high cues were perceived as more intense and more unpleasant than those preceded by low cues (Mean Intensity: High = 5.184, Low = 4.564, *p* < .001; Mean Valence: High = -0.786, Low = -0.435, all *p’s* < .01; see Figure 2B). There were no interactions between Group and Cue or between Group, Cue, and Trial in any model (none of those interactions survived model comparison and they remained non-significant even when included in the models), indicating that learned cues modulated perception similarly across domains.

Consistent with analyses across all trials, we observed significant main effects of Group on intensity ratings (B _Heat vs Salt_ = -1.146, *p* < .001) and valence ratings (B _Salt vs Sugar_ = -3.461, *p* < .001) in response to medium stimulation, driven by higher intensity in the Salt group than the Heat group (Heat: Mean = 3.581, SD = 1.783; Salt: Mean = 5.962, SD = 2.359; ; see Table S3-1) and higher valence ratings in the Salt group than the Sugar group (Salt: Mean = -3.630, SD = 4.608; Sugar: Mean= 2.956, SD = 2.783; see Table S3-1). There were no differences in intensity between Salt and Sugar groups or in valence between Pain and Salt (all *p’s* > 0.40).

Finally, we observed a main effect of Trial on valence ratings (B = -0.036, *p* < .001), such that stimuli were perceived as more unpleasant over time. See complete results in Table S2-1.

Results were similar when we modeled main effects of Aversiveness and Modality (see Table S3-2 and S3-3). Thus, our finding of main effects of Cue on valence and intensity ratings regardless of Group, provide behavioral evidence for a domain-general effect of predictive cues on subjective perception across modalities and aversiveness.

#### Learned cues induce conscious expectations

We also asked whether learned cues influenced participants’ explicit expectations. When we modeled expected intensity, we observed a significant main effect of Group, driven by higher intensity ratings in the Salt group than the Sugar Group (B _Salt vs Sugar_ = 0.516, *p* = .018; Salt: Mean = 5.820, SD = 2.296; Sugar: Mean = 4.425, SD = 2.029). We also observed a main effect of Cue (B = 1.312, *p* < .001), such that participants anticipated a higher intensity in response to High Cues (Mean = 6.253, SD = 1.972) relative to Low Cues (Mean = 3.667, SD = 2.024). This effect was qualified by a significant interaction of Group and Cue, driven by stronger cue-induced differences in expected intensity in the Heat Group relative to the Salt (B _Heat vs Salt_ = 0.612, *p* < .0018) and in the Salt group relative to the Sugar Groups (B _Salt vs Sugar_ = 0.499, *p* < .001). See Figure 2C.

We observed similar effects when we examined expected valence (Figure 2C). There was a main effect of Group, driven by higher expected unpleasantness in the Salt Group compared to the Heat Group (B _Heat vs Salt_ = -1.021, p = .019; Salt: Mean = -2.016, SD = 23.781; Sugar: Mean = -3.640, SD = 4.145) and greater pleasantness in the Sugar Group compared to the Salt Group (B _Salt *vs* Sugar_ = -3.581, *p* < .001; Sugar: Mean *=* 2.492, *SD* = 2.943). We also observed a main effect of Cue (B = -1.187, *p* < .001), indicating that participants expected overall greater unpleasantness with the High Cue (Mean = -2.215, SD = 5.102) than with the Low Cue (Mean = 0.105, SD = 3.441), and a significant Group x Cue interaction, characterized by larger cue-induced differences in expected valence when comparing the Heat Group to the Salt Group (B *_Heat vs_* _Salt_ = -1.157, *p* < .001) and the Salt Group to the Sugar group (B _Salt vs Sugar_ = -1.722, *p* < .001). See complete results in Table S4-1.

Results for models that measure the impact of Modality or Aversiveness rather than Group were consistent with our main model (Tables S4-2 and Table S4-3). These results underscore the significant role of predictive cues in modulating expectations of both intensity and valence, even without verbal instructions. Together, the behavioral results indicate that associations between predictive cues and the stimuli we administered led to changes in conscious expectations and perception across domains.

### Neural activation varies as a function of outcome magnitude and modality

#### Univariate and multivariate analyses of high vs. low stimuli

We first examined how variations in stimulus level affected brain responses during outcome delivery separately for each group. We examined results using univariate whole brain comparisons, multivoxel pattern analysis, and applying validated neural signature patterns. Results in the Heat Group survived small volumes correction (SVC) within *a priori* brain regions associated with pain and placebo analgesia (see Table 1), while results within the Salt and Sugar groups did not survive correction within these networks. We therefore report additional results from whole brain exploratory searches in Supplemental Table 5.

**Table 1.**
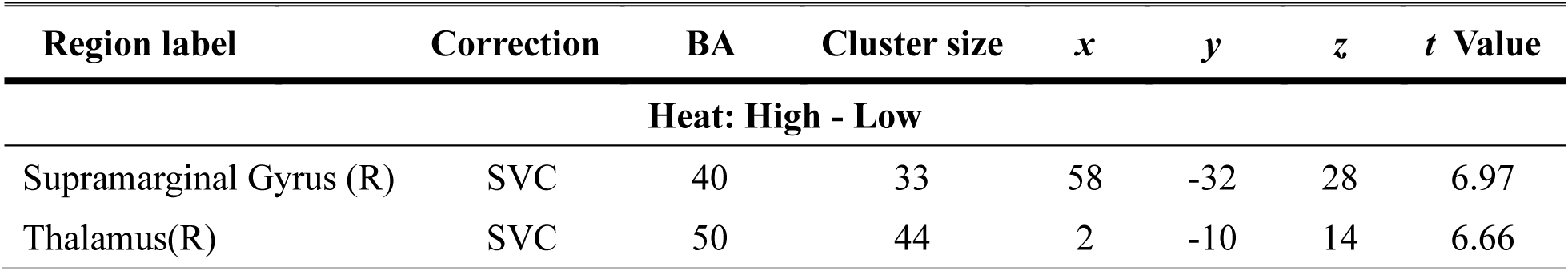
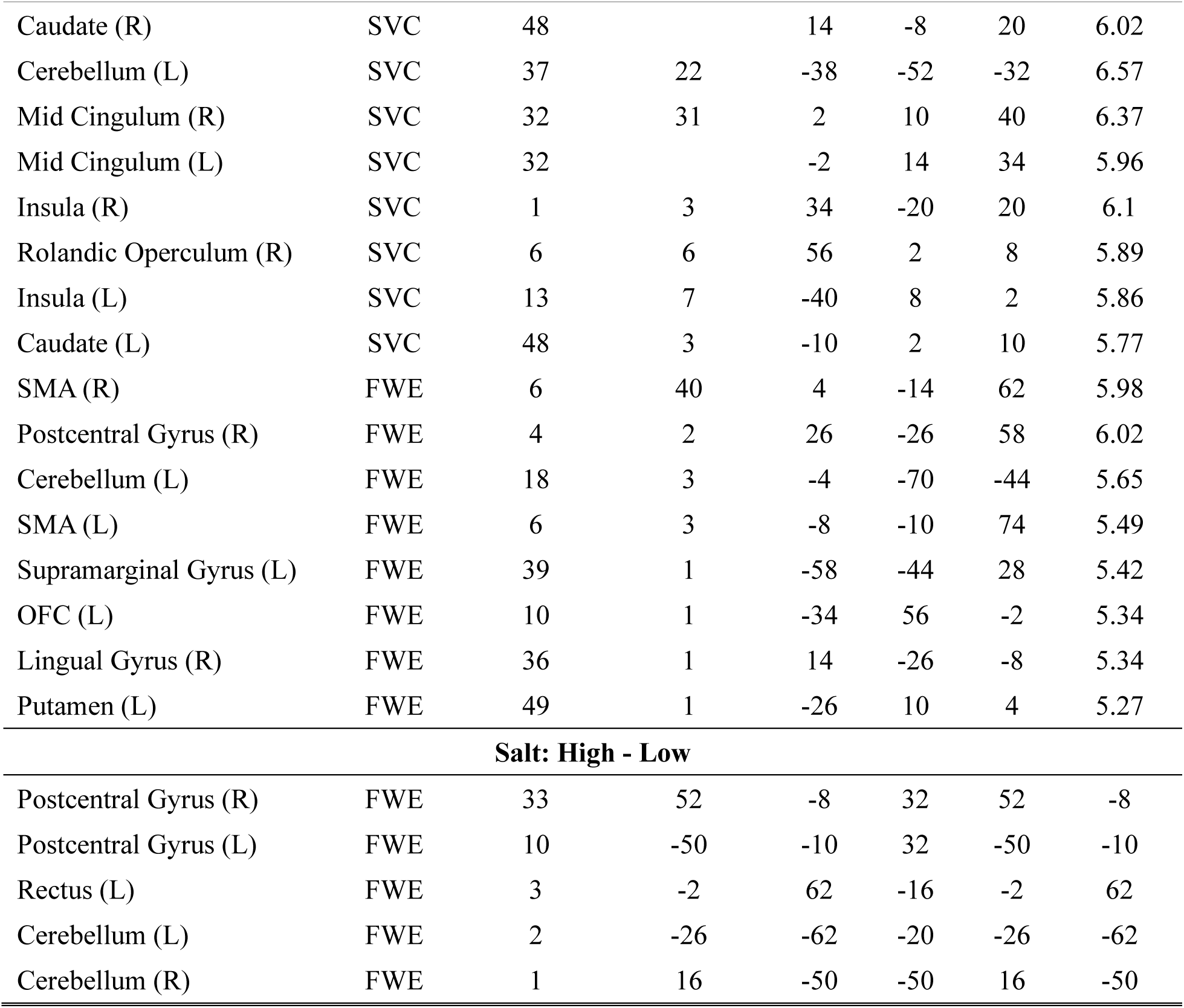
Results of mass-univariate functional analyses regarding stimulation level for Heat, Salt, and Sugar in the MNI space. Results are thresholded with Small Volume Correction (SVC; *p* < 0.05, FWE correction) in the pain-placebo analgesia mask (Atlas & Wager, 2014), or only cluster-wise FWE correction in the whole brain (FWE). Only Heat Group survived SVC with the pain-placebo analgesia mask. Only Heat and Salt showed significant whole-brain after FWE correction. No corrected results are survived for sugar. BA = Brodmann area, L = left hemisphere, mid = middle, R = right hemisphere, SMA = Supplementary Motor Area, OFC = Orbital Frontal Cortex.

| Region label | Correction | BA | Cluster size | $x$ | $y$ | $z$ | $t$ Value |
| --- | --- | --- | --- | --- | --- | --- | --- |
| **Heat: High - Low** |  |  |  |  |  |  |  |
| Supramarginal Gyrus (R) | SVC | 40 | 33 | 58 | -32 | 28 | 6.97 |
| Thalamus(R) | SVC | 50 | 44 | 2 | -10 | 14 | 6.66 |
| Caudate (R) | SVC | 48 |  | 14 | -8 | 20 | 6.02 |
| Cerebellum (L) | SVC | 37 | 22 | -38 | -52 | -32 | 6.57 |
| Mid Cingulum (R) | SVC | 32 | 31 | 2 | 10 | 40 | 6.37 |
| Mid Cingulum (L) | SVC | 32 |  | -2 | 14 | 34 | 5.96 |
| Insula (R) | SVC | 1 | 3 | 34 | -20 | 20 | 6.1 |
| Rolandic Operculum (R) | SVC | 6 | 6 | 56 | 2 | 8 | 5.89 |
| Insula (L) | SVC | 13 | 7 | -40 | 8 | 2 | 5.86 |
| Caudate (L) | SVC | 48 | 3 | -10 | 2 | 10 | 5.77 |
| SMA (R) | FWE | 6 | 40 | 4 | -14 | 62 | 5.98 |
| Postcentral Gyrus (R) | FWE | 4 | 2 | 26 | -26 | 58 | 6.02 |
| Cerebellum (L) | FWE | 18 | 3 | -4 | -70 | -44 | 5.65 |
| SMA (L) | FWE | 6 | 3 | -8 | -10 | 74 | 5.49 |
| Supramarginal Gyrus (L) | FWE | 39 | 1 | -58 | -44 | 28 | 5.42 |
| OFC (L) | FWE | 10 | 1 | -34 | 56 | -2 | 5.34 |
| Lingual Gyrus (R) | FWE | 36 | 1 | 14 | -26 | -8 | 5.34 |
| Putamen (L) | FWE | 49 | 1 | -26 | 10 | 4 | 5.27 |
| **Salt: High - Low** |  |  |  |  |  |  |  |
| Postcentral Gyrus (R) | FWE | 33 | 52 | -8 | 32 | 52 | -8 |
| Postcentral Gyrus (L) | FWE | 10 | -50 | -10 | 32 | -50 | -10 |
| Rectus (L) | FWE | 3 | -2 | 62 | -16 | -2 | 62 |
| Cerebellum (L) | FWE | 2 | -26 | -62 | -20 | -26 | -62 |
| Cerebellum (R) | FWE | 1 | 16 | -50 | -50 | 16 | -50 |

Relative to low heat stimulation, high heat stimulation elicited increases in brain regions associated with pain and placebo analgesia including the anterior and the mid-cingulate cortex (ACC/MCC), the right supramarginal gyrus, the anterior insula (aIns), the right Rolandic operculum, the posterior insula, bilateral caudate nucleus, the supplementary motor area (SMA), the frontal gyrus, and the left cerebellum (FWE correction). Whole brain exploratory search (*p* <.001 with 10 continuous voxels; applied consistently across all univariate and MVPA analyses) additionally identified the secondary somatosensory gyrus (SII) and the frontal gyrus (Figure 3A and Table 1). Consistent with these findings, one sample t-test showed that changes in stimulus level (High > Low) were significantly associated with the expression of all three neural signature patterns (Mean: NPS = 0.987, SD = 0.572, SIIPS = 236.163, SD = 161.649, Negative Affect = 0.042, SD = 0.041; all *t’s* > 3.905, *p’s* < 0.005).

**Figure 3.**
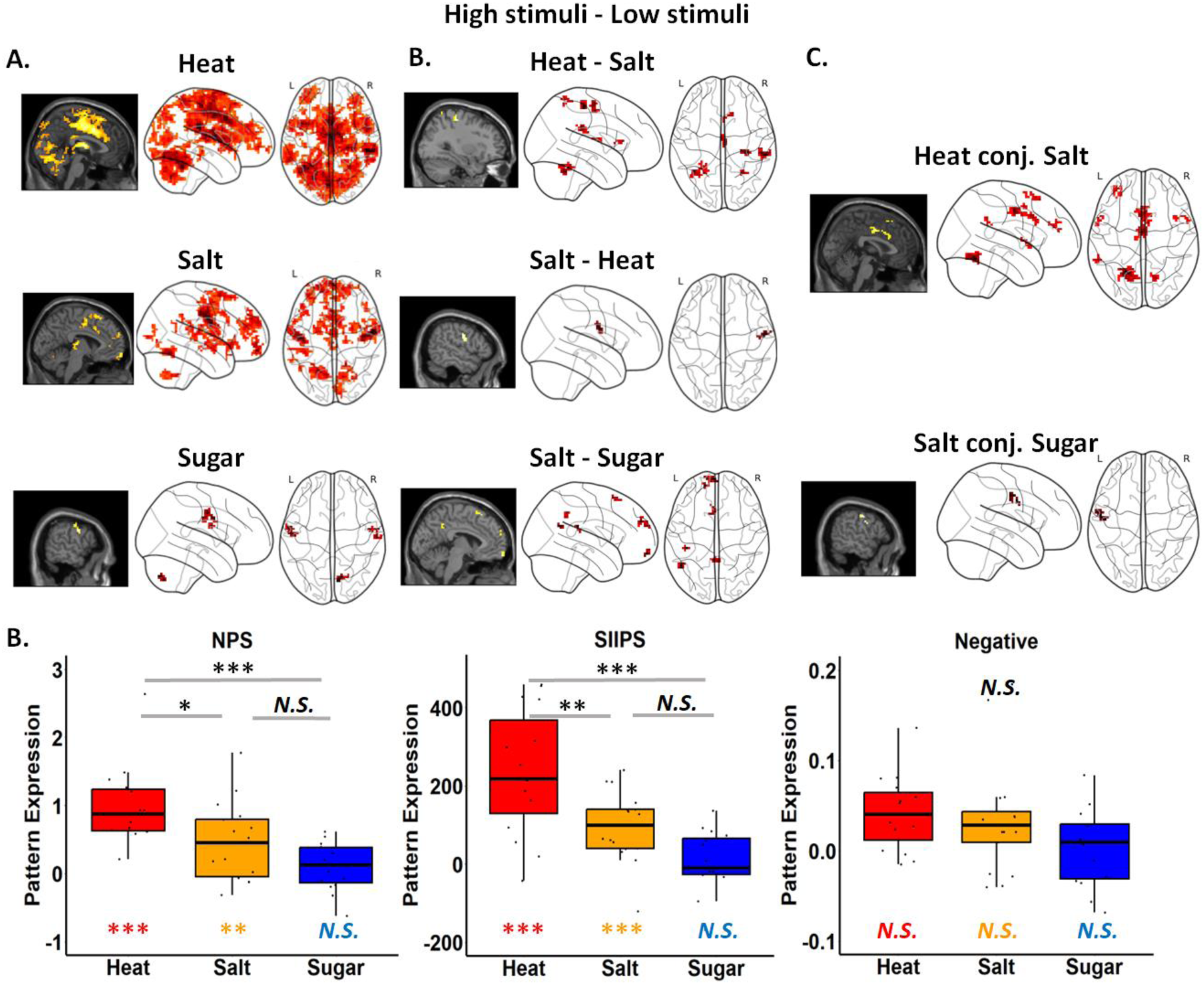
Brain responses to variations in stimulus level. (A) Neural activations in response to changes in stimuli (High > Low) for Heat, Salt, and Sugar groups. The Heat group exhibited the strongest and broadest activations, notably in the mid-cingulate cortex (ACC/MCC), anterior insula (aIns), posterior insula, and secondary somatosensory gyrus (SII). The Salt group showed key brain activations in the frontal gyrus, insula, thalamus, amygdala, and SII. Activations in both groups were largely consistent with previous studies. The Sugar group demonstrated activations in the inferior frontal gyrus, motor cortex, temporal cortex, and SII. (B) Group comparisons revealed distinct brain activations for Modality and Aversiveness. For Modality: 1) The contrast of Heat vs. Salt showed activations in the supramarginal gyrus, SII, precentral gyrus, frontal gyrus, MCC, parietal lobe, temporal lobe, and cerebellum. 2) The contrast of Salt vs. Heat showed significant activations in the postcentral gyrus. For Aversiveness: The comparison of Salt vs. Sugar revealed activations in the middle and posterior cingulate cortex, superior medial frontal gyrus, middle temporal gyrus, and SII. (C) Conjunction analyses showed shared activations for both modality and aversiveness. For modality, the Heat and Salt groups showed overlapping activations in the ACC/MCC, cerebellum, superior and middle frontal gyrus, temporal pole, right insula, and SII. For aversiveness, the Salt and Sugar groups showed common activations in the precentral gyrus, motor cortex, and SII. (D) Neural signature pattern analysis examined three brain patterns. For the two pain-related patterns, NPS (nociception-specific pattern) and SIIPS (non-nociception specific but pain-relevant pattern), both Heat and Salt groups showed significant expression above zero, with Heat exhibiting higher expressions compared to Salt and Sugar. For the Negative affect pattern, which is domain-general, both Heat and Salt groups showed significant expressions, but no group differences were observed. These results indicate domain-specific neural activations for Heat that cannot be entirely attributed to salience or aversiveness. Group differences are marked with black asterisks (significant) or “N.S.” (not significant). Within-group differences (one-sample t-test) are indicated in the color corresponding to the data bars in the plot. *** *p* < .001, ** *p* < .01, * *p* < .05.

For salt concentration (High > Low), whole brain exploratory analyses revealed significantly increased activation in areas involved in gustatory processing, including the superior and middle frontal gyrus, and SII (Figure 3A and Table 1). We further applied whole brain FWE correction and observed robust activations in SII, the middle frontal gyrus, and the cerebellum. Similar to the Heat Group, one sample t-test showed that changes in stimulus level ([High > Low]) were associated with significant expression of all three signature patterns (Mean _NPS_ = 0.451, SD _NPS_ = 0.564, Mean _SIIPS_ = 96.512, SD _SIIPS_ = 88.780, Mean _Negative Affect_ = 0.034, SD _Negative Affect_ = 0.062; all *t’s* > 2.312, *p’s* < 0.05).

Finally, we evaluated how changes in sucrose concentration (High > Low) impacted activation in the Sugar Group. In contrast to the Heat and Salt Groups, we did not observe any significant activations that survived FWE correction. Whole brain exploratory analyses revealed that high sucrose concentration was associated with increased activation in the precentral and postcentral gyrus, cerebellum, and SII based on univariate results (Figure 3A and Table 1). We did not observe any effect of changes in stimulus level on neural signature pattern expression (Mean _NPS_ = 0.102, SD _NPS_ = 0.351, Mean _SIIPS_ = 7.101, SD _SIIPS_ = 78.203, Mean _Negative Affect_ = 0.004, SD _Negative Affect_ = 0.042; all *t’s* < 1.131, *p’s* > 0.2).

We also conducted MVPA to identify which brain regions reliably discriminated changes in stimulus intensity in each group. MVPA results largely aligned with the univariate findings, that Heat survived the pain-placebo analgesia mask and activated broader and strongest activations around the whole brain compared to Salt and Sugar (see MVPA results in Figure S1 and Table S6). The consistency between the univariate and MVPA results further confirmed the robustness of magnitude-related brain activations across domains.

#### Comparisons between groups using neural signature patterns and univariate results

Given the differences across modalities in behavioral ratings observed for high vs. low stimuli, we further evaluated comparisons between groups. Comparisons between Heat and Salt Groups isolate effects of Modality on responses to aversive stimuli. Based on the whole brain search, changes in stimulus intensity were associated with larger activations in the Heat Group compared to the Salt Group in regions including SII, cerebellum, thalamus, and dorsal striatum/caudate nucleus, the supramarginal gyrus, the precentral gyrus, the middle and superior frontal gyrus, MCC, the parietal lobe, and the temporal lobe (Figure 3B and Table S5); the only region that displayed stronger stimulus-related activations in the Salt Group than the Heat Group was the postcentral gyrus.

We next compared stimulus-related increases between Salt and Sugar Groups to isolate effects of Aversiveness on responses to taste stimuli. Relative to the Sugar Group, the Salt Group showed stronger stimulus-related activations in the middle and posterior cingulate cortex, the superior medial frontal gyrus, the middle temporal gyrus, and SII based on whole brain search (Figure 3B and Table S5). No area showed larger increases in the Sugar Group than in the Salt Group.

Finally, we used conjunction analyses to ask whether any effects of stimulus outcome were shared across modalities (Figure 3C and Table S5). Whole brain search analyses revealed shared activations for Heat and Salt within the pain-placebo analgesia mask in SMA, motor cortex, ACC, aIns, and cerebellum, the superior and middle frontal gyrus, the temporal pole, the right insula, and SII, though none of these activations survived SVC. Common regions between the Salt and Sugar Groups displayed shared increases in the precentral gyrus, the motor cortex, and SII. Finally, only one cluster in the left cerebellum exhibited a conjunction across all three groups (k = 2).

Finally, we used one-way ANOVAs to assess group differences in stimulus level effects on neural expression [High > Low] for each neural signature pattern (see Figure 3D). For NPS and SIIPS, we observed a significant Group effect (NPS: F = 11.531, *p* < .001, SIIPS: F = 15.251, *p* < .001), with post-hoc comparisons revealing a stronger expression for Heat compared to Salt (NPS: *p* = 0.014, SIIPs: *p* = 0.003) and Sugar (NPS: *p* < .001, SIIPs: *p* < .001). No significant differences were found between Salt and Sugar (*p’s* > 0.09) for either NPS or SIIPS. For the negative affect pattern, no significant group effect emerged (F = 2.458, *p* = 0.097). See results of neural signature pattern comparison in Figure 3D.

### Common effects of learned cues on orbitofrontal cortex activation during anticipation across modalities

Given that changes in stimulus level strongly shaped perception and brain responses during stimulus outcome, and cues paired with these outcomes were associated with differences in expectations, we next tested how cues influenced brain activation prior to outcome delivery, i.e., during anticipation. Whole-brain exploratory analyses revealed activation in the bilateral OFC (Figure 4), with no additional activations observed within, across, or between groups.

**Figure 4.**
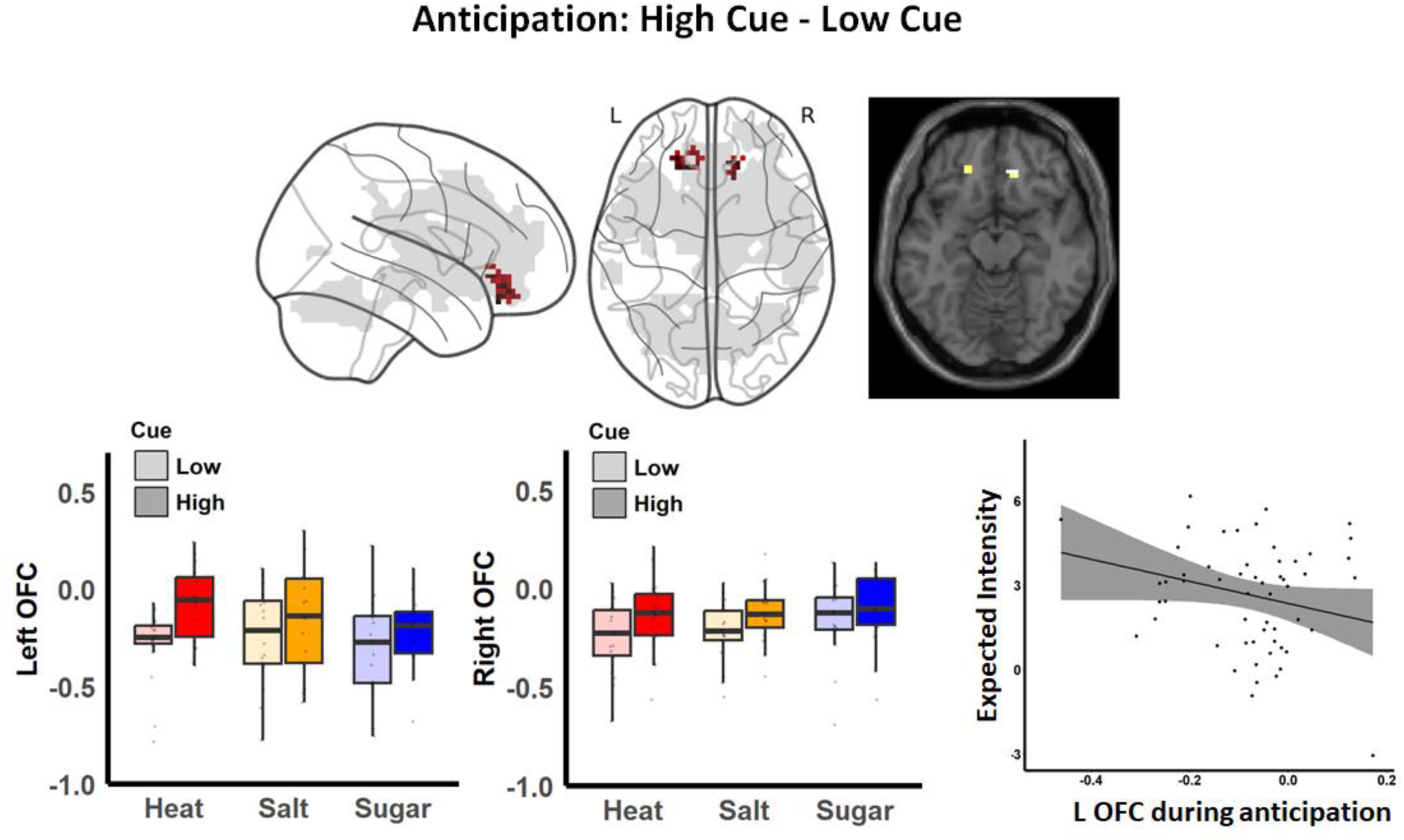
Brain responses to cues during anticipation. Significant activations in the bilateral orbitofrontal cortices (OFC) were observed during anticipation across all groups. These activations (shown in red) overlapped with our previous pain-placebo analgesia mask (shown in gray; Atlas & Wager, 2014), though they did not survive small volume correction. The absence of group differences suggested the domain-generality of these activations. Pearson correlation analysis indicated a trend towards a correlation between cue-related activations in the left OFC and intensity ratings across groups (r = -0.25, *p* = 0.063). This evidence further supports the relevance of OFC activation to anticipation, particularly in relation to expected intensity.

Consistent with our preregistration, we evaluated whether OFC anticipatory activation was correlated with participants’ expected intensity and/or valence. We extracted averaged ROI activations separately for the left and right OFC across all blocks. Spearman correlation analysis revealed a trend toward correlation between cue effects on the left OFC during anticipation and cue effects on expected intensity (r = -0.25, *p* = 0.063). No significant correlations were detected for the right OFC or for expected valence ratings (all abs(r) < 0.12, *p* > 0.3; Figure 4). These findings demonstrate the domain-general engagement of the OFC based on predictive cues during the anticipation of pleasant and unpleasant stimuli, regardless of modality.

### Mediation of common cue effects on subjective intensity and valence

#### Intensity Mediation

Given that cues affected anticipatory activation and subjective ratings similarly across groups, we next used mediation analysis to isolate brain mediators of dynamic cue effects on subjective perception on medium trials regardless of group. We first searched for mediators of dynamic cue effects on subjective intensity (see Figure 5A, Table 2, and Supplement Figure 2A). Path *a* in our intensity mediation model isolates brain regions whose response to medium stimuli varies as a function of cue [High vs Low]. Path *a* effects were not associated with significant expression of any signature pattern. Positive Path *a* effects [High > Low] were observed in the left dorsomedial prefrontal cortex (DMPFC) based on whole-brain FDR correction (see Figure 5A and Table 2). Whole brain exploratory analyses revealed positive Path *a* effects in the right medial OFC, left inferior frontal gyrus (IFG), left DMPFC, and MPFC / rostral ACC, and negative path *a* effects [Low > High] in the left superior temporal gyrus, the right IFG, the right angular gyrus, the left postcentral gyrus, the left inferior parietal lobule (IPL), and the right DLPFC, among other regions (see Table S7-1 and Figure S2A).

**Figure 5.**
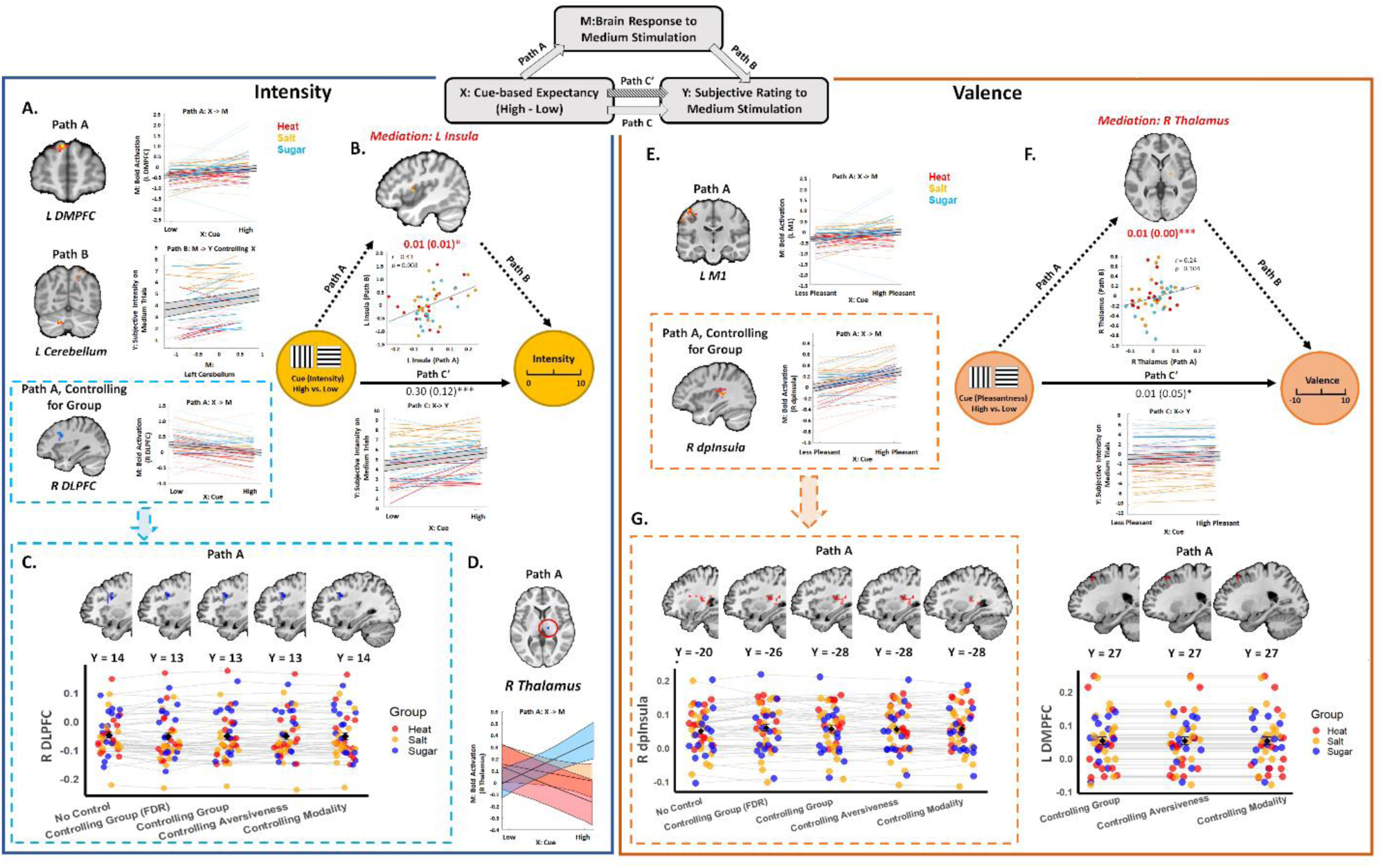
Mediators of cue effects on subjective perception across modalities. This figure presents results of voxelwise multilevel mediation analyses that searched for brain mediators of cue effects on subjective experience (see schematic diagram at the top center). The left panel depicts significant path effects for Intensity as the dependent measure (Y), and the right panel illustrates corresponding effects for Valence (Pleasantness). Slope plots below each path diagram illustrate behavioral results (i.e., Path C). Complete voxelwise results are reported in Supplementary Tables 5. (A) Path A and Path B effects for Intensity (FDR-corrected). Individual slopes representing each effect are shown on the right, with the black line indicating the average slope across all participants. We identified a positive Path A effect (High Cue > Low Cue) in the left DMPFC, driven by higher DMPFC activation with high cues across all groups. and a positive Path B effect in the cerebellum, driven by positive associations between activation and subjective intensity, when controlling for Cue. After controlling for Group, a negative Path A effect emerged in the right DLPFC, driven by higher DLPFC activation on medium trials preceded by low cues relative to high cues. (B) Mediators of cue effects on intensity ratings. Whole brain uncorrected results indicated that the left insula positively mediated cue effects on intensity ratings (Path ab = 0.01, STE = 0.01, *p* < 0.05). Mediation was driven by the covariance between Paths a and Paths b, such that individuals who had stronger cue effects on insula activation on medium trials also had stronger associations between insula and subjective intensity, controlling for cue (see scatterplot). (C) Comparison of Intensity path effects with and without moderators. The Path A effect in the right DLPFC remained consistent and robust irrespective of controlling for Group, Aversiveness, and Modality (whole-brain, uncorrected). (D) FDR corrected moderation results. In the right thalamus, the Sugar group exhibited a significantly more positive Path A effect (increased activation on medium trials preceded by High cues) than the Salt and Heat groups, whereas the Heat group demonstrated a significantly more negative effect (larger deactivation on medium trials preceded by High cues) compared to the Salt group. (E) Path A and Path B effects for Valence. All results passed FDR correction. A positive Path A effect (cue: high < low for Heat and Salt; high > low for Sugar) was observed in the left Premotor Cortex (M1), and after controlling for Group, a positive Path A effect was found in the right dorsal posterior insula (dpInsula). Both regions showed higher activation on medium trials preceded by cues that predicted less aversive outcomes. Path B effects did not survive FDR-correction; see Tables S5 for complete results. (F) Mediators of cue effects on subjective valence. FDR-corrected results indicated the right thalamus positively mediated cue effects on pleasantness ratings (Path ab = 0.01, STE < 0.001, p < 0.001). Mediation was driven by covariance, such that individuals who had stronger cue effects on thalamus activation on medium trials also had stronger associations between thalamus and subjective intensity, controlling for cue, although correlations at the cluster level were not significant (see scatterplot). (G) Comparison of Valence path effects with and without moderators. The Path A effect in the right dorsal posterior insula and left DLPFC demonstrated similar activations regardless of moderation or control for Group, Aversiveness, and Modality. Positive activations are displayed in warm colors, and negative activations are in cool colors. Dots and lines represent groups: Heat (red), Salt (yellow), and Sugar (blue). Values outside parentheses denote estimated mediation path coefficients, while values inside parentheses indicate standard errors. Significance levels: * *p* < 0.05, ** *p* < 0.01, *** *p* < 0.001

**Table 2.** Single-trial mediation results on subjective intensity and valence. Significant activations were presented for each path. All results passed False Discovery Rate (FDR) corrections. DMPFC: Dorsomedial Prefrontal Cortex; SPL: Superior Parietal Lobule. See uncorrected results in Supplementary Table 7.

| INTENSITY |  |  |  |  |  |  |  |
| --- | --- | --- | --- | --- | --- | --- | --- |
| Path *a* |  | Peak |  |  | Voxel Size | Z | P |
|  |  | *x* | *y* | *z* |  |  |  |
|  | **High > Low**<br>DMPFC (L) | -10 | 58 | 38 | 6 | 3.55 | 0.0003 |
| Path *b* | **Positive Association** |  |  |  |  |  |  |
|  | Temporal Pole (L) | -44 | -2 | -46 | 1 | 4.44 | <.0001 |
|  | Cerebellum (L) | -16 | -68 | -34 | 7 | 4.41 | <.0001 |
|  | Pons | 4 | -14 | -34 | 1 | 4.59 | <.0001 |
|  | Midbrain | -2 | -44 | -28 | 15 | 4.32 | <.0001 |
|  | Superior Occipital Gyrus (L) | 16 | -74 | 52 | 21 | 4.79 | <.0001 |
|  | Superior Occipital Gyrus (R) | -10 | -74 | 52 | 7 | 6.1 | <.0001 |
| VALENCE |  |  |  |  |  |  |  |
| Path *a* |  | Peak |  |  | Voxel Size | Z | P |
|  |  | *x* | *y* | *z* |  |  |  |
|  | **High > Low** |  |  |  |  |  |  |
|  | M1 (L) | -46 | -20 | 56 | 46 | 4.7 | <.0001 |
|  | SPL (R) | -10 | -46 | 64 | 6 | 5.53 | <.0001 |
|  | DMPFC (L) | -14 | -2 | 70 | 4 | 4.81 | <.0001 |
| Path *ab* | **Positive Mediation**<br>Thalamus (R) | 20 | -14 | 2 | 2 | 4.47 | <.0001 |

Path *b* evaluates the relationship between brain activation and subjective intensity controlling for cue. We observed significant Path *b* effects on expression of the NPS (*t*(1,46) = 2.15, *p* = .037; Mean = 0.19, SD = 0.60) and marginal effects on expression of the SIIPs (*t*(1,46) = 1.96, *p* = .056; Mean = 65.7, SD = 230.08) and the negative affect pattern (*t*(1, 46) = 2.01, *p* = 0.05, Mean = 0.02, SD = 0.07). Whole brain FDR correction revealed positive Path *b* effects in the temporal pole, left cerebellum, midbrain, and bilateral superior occipital cortex (see Figure 5A and Table 2). Uncorrected voxel-wise results are included in Figure S2A, Table S7-1.

Whole brain correction did not reveal any significant mediators, nor were mediation effects associated with significant expression of any signature pattern. However, uncorrected results revealed positive mediation by the left anterior/middle insula (Figure 5B and Table S7-1), in the same region that was previously found to mediate cue effects on subjective pain (Atlas et al., 2010). Extracting from this region revealed that it was driven by covariance between Paths *a* and *b*, such that individuals who showed stronger cue effects on left insula activation (Path *a*) also showed stronger associations between left insula activation and subjective intensity (Path *b*), as visualized in the scatterplot in Figure 5B. We also observed negative mediation by the left visual cortex, driven by negative covariance between Paths *a* and *b*, such that individuals who showed stronger cue effects on visual cortex had more negative associations between activation and subjective intensity, consistent with suppression effects (Figure S2A Table S7-1).

#### Valence mediation

Our second mediation model searched for mediators of cue effects on subjective pleasantness, while taking group differences in aversiveness into account (see Methods). Path *a* identified brain regions that showed greater stimulus-evoked activation on medium trials in response to cues that predicted less aversive / more appetitive outcomes, regardless of Group. We observed marginal Path *a* effects on expression of the NPS (*t*(1, 45) = 1.87, *p* = .068; Mean = 0.05, SD = 0.19) and the SIIPS (*t*(1,45) = -1.78, *p* = .08; Mean = -15.93, SD = 60.56), but no impact on the negative affect pattern (*p* > 0.7). Whole brain FDR correction revealed positive Path *a* effects in left motor cortex (Figure 5E), DMPFC, and left superior parietal lobule (see Table 2). Exploratory uncorrected analyses additionally revealed positive Path *a* effects in the OFC, amygdala, insula, and DLPFC (see Figure S2B and Table S7-6). No voxels displayed negative Path *a* effects (i.e., greater activation for cues that predicted the more aversive outcome).

Path *b* identifies regions that predict increases in pleasantness on medium trials, while controlling for cue, regardless of group. Path *b* effects were inversely related to expression of the NPS (*t*(1, 45) = -2.62, *p* = 0.01; Mean = -0.47, SD = 1.22) and marginally related to expression of the negative affect pattern (*t*(1, 45) = -1.88, *p* = 0.066; Mean = -0.03, SD = 0.12). There were no associations with SIIPs expression (*p* > 0.5) and no effects survived FDR correction. Uncorrected whole brain exploratory analyses revealed positive Path *b* effects in the cerebellum, the right hippocampus/amygdala, brainstem, the left Rolandic operculum, the right SMA, and the left visual cortex (see Table S7-6). There were no regions with negative Path *b* effects.

Cue effects on valence were positively mediated by the right thalamus, based on whole brain FDR correction (see Figure 5F and Table 2). Extracting from this region revealed that effects were driven by positive covariance between paths, although correlations were not statistically significant (Figure 5F). Uncorrected analyses additionally revealed mediation by left VLPFC, bilateral DMPFC, right DLPFC, and midbrain in an area near the ventral tegmental area (see Figure S8-6). There were no associations between mediation effects and pattern expression (all *p*’s > 0.2).

### Moderated mediation: Evaluating group differences in cue effects on subjective intensity and valence

#### Moderation of intensity effects by group, modality, and aversiveness

We used moderated mediation to determine whether cue effects on medium trials varied across Groups (Pain vs. Salt, and Salt vs. Sugar), Modality (Heat vs. Tastes), or Aversiveness (Aversive vs. Appetitive), and to evaluate results when controlling for Group/Modality/Aversiveness. When controlling for Group, we observed negative Path *a* effects in the right DLPFC based on FDR-correction (see Figure 5A and Supplementary Table S7 (2-3)); the same region was observed in uncorrected whole brain exploratory analyses when controlling for Group, Aversiveness or Modality (see Figure 5C and Supplementary Table 7 (2-5)) or omitting moderators (see Figure 5 and Table S7-1). We observed moderation of Path *a* effects in right thalamus by Group, such that participants in the Sugar Group showed greater taste-induced thalamus activation following High cues relative to Low cues, but participants in the Salt and Heat groups showed deactivation with High cues relative to Low cues, and negative Path *a* effects were stronger in the Heat Group than the Salt Group (see Figure 6A). There were no other mediation or moderation effects that survived FDR-correction.

**Figure 6.**
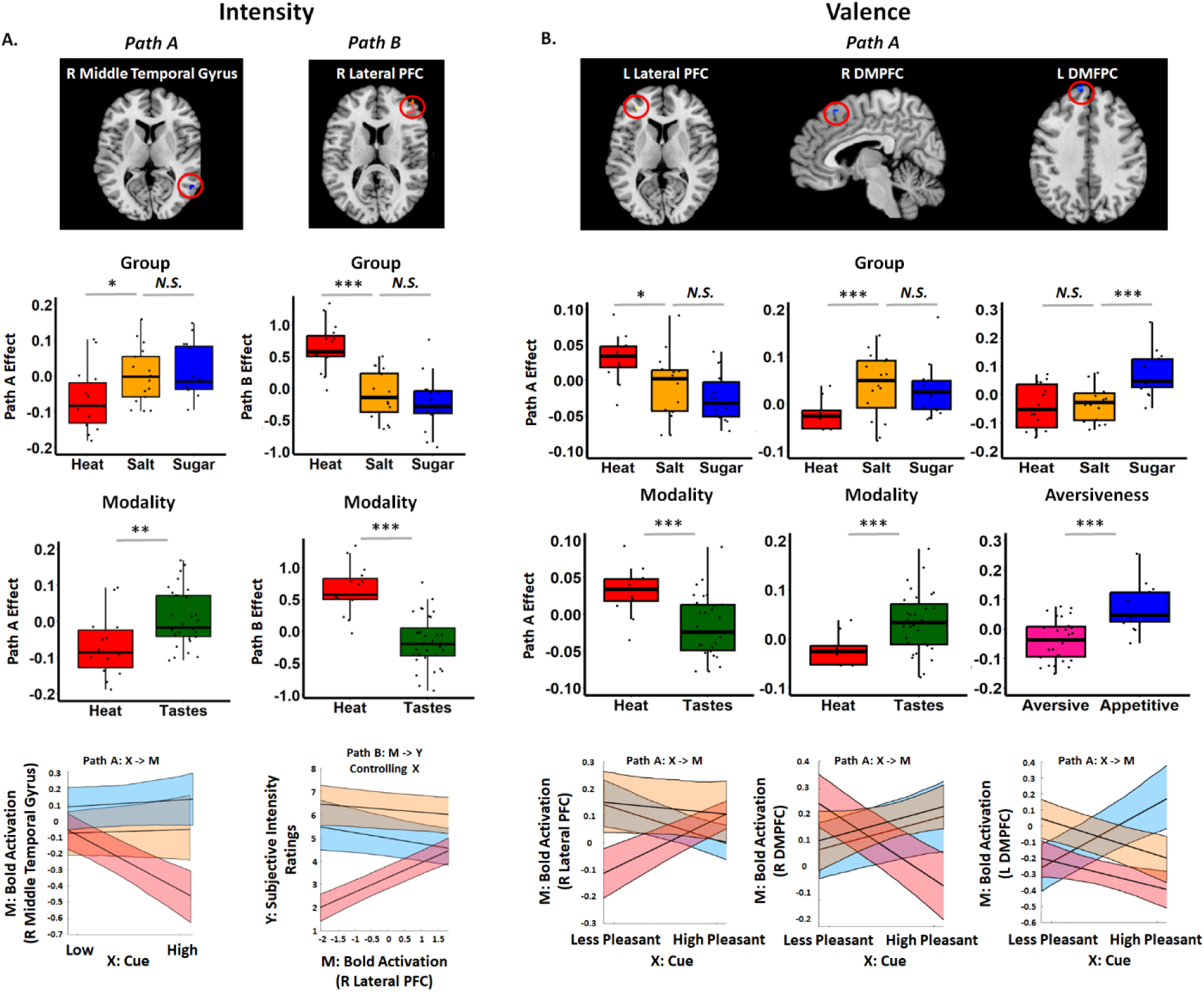
Moderated Mediation: Results by Group, Modality, and Aversiveness. We included moderators to measure whether mediation effects varied across groups and as a function of modality and/or aversiveness. Only moderation in the thalamus survived FDR-correction (see Figure 5D). All effects here are presented at an uncorrected voxelwise threshold of p < .001. (A) Intensity Model Moderators. We observed significant moderation effect of Group and Modality on Path A effects in the right temporal gyrus, primarily driven by a greater right temporal gyrus deactivation on medium trials following High cues in the Heat Group, but not compared to Salt and Sugar (Taste) Groups. Group and Modality also moderated Path B effects in the right lateral prefrontal cortex (PFC), driven by positive associations between lateral PFC and subjective intensity, controlling for cue, in the Heat Group but not the Salt and Sugar (Taste) Groups. (B) Valence Model Moderators. Group and Modality moderated Path A effects with positive moderation in the left lateral PFC, primarily driven by a higher positive lateral PFC activation with the less aversive cue in the Heat Group but not the compared to Salt or Sugar (Taste) Groups, and a negative moderation effect in the right dorsomedial prefrontal cortex (DMPFC), driven by deactivation with the less aversive cue in the Heat Group, and activation in the Salt and Sugar (Taste) Groups. We also observed moderation of Path A effects by Group and Aversiveness in the left DMPFC, driven by higher DMPFC activation with the more pleasant cue in the Sugar Group, but more deactivation with the less aversive cue in the in the Salt and Heat (Aversive) Groups. Cluster-wise slope estimates from mixed models are shown in the plot at the bottom right as a function of group. Positive activations are represented in warm colors, and negative activations in cool colors.

Exploratory uncorrected results indicated that Path *a* effects [High vs Low] differed as a function of Group in the right middle temporal gyrus (see Figure 6A and Table S7-2). Extracting from this region confirmed that there were stronger cue effects on medium stimulus activation in the Salt Group than the Heat Group (*t* _Heat vs Salt_ = 2.555, *p* = 0.016; Mean _Heat_ = 0.076, SD _Heat_ = 0.093, *p* = 0.007; Mean _Salt_ = 0, SD _Salt_ = 0.077, *p* > 0.9), Consistent with these findings, we observed moderation of Path a effects by Modality in the right middle temporal gyrus (Table S7-4). There were no regions that showed differential Path a effects as a function of Aversiveness. We also observed moderation of Path *b* effects in the cerebellum, right Supramarginal Gyrus, and right lateral PFC, with stronger associations between brain activation and subjective intensity in the Heat Group than the Salt Group, which were also observed when we included Modality as a moderator (see lateral PFC as an example in Figure 6, and the complete results in Supplementary Table 7 (2-4)). We did not observe moderated mediation (path ab) in any voxels for any of the group-level factors.

Finally, we used three-way ANOVAs to test whether path effects varied across groups. We did not observe group differences in pattern expression for Path *a* or Path *b* for any signature pattern (all *p*’s > 0.2). However, we observed marginal group differences in associations between mediation effects and expression of the SIIPS pattern (F(2, 44) = 2.660, *p* = .081), which effect was mainly driven by a more positive mediation in the Heat Group (Mean _Heat_ = 3.742, SD _Salt_ = 12.740) compared to the Salt Group (*t*(1, 30) =1.765, *p* = 0.099; Mean _Salt_ = -6.017, SD _Salt_ = 13.387).

#### Moderation of group effects for valence

When controlling for Group in our valence mediation model, whole-brain FDR correction revealed significant positive Path a effects in the right dorsal posterior insula and left superior parietal lobule (See Figure 5E and Supplementary Table 7 (7-8)). Exploratory uncorrected analyses additionally revealed Path *a* effects in the left DMPFC, right thalamus, and right middle insula (see Table S7-7). Results were similar when we controlled for Aversiveness and Modality (Figure 5G and Supplementary Table 7 (9-10)).

Path *a* effects varied as a function of Group in several regions based on uncorrected results; there were no differences that survived whole-brain FDR correction. When evaluating differences in Path *a* effects between Heat and Salt Groups, we observed positive moderation in the left lateral PFC and negative moderation in the right DMPFC. Extracting from these regions confirmed these differences, such that the Heat Group exhibited positive Path *a* effects [Low > High] in the left lateral PFC, whereas there were no cue effects on taste-evoked activation in the Salt Group (t _Heat vs Salt_ =2.51, *p* = 0.018; Mean _Heat_ = 0.41, SD _Heat_ = 0.040, *p* = 0.002; Mean _Salt_ = 0.006, SD _Salt_ = 0.061, *p* > 0.68). The two groups also showed differences in right DMPFC, such that the Heat Group exhibited greater heat-evoked DMPFC activation with High cues, whereas the Salt Group exhibited greater taste-evoked activation with Low cues (*t*(1, 30) _Heat vs Salt_ = -3.6209, *p* < .001; Mean _Heat_ = -0.062, SD _Heat_ = 0.062, *p* = 0.003; Mean _Salt_ = 0.03, SD _Salt_ = 0.076, *p* > 0. 12). Aligned with these activations, we found significant moderation by Modality in the left lateral PFC and the right DMPFC (Figure 6B and Supplementary Table 7 (7-10)). There were no differences between Salt and Sugar in either of the two clusters. When we tested for moderation by Salt vs. Sugar, we observed significant moderation in the left DMPFC. Extracting from this region revealed that both groups showed greater taste-evoked activation of left DMPFC in response to High cues relative to Low cues (t _Sugar vs Salt_ = 4.2462, *p* < .001; Mean _Sugar_ = 0.095, SD _Sugar_ = 0.109, *p* = 0.0047; Mean _Salt_ = - 0.045, SD _Salt_ = 0.075, *p* = 0.026), consistent with FDR-corrected Path A effects in our main intensity mediation model (Figure 5A). Consistent with these activations, we observed significant moderation by Aversiveness in the left DMPFC (Figure 6C and Table S7-10). There was no significant difference between Heat and Salt Groups in Path A effects in this cluster. Interestingly, we did not observe moderation of Path *b* or mediation effects by any level 2 factor (Group, Aversiveness, or Modality), nor did we observe significant mediation effects while controlling for level 2 factors. There were no group differences in expression of any signature pattern for any path (all p’s > 0.1).

## Discussion

Previous studies have shown that expectation can modulate first-hand experiences of pain and other aversive stimuli, particularly involving activations in regions such as the OFC and the aIns (Levy & Glimcher, 2012; Sharvit et al., 2019). As the majority of studies have used instructions or explicit cues to induce expectations (Atlas, 2023), only a few studies have examined effects of pure uninstructed learning on subjective perception (Colloca & Benedetti, 2009; Jensen et al., 2012; Atlas et al., 2022)and it remains unclear whether learned expectancy effects differ in their impact on perception across different modalities. In this study, we identified the effects of learned cues on both anticipation and perception, observed both at the behavioral and neural levels. By applying three neural signature patterns, we confirmed that neural responses to changes in hedonic stimuli include both domain-general (i.e., negative affect pattern) and domain-specific responses (i.e., stronger NPS and SIIPS expression in the Heat group than Salt and Sugar groups). Signature patterns were also implicated in mediation analyses, particularly when taking valence into account. Responses to predictive cues were consistent across domains, with low cues eliciting larger medial OFC deactivation during anticipation regardless of modality or aversiveness. Whole brain multilevel mediation analyses revealed additional evidence of common effects: All groups showed right DLPFC deactivation with high cues, and stronger right dorsal posterior insula activation with the more pleasant cue, and cue effects on subjective intensity were mediated by responses to stimuli in the left middle insula, while cue effects on subjective valence were mediated by the right thalamus. We found only minor evidence of domain-specificity by testing moderation by either modality (Pain vs. Taste) or Aversiveness (Aversive vs. Appetitive), with the strongest differences in the right thalamus - posterior to the thalamic region engaged in valence mediation. These findings suggest that predictive cues modulate perception primarily through domain-general mechanisms. Here we discuss these findings and their implications in more detail.

As the magnitude of stimulation increased, participants generally rated the temperature or concentration higher across all stimuli. They also reported increased unpleasantness for heat and salt and – to a lesser extent – pleasantness for sugar as magnitude increased, during both the calibration and fMRI visits. These differences were accompanied by changes in primary circuits identified in prior work. Specifically, high-magnitude heat stimuli, compared to low-magnitude stimuli, induced robust pain-related activations in a network of regions identified via meta-analysis in previous pain and placebo studies (Jensen et al., 2016), such as the ACC, MCC, and insula. In contrast, for salt and sugar, activations related to high vs. low magnitude were primarily observed in gustatory processing areas (Small et al., 2003), such as SII, middle frontal gyrus, and temporal regions. These activations are consistent with findings from other studies using event-related potentials (Ohla et al., 2010) and neuroimaging (Kobayashi et al., 2004; Avery et al., 2020) which linked these regions to intensity-related gustatory perception or imagery. Overall, our results provide strong evidence that the stimuli manipulation was effective, with heat inducing pain-related neural activations and salt and sugar eliciting gustatory processing-related activations. This establishes a solid foundation for our subsequent analysis of cue effects.

Classification further revealed both pain-related neural representations and general aversive processing. Pain-related neural signature patterns revealed modality-specificity, with greater expression of a classifier developed to predict both nociceptive pain (NPS, Wager et al., 2013) and stimulus-independent pain (SIIPS, Woo et al., 2017) in the Heat Group, compared to Salt and Sugar Groups. Interestingly, changes in salt concentration were associated with significant activation of the NPS and SIIPS patterns, albeit to a lesser extent than changes in temperature. While one could argue that the stronger neural expression for heat is simply due to its overall higher activation levels, the absence of a significant difference in the domain-general negative affect pattern (Čeko et al., 2022) between heat and the taste stimuli suggests that the neural signatures exhibit selective sensitivity to different modalities. The evidence indicates that heat-related activations cannot be attributed solely to nociception but instead reflect nociceptive- and pain-specific processing as well as processes involved in general aversiveness.

When presented with a high cue compared to a low cue, participants expected higher intensity and unpleasantness for heat and salt, and higher intensity and pleasantness for sugar. Interestingly, salt was expected to be the most intense, while sugar was expected to be the most pleasant, which was consistent with cue effects on subjective perception of moderate stimuli, as discussed further below. However, we observed no differences across groups when we examined cue-evoked anticipatory activation.

Neuroimaging results showed strong bilateral medial OFC activation in response to high versus low cues, independent of modality or aversiveness. Previous studies support the role of the medial OFC in encoding a general anticipatory value signal for both aversive and positive effects (Nitschke, Sarinopoulos, et al., 2006; Levy & Glimcher, 2011; Roy et al., 2012; Metereau & Dreher, 2015). Additionally, we found a trend toward a negative relationship across participants between cue effects on OFC activation and expected intensity ratings, suggesting that participants who expected higher intensity showed less stronger effects of predictive cues on anticipatory OFC activation. In a monetary reward decision making study, Yan and colleagues (2016) found the multivariate pattern of left and medial OFC matched the pattern for magnitude of reward during anticipation before making a choice. For anticipation after making a choice, they found a correlation between the patterns of the left OFC and valence. We speculate motivation or the specific anticipated outcome may play a role for the representation of magnitude or valence in OFC during anticipation. Given that participants in our study were self-learning without explicit instructions, they likely anticipated a general cue value (magnitude/intensity) rather than detailed outcomes. We note that we asked participants about subjective expectations between each block, rather than on every trial, to avoid drawing attention to the purpose of the study, which precluded more dynamic evaluations between OFC anticipatory activation and dynamic expectations. Future studies should incorporate trial-based assessments to further clarify associations between explicit expectations and OFC representation during anticipation.

Our study revealed behavioral and neuroimaging evidence for significant cue effects on perception across different sensory domains. Behaviorally, we observed significant main effects of cue on both intensity and valence ratings for the key medium stimuli which were crossed with cues. In all models, high cues were perceived as more intense and unpleasant (heat, salt)/pleasant (sugar) than low cues. Notably, the absence of significant interactions between cue and group in any model suggests that the cue effect on subjective perception was consistent across different domains, although expectations did vary across groups. Single trial mediation results revealed domain-general mediators of cue effects, whether we ignored outcome aversiveness (intensity mediation) or took aversiveness into account (valence mediation). Regardless of aversiveness, high cues modulated stimulus-evoked activation in the prefrontal cortex, with high cues eliciting increased activation in left DMPFC and decreased activation in right DLPFC, relative to low cues. Previous studies have implicated these regions in cue-based modulation of subjective pain (Atlas et al., 2010; Atlas & Wager, 2014; Woo et al., 2017) and most studies of placebo analgesia show increases in DLPFC with expectations for pain reduction (Wager et al., 2004; Atlas & Wager, 2014), consistent with a general role in emotion regulation and cognitive control (Kohn et al., 2014; Morawetz et al., 2017). One study showed that both DMPFC and DLPFC are engaged in conflict decision-making and in modulating the magnitude of value change during reversal learning (Mitchell et al., 2009). The opposing directions of activation may reflect distinct cognitive processes underlying DMPFC and DLPFC in managing conflicting information.

We also accounted for differences in aversiveness across groups and measured the impact of cues representing relative aversiveness/pleasantness (Heat and Salt: Low Cue > High Cue, Sugar: High Cue > Low Cue). FDR correction showed significant cue effects on activation in the right dorsal posterior insula when accounting for valence and controlling for group. This indicates valence-specific cue effects on affect in the brain. The dorsal posterior insula has been hypothesized to be pain-specific (Segerdahl et al., 2015) and was found to show pain-specific processing in previous studies of expectancy-based modulation across domains (Sharvit et al., 2019). Our finding of similar cue-induced modulation on activation regardless of group is inconsistent with these results and suggest that modulation of this region is *not* specific to pain. Interestingly, all groups showed greater activation of this region with the relatively less aversive cue, which is also inconsistent with prior work (Atlas et al., 2010). Our findings may be consistent with models that suggest this region plays a role in detecting aversiveness of internal states and shifting behavioral strategies (Gehrlach et al., 2019). Accounting for differences in valence also revealed common cue effects in the left DMPFC, based on FDR-correction, and in a network of regions implicated in emotion and affective processing based on uncorrected results, including the bilateral amygdala, medial orbitofrontal cortex, bilateral insula, dorsal anterior cingulate, and other regions. Despite the fact that these common emotion-related regions were impacted by predictive cues, with greater activation for the cue that predicted the more pleasant outcome, we observed no effect of cues on the negative affect signature pattern. Instead, we found opposing trends for cue effects on the NPS and SIIPS: the NPS exhibited greater activation following pleasant cues, whereas the SIIPS demonstrated greater activation following unpleasant cues.

Importantly, we found that brain mediators of predictive cue effects on behavioral ratings were largely consistent across domains. Specifically, we observed: 1) Dynamic cue effects on intensity were mediated by stimulus-evoked activation in the left anterior/middle insula and the left visual cortex; and 2) Dynamic cue effects on valence ratings were mediated by the right thalamus based on FDR-correction. All mediators were driven by covariance, such that individuals who had stronger cue effects on stimulus-evoked activation also had stronger associations between activation and subjective ratings (positive associations for insula and thalamus; negative associations in visual cortex). The absence of modality-specific moderation for either rating further supports the shared nature of these effects. The insula and thalamus regions we identified are nearly identical to regions previously identified in mediating cue effects on pain (Atlas et al., 2010); the same insula region has also been shown to be modulated by predictive cues during taste perception (Nitschke, Dixon, et al., 2006). The thalamus, which is anatomically and functionally connected to the striatum, has been implicated in coding reward and valence, particularly in the context of action-specific values (Liu et al., 2011; FitzGerald et al., 2012). The visual cortex has been shown to encode pain-related cue information (Kim et al., 2024) and interact with the attention network in response to visual cues, which in turns engage the gustatory networks (Veldhuizen et al., 2012). This comprehensive evidence for shared, rather than domain-specific, cue effects on behavioral outcomes suggests that participants across different domains utilize similar implicated information and underlying psychological processes from cues onto the behavioral outcomes. These findings align with our previous placebo studies, which demonstrated the transferability of placebo effects across domains (e.g., from heat or noise to negative pictures or empathy for pain) (Zhang et al., 2011; Zhao et al., 2015; Zhao et al., 2020). The success of these transfers was predicated on the shared belief in the treatment’s ability to generally improve personal suffering.

Our analysis unveiled a few modality- and aversiveness-moderated cue effects in the brain. The only moderation effect that survived correction was a Path *a* effect (High Cue - Low Cue) in the right thalamus, moderated by Group. This effect was primarily driven by greater taste-induced thalamic activation in the Sugar group, whereas the Salt and Heat groups showed deactivation, with the strongest deactivation observed in the Heat group. This finding aligns with previous evidence that thalamic activity contributes critically to cue-related processing in the gustatory cortex and taste reward mechanisms (Schroy et al., 2005; Samuelsen et al., 2013). For cues representing intensity, moderation analysis revealed that the right temporal gyrus was differently modulated by modality (Heat vs. Tastes), with heat showing higher negative activation for high vs. low intensity-relevant cues compared to tastes. Previous studies have observed deactivation of the temporal gyrus during nociceptive stimulation, with anti-hyperalgesic effects more strongly associated with reduced pain-evoked deactivations than activations (Iannetti et al., 2005; Kong et al., 2010). Signature pattern analysis showed a trend toward moderated mediation for the SIIPS pattern, primarily driven by a more positive mediation effect for the Heat Group compared to the Salt Group. This finding provides further support for the specificity of the SIIPS pattern to Heat relative to other modalities (Woo et al., 2017). For cues representing pleasantness (Heat and Salt: Low Cue > High Cue, Sugar: High Cue > Low Cue), moderation analysis revealed evidence of both aversiveness and modality effects. Sugar elicited more positive activation in left DMPFC compared to Salt, while Heat showed less right DMPFC activation relative to Salt in response to cue expectations. The medial prefrontal cortex has been implicated in processing both the physicochemical and hedonic properties of taste (Jezzini et al., 2013). Though we also found higher activation of lateral PFC in Heat compared to Salt/Sugar, consider the broader and stronger activation in heat, we could not attribute this activation to modality specificity.

This study has several limitations. Firstly, we encountered difficulties in delivering a sucrose solution of comparable high concentration to that used in the initial visit. This resulted in overall smaller activation and effect sizes in the Sugar group. Future studies should explore more effective methods for delivering sucrose solutions or consider alternative rewarding stimuli to ensure matched perception across domains. Secondly, while our between-group design was initially intended to elucidate cue effects across domains, it may limit the ability to compare perception and cue effects within individuals. To address this, we propose implementing a within-subject design in future studies, allowing each participant to experience both heat and taste stimuli for more direct comparisons.

In conclusion, our study demonstrates that cue effects can be induced through uninstructed associative learning across multiple sensory domains. Heat stimuli exhibited unique representations compared to tastes, selectively activating nociceptive- and pain-related neural patterns. During the anticipation phase, we observed shared activation in the OFC associated with cue effects across domains. In the outcome stage, cue effects on behavioral outcomes revealed predominantly domain-general activations across modalities for both intensity, such as in the insula, and pleasantness, such as in the thalamus. Some domain-specific activations were observed for predictive cues without being necessarily relevant to behavior. These findings suggest that while different sensory modalities engage distinct functional processes and neural activations, learned cues influence anticipation and behavioral outcomes in a largely domain-general manner, spanning both aversive and appetitive experiences. Overall, this highlights both shared and distinct neural representations underlying the processing of uninstructed, expectation-based information across diverse sensory experiences.

## Methods

The study was approved by the National Institutes of Health Institutional Review Board (Protocol 15-AT-0132; ClinicalTrials.gov Identifier: NCT02446262) and followed the Declaration of Helsinki guidelines for ethical conduct of human research. Analyses were preregistered through Aspredicted (https://aspredicted.org/GXN_FYL).

### Participants

Healthy volunteers from a community sample were screened for this two-visit study. Potential volunteers were deemed ineligible if they had a history of sensory, neurological, or psychiatric disorders, substance abuse, or any major medical condition that could affect somatosensation. Additionally, volunteers who regularly took medication known to affect pain or heat perception were excluded. Eligible volunteers were between the ages of 18 and 50, right-handed, not pregnant, and did not have any condition or device that would pose a risk during the fMRI portion of the study. Participants were also required to have abstained from recreational drugs for the past month and from pain relievers within a time frame equivalent to five half-lives. 78 participants provided consent during an initial screening and Calibration Visit (see Procedure in Figure 1A). 60 participants (37 females, age: 29.58 ± 8.57 years) were determined to be eligible and subsequently completed fMRI scanning on a follow-up visit.

### Materials

#### Thermal pain stimulation

A Medoc Pathway Pain and Sensory Evaluation System (Medoc Advanced Medical Systems Ltd., Ramat Yishay, Israel) with Medoc Station software was used to deliver thermal stimuli to participants’ left volar forearm. We used an Advanced Thermal Stimulator (ATS) thermode with a square contact area of 16 x 16 mm². As addressed below, thermal stimuli lasted for 8 seconds (1.5 seconds ramp up, 5 seconds at peak, 1.5 seconds ramp to baseline) and ranged from 32°C to 50°C.

#### Appetitive and aversive tastants

We administered liquid tastants to all participants during the Calibration Visit and to Sugar and Salt groups during the fMRI Visit. Sterile solutions were prepared by the NIH Compounding Pharmacy. A combination of Potassium Chloride and Sodium Bicarbonate (2.5 mm NaHCO3 + 25 mm KCl; 1 mL per trial), which mimics saliva, was used as the neutral solution, consistent with previous taste-fMRI studies (Simmons et al., 2013; Avery et al., 2020). Liquid sodium chloride ranging from 0.2-2M (0.5 mL per trial) was used as the aversive solution. Liquid sucrose ranging from 0.2-2M (0.5 mL per trial) was used as an appetitive solution. During the Calibration Visit, participants self-administered tastants using syringes. During the fMRI Visit, a custom-built pneumatically-controlled gustometer (Simmons et al., 2013; Avery et al., 2020) was used to precisely deliver pressurized tastants through IV-tubing to a 3D-printed mouthpiece. Because we used IV tubes for tastant delivery, we used a maximum of 1M sucrose solution during the fMRI Visit as higher concentrations were more viscous and thus did not move smoothly through the tubing. On each visit, we ensured that participants did not consume more than 1400 mg of sodium (salt), in accordance with the Food and Drug Administration’s recommended daily allowance, and instructed participants to reduce sugar and salt intake for the rest of the day to stay within World Health Organization guidelines.

#### Task delivery and ratings

Stimuli were presented using Psychtoolbox (http://psychtoolbox.org/) and synched with painful stimulation and taste delivery using a LabView (National Instruments) program. During the Calibration Visit, participants provided verbal ratings of intensity, unpleasantness, and pleasantness using a 0-10 continuous visual analogue scale (VAS), where 10 was defined as most intense/unpleasant/pleasant sensation imaginable. We computed the mean intensity and valence ([unpleasantness – pleasantness] for heat and salt; [unpleasantness – pleasantness] for sugar) for use in linear regressions to select temperatures and concentrations for the subsequent fMRI visit. During the fMRI experiment, participants provided ratings using an MRI-compatible trackball (Current Designs, Philadelphia PA). Intensity was again rated on a 0-10 VAS, while valence was measured on a bivalent scale ranging from -10 (most unpleasant imaginable) to 10 (most pleasant imaginable).

### Procedures

#### Day 1: Calibration Visit

Following an initial phone screen, volunteers deemed potentially eligible were invited to complete a Calibration Visit (Figure 1A) at the NIH Outpatient Clinic. 78 participants provided informed consent and underwent a nursing evaluation and a medical exam if they had not had one within the prior year. 7 participants were determined to be ineligible. Following consent and confirmation of eligibility, 71 participants completed questionnaires (which are outside the scope of the current study and may be analyzed in future research; see Supplemental Methods), followed by a calibration procedure in which they rated the intensity and valence of varying levels of heat, salt solution, sucrose solution, and neutral solution in counterbalanced order. Calibration procedures were adapted from an adaptive staircase pain calibration procedure we had used previously (see Amir et al., 2021 for complete details). In brief, on each trial for each modality, participants experienced the stimulus and rated its intensity, pleasantness, and unpleasantness following offset using a 0-10 VAS. Each VAS included the following anchors: 0 (no sensation), 5 (moderate intensity/pleasantness/unpleasantness), and 10 (most intense/pleasant/unpleasant sensation imaginable). When rating heat stimuli intensity, participants were also instructed that 1 corresponded to nonpainful warmth, 2 corresponded to the beginning of pain sensation (i.e., pain threshold), and 8 corresponded to highest tolerable pain (i.e., pain tolerance), consistent with prior work (Atlas et al., 2010; Amir et al., 2022). Heat temperatures were selected iteratively based on an iterative regression of temperature and intensity across 24 trials, consistent with previous work (Amir et al., 2021). Tastant concentrations were prepared in advance by the NIH Pharmacy. We tested 11 concentrations varying from 0-2M for salt and sugar (22 trials total) and 5 concentrations for the neutral solution varying from 0 to 100% in 25% increments (10 trials total). Following the calibration, we evaluated the overall correlation between stimulus level and perception based on intensity ratings. Consistent with previous work (Atlas et al., 2010; Amir et al., 2022), participants were only eligible to continue with a given modality if their correlation coefficients (r^2^) between stimulus intensity and perceived intensity exceeded 0.4. For each participant and each modality, we applied a regression model that related stimulus temperature or concentration with the mean intensity/valence to determine stimulus levels corresponding to low (2), medium (5), and high (8) intensity and valence perception. These individual levels were used during the fMRI visit. As we could not deliver higher than 1M sucrose through the IV tubing, maximum sucrose concentrations were adjusted (i.e., if the high intensity exceeded 1M, participants received 1M sucrose as their maximum concentration). Finally, for neutral solutions, we selected the concentration whose mean intensity was closest to 0.

#### Day 2: fMRI Visit

Following the Calibration Visit, eligible participants were pseudorandomly assigned to one of three task groups (Heat, Salt, or Sugar; Figure 1B). Participants were excluded from Sugar or Salt groups if they perceived salt as pleasant or sugar as unpleasant, respectively, and from the Heat group if their tolerance exceeded safe limits (see Supplemental Methods). Group assignment was done prior to the fMRI visit so that the NIH Pharmacy could prepare tastants in advance using the concentrations from the Calibration Visit. Upon arrival, participants provided informed consent for fMRI scanning, confirmed continued eligibility, and completed a set of clinical visit and state assessment questionnaires that may be analyzed in future research. Participants then underwent a follow-up calibration to verify the individual threshold and tolerance levels. Complete details can be found in Supplemental Methods.

Following calibration, participants completed a short practice task and were then situated in the MRI scanner and underwent structural scans followed by five runs of fMRI scanning (12 trials per run). All participants were instructed to focus on the relationship between the visual cue and the intensity of the following stimulus. As shown in Figure 1B, participants first underwent conditioning (10 trials) during which one cue was followed by a stimulus calibrated to elicit high intensity ratings (level 8; “High”; *M*_Heat_ = 48.66°C, *SD*_Heat_ = 1.35; *M*_Salt_ = 1.71M, *SD*_Salt_ = 0.33; *M*_Sugar_ = 0.99M, *SD*_Sugar_ = 0.04) and a second cue was followed by a stimulus calibrated to elicit low intensity ratings (level 2; “Low”; *M*_Heat_ = 42.87°C, *SD*_Heat_ = 1.44; *M*_Salt_ = 0.35M, *SD*_Salt_ = 0.18; *M*_Sugar_ = 0.27M, *SD*_Sugar_ = 0.12). During a subsequent test phase (50 trials), each cue was followed by either the conditioned level or a stimulus calibrated to elicit ratings of medium intensity (level 5; “Medium”; *M*_Heat_ = 45.86°C, *SD*_Heat_ = 1.18; *M*_Salt_ = 1.01M, *SD*_Salt_ = 0.31; *M*_Sugar_ = 0.66M, *SD*_Sugar_ = 0.09), consistent with previous work (Atlas et al., 2010). As shown in Figure 1B, each trial began with a 2-second visual cue (counterbalanced across participants) followed by a 2-3 second jittered delay, during which a fixation cross was displayed. The 8-s heat or tastant outcome stimulus was then delivered via the thermode or gustometer while word “Focus” was presented on the screen. Participants in the Salt and Sugar groups were then prompted to swallow the tastant (“Swallow”) while participants in the Heat group were told to “Wait”. This period lasted 1 second. After a 1-2 second transition, participants used the trackball to rate perceived intensity and valence (5 seconds, respectively). Missing trials (mean per subject < 13%) were omitted from behavioral analyses and single-trial fMRI analysis. Following a 1-2 seconds jittered delay, participants in the Salt and the Sugar groups received 1mL of the neutral solution (*M*_Salt_ = 73.75% concentration, *SD*_Salt_ = 28.65; *M*_Sugar_ = 73.75% concentration, *SD*_Sugar_ = 30.65) and were asked to rinse and then swallow the liquid. Participants in the Heat group saw the word “Wait” during this delay. A 2-3 second fixation cross was presented prior to the next trial. Participants rated expected intensity and valence in response to each cue at the end of each run. See Design and Procedure in Figure 1B. After the fMRI scan, participants completed post-task questions/questionnaires regarding perception during the scan and in general (see Supplemental Methods) and then were debriefed and dismissed.

### Behavioral data analysis

Behavioral analyses were conducted using Lme4 (https://cran.r-project.org/web/packages/lme4/index.html) in R version 4.2.1 (https://www.r-project.org/). We analyzed both expectancy ratings and ratings in response to stimulation, and separately analyzed intensity and valence ratings. Consistent with our preregistration, our main models included fixed effects for Cue, Stimulus Level, Time (trial for stimulus ratings, block for expectancy ratings), and Group (Heat, Salt, and Sugar; dummy-coded). Follow-up analyses were restricted to medium trials. We also conducted exploratory analyses that modeled Modality (Heat vs Tastes) and Aversiveness ([Heat and Salt] vs Sugar) rather than Group as second-level factors. Aversiveness and Modality were dummy coded. Cue, Stimulus Level, and Time were mean-centered. Model comparison was used to evaluate goodness of fit and specification of random factors (Stimuli, Cue, Time, and Subject) and interactions. For details on model specification and results of model comparison, see Supplemental Methods, Table S1-1, Table S1-2, and Table S1-3.

### fMRI acquisition

fMRI data were collected at the NIMH fMRI Facility at the NIH Clinical Center using a General Electric (GE) 3 Tesla MR-750 scanner with a 32-channel head coil. Images were acquired using a multi-echo echo-planar imaging (EPI) sequence (echo times [TEs] = 8.8/21.2/33.6 ms; 35 oblique slices for each echo; slice thickness = 3 mm; in-plane resolution = 3 x 3 mm; square field of view [FOV] = 216 mm; bottom-up sequential acquisitions; repetition time [TR] = 2000 ms; flip angle = 75°; voxel size = 3 x 3 x 3 mm; acquisition matrix: 72 x 72). Each slice was oriented in the axial plane but rotated 30 degrees clockwise relative to the AC-PC line to reduce dropout in the VMPFC (Deichmann et al., 2003). This orientation resulted in some areas at the top of the parietal and occipital lobes being outside the scanning window for many participants, and as such, our results are agnostic as to the role of superficial parietal and occipital lobes. Ultra-high resolution T1-weighted Magnetization-Prepared Rapid Gradient-Echo and Proton Density sequences were used to provide anatomical references for the fMRI analysis. The parameters for these sequences were: axial prescription; 172 slices per slab; slice thickness = 1 mm; spacing between slices = 1 mm; square FOV = 256 mm; image matrix = 256 x 256; TE = 3.504 ms; TR = 7948 ms; flip angle = 7°; voxel size = 1 x 1 x 1 mm.

### fMRI preprocessing

All three echoes of fMRI data were preprocessed using AFNI (http://afni.nimh.nih.gov/afni) with an Optimally Combined (OC) pipeline implemented using afni_proc.py (Cox, 1996). First, the initial 10 seconds of data were discarded to ensure the scanner reached a steady state. Next, a de-spiking interpolation algorithm (3dDespike) was applied to remove transient signal spikes from the EPI data. Slice timing correction was then performed using the 3dtshift function. To address spatial distortion artifacts, an EPI scan acquired with reverse blip was used for correction. Head motion was estimated using the 3dVolreg function. We combined across echoes using a voxel-wise linear weighted (optimal) combination to optimize signal-to-noise ratio across brain regions with differing T2* values (Poser & Norris, 2009; Posse, 2012). To enhance the signal-to-noise ratio, a 4 mm full-width at half-maximum (FWHM) Gaussian kernel was applied. Finally, the signal intensity for each volume was normalized on a voxel-wise basis to reflect percentage signal change. Anatomical scans were warped to standard MNI space using a nonlinear transformation implemented in AFNI’s @SSwarper (https://afni.nimh.nih.gov/pub/dist/doc/program_help/@SSwarper.html). The resulting nonlinear transformation matrices were applied to register subject-level statistical maps to MNI space, preserving the original voxel resolution of 3 x 3 x 3 mm without resampling. Subjects with an average Euclidean-normalized derivative of > 0.3 during the task were excluded from the group-level analysis due to excessive head motion. As a result, eight subjects were excluded from the original group of 60 subjects, leaving 52 subjects (18 Heat, 17 Salt, 17 Sugar) for the univariate analysis and Multivariate Pattern Analysis (MVPA). Due to technical issues with our trackball leading to missing ratings, additional 5 participants were excluded from mediation analyses examining associations with intensity (total n = 47, 15 Heat, 17 Salt, 15 Sugar) and additional 6 participants were excluded from mediation analyses examining associations with pleasantness (total n =46, 14 Heat, 17 Salt, 15 Sugar).

### Subject-level fMRI analyses

#### Hemodynamic response function

To ensure that our models captured brain responses to outcomes regardless of modality, we extracted raw activation from the bilateral middle insula and right dorsal posterior insula and fitted various hemodynamic response functions (HRFs) to the outcome-evoked response after preprocessing (see Supplemental Methods). AFNI’s MION HRF was determined to provide the best fit for stimulus-evoked responses across groups and was therefore used in first-level analyses.

*General linear models (GLM)* were implemented using AFNI’s 3dDeconvolve (https://afni.nimh.nih.gov/afni/doc/help/3dDeconvolve.html). For each individual, the regression model included regressors for each primary event of interest, which were constructed using the “regress_basis_multi” function in AFNI’s 3dDeconvolve program. The present analyses focus on responses to cues, which were modeled using 2-second boxcars convolved with AFNI’s default HRF (“BLOCK(2,1)” in AFNI), and responses to stimulus outcomes, which were modeled with an 8-second boxcar convolved with the MION HRF. We modeled separate regressors for each condition (i.e., High Cue, Low Cue, High Stimulus, Low Stimulus, High Cue + Medium Stimulus, Low Cue + Medium Stimulus). We also included a 1-second event at stimulus offset to model swallow-related activity, a 10-second boxcar to model the rating period, and a 4-second boxcar to model the rinse period; these were not analyzed in the present study. Motion parameters were incorporated as covariates of no interest to account for potential confounding effects.

*Single trial analyses* require estimates per trial, rather than per condition. Whole brain trial-level estimates were generated using AFNI’s 3dREMLfit program (https://afni.nimh.nih.gov/pub/dist/doc/htmldoc/statistics/remlfit.html). We included the same regressors of no interest and cues as the GLM, but modeled each outcome trial (i.e., heat or tastant delivery) separately with its own 8-second MION HRF. This generates a coefficient per voxel per trial which was used as a measure of trial-evoked activation and used in voxel-wise multilevel mediations, described in more detail below.

### Group-level fMRI analyses

#### Mass univariate analyses

For group-level analysis, we initially performed a full factorial design with Statistical Parametric Mapping (SPM12; https://www.fil.ion.ucl.ac.uk/spm/software/spm12/) to explore brain activation associated with varying stimuli intensities. Six contrasts were constructed, comparing responses to stimulus outcome as a function of Level (High vs. Low) for Heat, Salt, and Sugar, in both directions. Medium stimuli were excluded from this analysis to prevent confounding from cue effects, which could obscure activations attributable solely to stimulus level (i.e., changes in temperature or concentration). Subsequently, another full factorial design was employed to examine activations during anticipation as a function of Cue (High vs. Low), considering both directions and analyzing both across and within groups. A conjunction analysis was conducted to identify shared neural activations across all three groups and each pair of groups for each contrast.

#### Multivariate pattern analysis (MVPA)

We used MVPA implemented in the Decoding Toolbox (https://sites.google.com/site/tdtdecodingtoolbox/) via Matlab to further validate the activity identified in the univariate analysis. Group-level analysis was conducted using SPM. We performed a one-sample t-test to evaluate the whole-brain patterns capable of differentiating between High and Low cues with greater than 50% accuracy, based on individual accuracy maps extracted from all participants. For additional information, see subject-level MVPA analysis in Supplemental Methods.

#### Multilevel mediation and moderation analyses

Voxelwise multilevel mediation was implemented in Matlab using the M3 toolbox, which implements linear mixed models (for complete details, see (Wager et al., 2008; Wager et al., 2009; Atlas et al., 2010)). We used multi-level mediation analysis to identify neural activations that explained cue effects on subjective perception in response to medium stimuli and applied moderated mediation to determine whether effects differ across groups. Consistent with previous work (Atlas 2010, 2021), our mediation models designated variations in Cue (High - Low) as the input variable (X), ratings on medium trials as the outcome variable (Y), and voxelwise activation on medium trials as the potential mediator (M). Trial-by-trial voxel-wise activation was operationalized using single trial estimates from AFNI’s 3dREMLfit as described above. Because single trial estimates are sensitive to noise, we evaluated variance inflation factors (VIFs) in a design matrix incorporating trial estimates and nuisance regressors and removed any trials whose VIFs exceeded 2 prior to analysis, consistent with prior work (Atlas et al., 2010; Atlas et al., 2014; Atlas et al., 2022). As with prior work (Atlas et al., 2010), Path *a* in our mediation framework captures the effect of predictive cues on brain activation (X - M), Path *b* evaluates the association between activation and rating when controlling for cue (M - Y), and the mediation effect tests the indirect effect of cues on ratings through changes in brain activation, or the difference between the total effect (effect of X on Y) and the direct effect when controlling for the mediator. In multilevel mediation, the mediation effect (c - c’) is equivalent to the product of Paths *a* and *b* plus their covariance (i.e., c-c’ = a*b + cov(a,b); Kenny et al., 2003). We thus extracted responses from mediators and inspected outcomes to determine whether they were driven by consistent responses or variations across individuals. Significance was tested using bootstrapping to account for the covariance between paths A and B.

We separately searched for brain mediators of cue effects on subjective intensity and cue effects on subjective valence. Our intensity mediation treated Cue [High > Low] as the input variable for all groups. However, since high cues were associated with the more unpleasant outcome for Heat and Salt Groups, whereas they were associated with the more pleasant outcome for the Sugar Group, we accounted for group differences in overall aversiveness in our valence mediation. We accomplished this by reversing the cue contrast depending on stimulus aversiveness: While Path *a* for the Sugar Group was the same as in the intensity model (High > Low), Path *a* was coded as (Low > High) for the Heat and Salt Groups. Since trial-wise valence/pleasantness ratings served as Y, this leads to an overall positive association between X and Y regardless of group.

For each mediation, we evaluated mediation regardless of group and implemented bootstrapping to evaluate significance. We also conducted moderation analyses to explore whether path effects varied as a function of Group (two moderators: [Heat > Salt], and [Salt > Sugar]), Modality (Heat > Taste), or Aversiveness (Appetitive > Aversive), and to evaluate results when controlling for Group/Modality/Aversiveness.

### Statistical thresholding and regions of interest

#### Neural Signature Patterns

To explore whether pain-specific networks were modulated by stimulus intensity or learned expectations, we used three validated neural signature patterns (NPS, Wager et al., 2013; SIIPS, Woo et al., 2017; and the negative affect pattern, Čeko et al., 2022). We computed the dot product of the pattern weights and neural betas and contrast maps (e.g., High vs. Low stimulus level) and performed one-sample t-tests for each group and used one-way analysis of variance (ANOVA) for each pattern to determine whether there were significant differences in pattern responsiveness across groups. We anticipated identifying a distinct fit of Heat responses within pain-specific neural patterns (NPS and SIIPS) but expected no differences with the aversiveness-general pattern (the negative affect pattern) between Heat and Salt. Multiple comparisons among three groups were adjusted using Bonferroni correction. The same approach was used to evaluate whether there were significant effects on signature patterns for each path of our mediation analyses.

#### Regions of interest

Small volume correction was performed for each analysis, using a region-of-interest (ROI) mask from our previous meta-analysis, which identified placebo-related increased activations (Atlas & Wager, 2014). Consistent with our preregistration, we focus on responses to medium intensity pain and non-pain stimuli in SI, SII, ACC, insula, operculum, thalamus, periaqueductal gray, striatum, DLPFC, OFC/VMPFC, and the amygdala. Family Wise Error (FWE) correction (*p* < .05) was applied for small volume correction.

#### Univariate and MVPA analyses

For whole-brain exploratory analysis, we applied an initial voxel-level statistical threshold of *p* < .001 (uncorrected), with a cluster extent threshold of 10 contiguous voxels to enhance result robustness. We further performed FWE correction (*p* < .05) and reports all activations survived this stricter correction.

#### Mediation and moderation analyses

We report mediation and moderation results that survived whole-brain False Discovery Rate (FDR) correction (*q* < .05), since there were no results that survived correction within regions involved in pain and placebo analgesia. Whole brain uncorrected exploratory results are reported at a voxelwise threshold of *p* < .001 with at least 3 contiguous voxels; clusters were defined based on being contiguous with additional voxels at *p* < .005 and *p* < .01, consistent with prior work (Atlas et al., 2010).

## Supporting information

Supplement Results

## Acknowledgements

This research was supported by the Intramural Research Program of the NIH (ZIA-AT000030, PI LYA). We thank the NIMH Section on Instrumentation for development of the gustometer and associated software, the NIH Compounding pharmacy and metabolic kitchen for creation of the tastants used in the experiments, and Adebisi Ayodele for safety guidelines. We also thank Linda Ellison-Dejewski, Adebisi Ayodele, nurses from the OP4 Behavioral Health Clinic, and Dr. Daniel Pine for clinical support, the NIMH FMRI Facility for assistance with MRI scanning, Lawrence Tello and Carlos Melendez for assistance with pilot versions of the study, and Olga Oretsky, Rachel Weger, and Carolyn Amir for assistance with participant scheduling and determining eligibility during Calibration Visits.

