## Supplementary material for "Domain-general neural effects of associative learning and expectations on pain and hedonic taste perception": Table S2

**Table S2.** Results of the winning Group models for intensity ratings and valence ratings across all trials.^a^

|  | Dependent Variable | **Group Factor** | **Factors** | | **Estimate** | | | | | | **T Value** | | **P Value** |
| --- | --- | --- | --- | --- | --- | --- | --- | --- | --- | --- | --- | --- | --- |
|  | Intensity | Group (Heat vs. Salt vs. Sugar)^b^ | Group_PainvsSalt | | -1.043 | | | | | | -3.329 | | 0.002** |
|  |  |  | Cue | | 0.309 | | | | | | 5.054 | | <0.001*** |
|  |  |  | Stimuli | | 1.940 | | | | | | 17.556 | | <0.001*** |
|  |  |  | Trial | | 0.003 | | | | | | 1.181 | | 0.238 |
|  |  |  | Group_SaltvsSugar | | 0.149 | | | | | | 0.478 | | 0.634 |
|  |  |  | Group_PainvsSalt * Cue | | 0.091 | | | | | | 1.025 | | 0.310 |
|  |  |  | Group_PainvsSalt * Stimuli | | 0.355 | | | | | | 2.228 | | 0.030* |
|  |  |  | Stimuli * Cue | | -0.052 | | | | | | -0.416 | | 0.679 |
|  |  |  | Group_PainvsSalt * Trial | | -0.014 | | | | | | -4.382 | | <0.001*** |
|  |  |  | Cue* Trial | | -0.003 | | | | | | -1.341 | | 0.180 |
|  |  |  | Stimuli * Trial | | 0.009 | | | | | | 3.368 | | <0.001*** |
|  |  |  | Group_SaltvsSugar * Cue | | 0.063 | | | | | | 0.730 | | 0.468 |
|  |  |  | Group_SaltvsSugar * Stimuli | | 0.575 | | | | | | 3.670 | | <0.001*** |
|  |  |  | Group_SaltvsSugar * Trial | | 0.001 | | | | | | 0.203 | | 0.839 |
|  |  |  | Group_PainvsSalt * Cue * Stimuli | | 1.055 | | | | | | 5.832 | | <0.001*** |
|  |  |  | Group_PainvsSalt * Cue * Trial | | -0.001 | | | | | | -0.163 | | 0.871 |
|  |  |  | Group_PainvsSalt * Stimuli * Trial | | -0.003 | | | | | | 0.635 | | 0.525 |
|  |  |  | Stimuli * Cue * Trial | | 0.003 | | | | | | 1.239 | | 0.215 |
|  |  |  | Group_SaltvsSugar * Cue * Stimuli | | 0.507 | | | | | | 2.840 | | 0.006** |
|  |  |  | Group_SaltvsSugar * Cue * Trial | | -0.006 | | | | | | -1.815 | | 0.070 |
|  |  |  | Group_SaltvsSugar * Stimuli * Trial | | 0.008 | | | | | | 2.041 | | 0.041* |
|  |  |  | Group_PainvsSalt * Cue * Stimuli * Trial | | 0.005 | | | | | | 1.373 | | 0.170 |
|  |  |  | Group_SaltvsSugar * Cue * Stimuli * Trial | | -0.001 | | | | | | -0.259 | | 0.796 |
|  | Valence | Group (Heat vs. Salt vs. Sugar)^c^ |  | | | | **Estimate** | | | **T Value** | | | **P Value** |
|  |  |  | Group_PainvsSalt | | | | -0.459 | | | -0.833 | | | 0.408 |
|  |  |  | Cue | | | | -0.211 | | | -3.056 | | | 0.004** |
|  |  |  | Stimuli | | | | -1.670 | | | -9.468 | | | <0.001*** |
|  |  |  | Trial | | | | -0.034 | | | -9.952 | | | <0.001*** |
|  |  |  | Group_SaltvsSugar | | | | -3.413 | | | -6.231 | | | <0.001*** |
|  |  |  | Group_PainvsSalt * Cue | | | | -0.143 | | | -1.423 | | | 0.160 |
|  |  |  | Group_PainvsSalt * Stimuli | | | | -1.280 | | | -5.039 | | | <0.001*** |
|  |  |  | Stimuli * Cue | | | | -0.748 | | | -2.732 | | | 0.008** |
|  |  |  | Group_PainvsSalt * Trial | | | | -0.031 | | | 6.369 | | | <0.001*** |
|  |  |  | Cue* Trial | | | | -0.0002 | | | -0.083 | | | 0.934 |
|  |  |  | Stimuli * Trial | | | | -0.009 | | | -2.050 | | | 0.040 |
|  |  |  | Group_SaltvsSugar * Cue | | | | -0.014 | | | -1.401 | | | 0.167 |
|  |  |  | Group_PainvsSalt * Stimuli | | | | -2.154 | | | -8.614 | | | <0.001*** |
|  |  |  | Group_SaltvsSugar * Trial | | | | 0.004 | | | 0.818 | | | 0.413 |
|  |  |  | Group_PainvsSalt * Cue * Stimuli | | | | -0.757 | | | -1.926 | | | 0.059 |
|  |  |  | Group_PainvsSalt * Cue * Trial | | | | 0.001 | 0.215 | | | | | 0.830 |
|  |  |  | Group_PainvsSalt * Stimuli * Trial | | | | 0.007 | 1.193 | | | | | 0.233 |
|  |  |  | Stimuli * Cue * Trial | | | | 0.012 | 2.825 | | | | | 0.005** |
|  |  |  | Group_SaltvsSugar * Cue * Stimuli | | | | -0.079 | -0.204 | | | | | 0.838 |
|  |  |  | Group_SaltvsSugar * Cue * Trial | | | | 0.006 | 1.135 | | | | | 0.256 |
|  |  |  | Group_SaltvsSugar * Stimuli * Trial | | | | 0.001 | 0.135 | | | | | 0.823 |
|  |  |  | Group_PainvsSalt * Cue * Stimuli * Trial | | | | -0.009 | -1.376 | | | | | 0.140 |
|  |  |  | Group_SaltvsSugar * Cue * Stimuli * Trial | | | | -0.002 | -0.402 | | | | | 0.688 |
|  | Dependent Variable | **Group Factor** | **Factors** | **Estimate** | | | | | | **T Value** | | **P Value** | |
|  | Intensity | Aversiveness (Heat and Salt vs. Sugar)^d^ | Stimuli | 1.973 | | | | | | 17.305 | | <0.001*** | |
|  |  |  | Cue | 0.311 | | | | | | 5.009 | | <0.001*** | |
|  |  |  | Aversiveness | 0.135 | | | | | | 0.637 | | 0.527 | |
|  |  |  | Trial | 0.003 | | | | | | 0.777 | | 0.439 | |
|  |  |  | Stimuli * Cue | -0.153 | | | | | | -3.220 | | 0.001** | |
|  |  |  | Stimuli * Aversiveness | 0.419 | | | | | | 3.516 | | <0.001*** | |
|  |  |  | Cue * Aversiveness | 0.041 | | | | | | 0.630 | | 0.532 | |
|  |  |  | Stimuli * Trial | 0.010 | | | | | | 3.389 | | <0.001*** | |
|  |  |  | Cue * Trial | -0.003 | | | | | | -1.302 | | 0.193 | |
|  |  |  | Aversiveness * Trial | 0.002 | | | | | | 0.525 | | 0.601 | |
|  |  |  | Stimuli * Cue * Aversiveness | 0.371 | | | | | | 7.345 | | <0.001*** | |
|  |  |  | Stimuli *Cue * Trial | 0.003 | | | | | | 1.040 | | 0.298 | |
|  |  |  | Stimuli * Aversiveness * Trial | 0.006 | | | | | | 2.085 | | 0.037* | |
|  |  |  | Cue * Aversiveness * Trial | -0.004 | | | | | | -1.767 | | 0.077 | |
|  |  |  | Stimuli * Cue * Aversiveness * Trial | -0.001 | | | | | | -0.389 | | 0.697 | |
|  | Valence | Aversiveness (Heat and Salt vs. Sugar)^e^ | Stimuli | | | -1.662 | | | -9.459 | | | <0.001*** | |
|  |  |  | Cue | | | -0.192 | | | -2.789 | | | 0.007** | |
|  |  |  | Aversiveness | | | -2.591 | | | -7.370 | | | <0.001*** | |
|  |  |  | Trial | | | -0.036 | | | -4.888 | | | <0.001*** | |
|  |  |  | Stimuli * Cue | | | -0.741 | | | -9.701 | | | <0.001*** | |
|  |  |  | Stimuli * Aversiveness | | | -1.614 | | | -8.619 | | | <0.001*** | |
|  |  |  | Cue * Aversiveness | | | -0.091 | | | -1.244 | | | 0.218 | |
|  |  |  | Stimuli * Trial | | | -0.010 | | | -2.269 | | | 0.023* | |
|  |  |  | Cue * Trial | | | 0.0001 | | | 0.020 | | | 0.984 | |
|  |  |  | Aversiveness * Trial | | | 0.001 | | | 0.182 | | | 0.856 | |
|  |  |  | Stimuli * Cue * Aversiveness | | | -0.006 | | | -0.069 | | | 0.945 | |
|  |  |  | Stimuli *Cue * Trial | | | 0.012 | | | 2.607 | | | 0.009** | |
|  |  |  | Stimuli * Aversiveness * Trial | | | -0.0001 | | | -0.020 | | | 0.984 | |
|  |  |  | Cue * Aversiveness * Trial | | | 0.004 | | | 1.058 | | | 0.290 | |
|  |  |  | Stimuli * Cue * Aversiveness * Trial | | | -0.002 | | | -0.345 | | | 0.730 | |
| Dependent Variable | | **Group Factor** | **Factors** | **Estimate** | | | | | | **T Value** | | **P Value** | |
| Intensity | | Modality (Heat vs. Salt and Sugar)^f^ | Stimuli | 1.936 | | | | | | 16.009 | | <0.001*** | |
|  |  |  | Cue | 0.312 | | | | | | 5.210 | | <0.001*** | |
|  |  |  | Modality (Pain vs. Tastes) | -0.785 | | | | | | -3.745 | | <0.001*** | |
|  |  |  | Trial | 0.002 | | | | | | 0.723 | | 0.472 | |
|  |  |  | Stimuli * Cue | -0.093 | | | | | | -2.003 | | 0.045* | |
|  |  |  | Stimuli * Modality | 0.245 | | | | | | 1.877 | | 0.066 | |
|  |  |  | Cue * Modality | 0.072 | | | | | | 1.095 | | 0.278 | |
|  |  |  | Stimuli * Trial | 0.010 | | | | | | 3.494 | | <0.001*** | |
|  |  |  | Cue * Trial | -0.003 | | | | | | -1.387 | | 0.166 | |
|  |  |  | Modality * Trial | -0.010 | | | | | | -2.834 | | 0.006** | |
|  |  |  | Stimuli * Cue * Modality | 0.801 | | | | | | 15.897 | | <0.001*** | |
|  |  |  | Stimuli *Cue * Trial | 0.003 | | | | | | 1.115 | | 0.265 | |
|  |  |  | Stimuli * Modality * Trial | 0.002 | | | | | | 0.588 | | 0.557 | |
|  |  |  | Cue * Modality * Trial | -0.0002 | | | | | | -0.068 | | 0.946 | |
|  |  |  | Stimuli * Cue * Modality * Trial | -0.004 | | | | | | 1.338 | | 0.181 | |
| Valence | | Aversiveness (Heat and Salt vs. Sugar)^g^ | Stimuli | | | -1.707 | | | -7.007 | | | <0.001*** | |
|  |  |  | Cue | | | -0.196 | | | -2.871 | | | 0.006** | |
|  |  |  | Modality (Pain vs. Tastes) | | | -0.323 | | | -0.661 | | | 0.511 | |
|  |  |  | Trial | | | -0.034 | | | -5.055 | | | <0.001*** | |
|  |  |  | Stimuli * Cue | | | -0.786 | | | -10.366 | | | <0.001*** | |
|  |  |  | Stimuli * Modality | | | -0.941 | | | -3.588 | | | <0.001*** | |
|  |  |  | Cue * Modality | | | -0.115 | | | -1.543 | | | 0.128 | |
|  |  |  | Stimuli * Trial | | | -0.010 | | | -2.167 | | | 0.030* | |
|  |  |  | Cue * Trial | | | 0.0001 | | | 0.049 | | | 0.961 | |
|  |  |  | Modality * Trial | | | 0.025 | | | 3.393 | | | 0.001** | |
|  |  |  | Stimuli * Cue * Modality | | | -0.614 | | | -7.419 | | | <0.001*** | |
|  |  |  | Stimuli *Cue * Trial | | | 0.012 | | | 2.537 | | | 0.011** | |
|  |  |  | Stimuli * Modality * Trial | | | -0.0001 | | | -0.020 | | | 0.984 | |
|  |  |  | Cue * Modality * Trial | | | 0.0003 | | | 0.079 | | | 0.937 | |
|  |  |  | Stimuli * Cue * Modality * Trial | | | -1.707 | | | -7.007 | | | <0.001*** | |

^a.^ This table reports results of linear mixed models across all trials. *** = p <0.001, ** = p < 0.01, * = p < 0.05

^b^. The winning model of Group for intensity ratings: Intensity ~ Group_PainvsSalt * Cue * Stimuli * Trial + ~ Group_SaltvsSugar * Cue * Stimuli * Trial + (1 + Stimuli * Cue|Subject)

^c^. The winning model of Group for valence ratings: Valence ~ Group_PainvsSalt * Cue * Stimuli * Trial + ~ Group_SaltvsSugar * Cue * Stimuli * Trial + (1 + Stimuli * Cue|Subject)

^d^. The winning model of Group for intensity ratings: Intensity ~ Aversiveness * Cue * Stimuli * Trial + (1 + Stimuli + Cue + Trial |Subject)

^e^. The winning model of Group for valence ratings: Valence ~ Aversiveness * Cue * Stimuli * Trial + (1 + Stimuli + Cue + Trial |Subject)

^f^. The winning model of Group for intensity ratings: Intensity ~ Modality* Cue * Stimuli * Trial + (1 + Stimuli + Cue + Trial |Subject)

^g^. The winning model of Group for valence ratings: Valence ~ Modality * cue * stimuli * trial + (1 + stimuli + cue + trial |subject)
