## Supplementary material for "Domain-general neural effects of associative learning and expectations on pain and hedonic taste perception": Table S3

**Table S3-1.** Results of the winning Group models for intensity ratings and valence ratings in medium trials. *** 0.001, ** 0.01, * 0.05

| The wining model: Intensity ~ Group_PainvSalt * Trial + Group_SaltvSugar * Trial + Cue + (1 + Cue + Trial \|Subject) | | | | | |
| --- | --- | --- | --- | --- | --- |
|  | **Estimate** | | | **T Value** | **P Value** |
| Group_PainvsSalt | -1.146 | | | -3.764 | <0.001*** |
| Trial | 0.002 | | | 0.600 | 0.551 |
| Group_SaltvsSugar | 0.036 | | | 0.119 | 0.906 |
| Cue | 0.305 | | | 5.140 | <0.001*** |
| Group_PainvsSalt * Trial | -0.015 | | | -2.695 | 0.009** |
| Group_SaltvsSugar * Trial | 0.0001 | | | 0.011 | 0.991 |
| The wining model: Valence ~ Group_PainvSalt * Trial + Group_SaltvSugar * Trial + Cue + (1 + Cue + Trial \|Subject) | | | | | |
|  | | **Estimate** | **T Value** | | **P Value** |
| Group_PainvsSalt | | -0.373 | -0.661 | | 0.511 |
| Trial | | -0.033 | -4.004 | | <0.001*** |
| Group_SaltvsSugar | | -3.461 | -6.239 | | <0.001*** |
| Cue | | -0.201 | -3.013 | | 0.004** |
| Group_PainvsSalt * Trial | | -0.035 | -2.853 | | 0.006** |
| Group_SaltvsSugar * Trial | | 0.006 | 0.543 | | 0.589 |

**Table S3-2.** Results of the winning Aversiveness models for intensity ratings and valence ratings in medium trials. *** 0.001, ** 0.01, * 0.05

| The wining model: Intensity ~ Cue + (1 + Cue + Trial \|Subject) | | | |
| --- | --- | --- | --- |
|  | **Estimate** | **T Value** | **P Value** |
| Cue | 0.301 | 5.087 | <0.001*** |
| The wining model: Valence ~ Aversiveness + Trial + Cue + (1 + Cue + Trial \|Subject) | | | |
|  | **Estimate** | **T Value** | **P Value** |
| Aversiveness | -2.654 | -6.079 | <0.001*** |
| Trial | -0.036 | -4.082 | <0.001*** |
| Cue | -0.200 | -3.009 | 0.004** |

**Table S3-3.** Results of the winning Modality models for intensity ratings and valence ratings in medium trials. *** 0.001, ** 0.01, * 0.05

| The wining model: Intensity ~ Modality + Trial + Cue + (1 + Cue + Trial \|Subject) | | | |
| --- | --- | --- | --- |
|  | **Estimate** | **T Value** | **P Value** |
| Modality (Pain vs. Tastes) | -0.866 | -3.647 | <0.001*** |
| Trial | 0.003 | 0.653 | 0.517 |
| Cue | 0.305 | 5.137 | <0.001*** |
| Modality * Trial | -0.011 | -2.678 | 0.010** |
| The wining model: Valence ~ Trial + Cue + (1 + Cue + Trial \|Subject) | | | |
|  | **Estimate** | **T Value** | **P Value** |
| Trial | -0.036 | -4.106 | <0.001*** |
| Cue | -0.199 | -2.987 | 0.004** |
