## Supplementary material for "Domain-general neural effects of associative learning and expectations on pain and hedonic taste perception": Table S4

**Table S4-1.** Results of the winning Group models for expected intensity and valence ratings in medium trials. *** 0.001, ** 0.01, * 0.05

| The wining model: Expected Intensity ~ Group_PainvSalt * Cue + Group_SaltvSugar * Cue +Block + (1 + 1 \|Subject) | | | | | |
| --- | --- | --- | --- | --- | --- |
|  | **Estimate** | | | **T Value** | **P Value** |
| Group_PainvsSalt | -0.349 | | | -1.643 | 0.106 |
| Cue | 1.312 | | | 20.028 | <0.001*** |
| Group_SaltvsSugar | 0.516 | | | 2.447 | 0.018* |
| Block | 0.079 | | | 1.701 | 0.090 |
| Group_PainvsSalt * Cue | 0.612 | | | 6.499 | <0.001*** |
| Group_SaltvsSugar * Cue | 0.499 | | | 5.419 | <0.001*** |
| The wining model: Expected Valence ~ Group_PainvSalt * Cue * Block + Group_SaltvSugar * Cue * Block + (1 + 1 \|Subject) | | | | | |
|  | | **Estimate** | **T Value** | | **P Value** |
| Group_PainvsSalt | | -1.021 | -2.412 | | 0.019* |
| Cue | | -1.187 | -11.642 | | <0.001*** |
| Block | | -0.299 | -4.115 | | <0.001*** |
| Group_SaltvsSugar | | -3.581 | -8.513 | | <0.001*** |
| Group_PainvsSalt * Cue | | -1.157 | -7.884 | | <0.001*** |
| Group_PainvsSalt * Block | | 0.365 | 3.489 | | <0.001*** |
| Cue * Block | | -0.082 | -1.131 | | 0.259 |
| Group_SaltvsSugar * Cue | | -1.722 | -12.000 | | <0.001*** |
| Group_SaltvsSugar * Block | | -0.092 | -0.900 | | 0.368 |
| Group_PainvsSalt * Cue * Block | | -0.047 | -0.452 | | 0.651 |
| Group_SaltvsSugar * Cue * Block | | -0.002 | -0.024 | | 0.981 |

**Table S4-2.** Results of the winning Aversiveness models for expected intensity and valence ratings in medium trials. *** 0.001, ** 0.01, * 0.05

| The wining model: Expected Intensity ~ Aversiveness* Cue + Block + (1 + 1\|Subject) | | | |
| --- | --- | --- | --- |
|  | **Estimate** | **T Value** | **P Value** |
| Aversiveness | -0.269 | -1.516 | 0.135 |
| Cue | 1.466 | 20.645 | <0.001*** |
| Block | 0.079 | 1.692 | 0.091 |
| Aversiveness * Cue | 0.458 | 6.446 | <0.001*** |
| The wining model: Expected Valence ~ Aversiveness *Cue * Block+ (1 + 1 \|Subject) | | | |
|  | **Estimate** | **T Value** | **P Value** |
| Aversiveness | -2.223 | -3.629 | <0.001*** |
| Cue | 2.304 | 4.111 | <0.001*** |
| Block | -0.321 | -2.675 | 0.008** |
| Aversiveness * Cue | 1.354 | 2.417 | 0.016* |
| Aversiveness * Block | 0.211 | 1.761 | 0.079 |
| Cue * Block | 0.222 | 1.314 | 0.190 |
| Aversiveness * Cue * Block | 0.120 | 0.712 | 0.477 |

**Table S4-3.** Results of the winning Modality models for expected intensity and valence ratings in medium trials. *** 0.001, ** 0.01, * 0.05

| The wining model: Expected Intensity ~ Modality * Cue + Block + (1 + 1\|Subject) | | | |
| --- | --- | --- | --- |
|  | **Estimate** | **T Value** | **P Value** |
| Modality (Pain vs. Tastes) | 0.399 | 2.321 | 0.024* |
| Cue | 1.176 | 16.707 | <0.001*** |
| Block | 0.079 | 1.669 | 0.096 |
| Modality * Cue | 0.364 | 5.167 | <0.001*** |
| The wining model: Expected Valence ~ Modality * Cue + Block + (1 +1\|Subject) | | | |
|  | **Estimate** | **T Value** | **P Value** |
| Modality (Pain vs. Tastes) | -2.700 | -8.294 | <0.001*** |
| Cue | -0.748 | -6.798 | <0.001*** |
| Block | -0.315 | -4.254 | <0.001*** |
| Modality * Cue | -1.282 | -11.658 | <0.001*** |
