## Supplementary material for "Domain-general neural effects of associative learning and expectations on pain and hedonic taste perception": Table S5

**Table S5.** Results of mass-univariate functional analyses regarding stimulation level for Heat, Salt, and Sugar in the MNI space. Results are reported under *p* < 0.001 with 10 continuous voxels, uncorrected in the whole brain (See corrected results in Table 1). BA = Brodmann area, L = left hemisphere, mid = middle, R = right hemisphere, SMA = Supplementary Motor Area, OFC = Orbital Frontal Cortex, IFG = Inferior Frontal Gyrus.

| Region label | | BA | | Cluster size | | *x* | *y* | *z* | *t*-value |
| --- | --- | --- | --- | --- | --- | --- | --- | --- | --- |
|  | **Heat: High - Low** | | | | | | | | |
| Temporal Pole (R) | | 44  10  31  7  17  45 | | 494 | 52 | | 16 | -4 | 5.23 |
| Mid Frontal Gyrus (L) | |  |  | 180 | -34 | | 40 | 28 | 4.42 |
| Precuneus (L) | |  |  | 226 | -16 | | -44 | 46 | 4.63 |
| Precuneus (R) | |  |  | 184 | 4 | | -80 | 52 | 4.67 |
| Calcarine (R) | |  |  | 183 | 2 | | -74 | 10 | 4.78 |
| IFG (L) | |  |  | 24 | -38 | | 22 | 10 | 4.45 |
|  | **Salt: High - Low** | | | | | | | | |
| SMA (R) | | 6  8  24  24  6  50  18  10  47  21  38  40  49  11  49  17  50  48 | | 128 | 4 | | 10 | 62 | 5.12 |
| SMA (L) | |  |  |  | -4 | | 26 | 58 | 5.04 |
| Superior Frontal Gyrus (L) | |  |  | 137 | -2 | | -8 | 44 | 5.04 |
| Mid Cingulum (L) | |  |  | 137 | -2 | | -8 | 44 | 5.04 |
| Precentral Gyrus (R) | |  |  | 11 | 76 | | 8 | 20 | 5 |
| Thalamus (L) | |  |  | 26 | -4 | | -20 | 10 | 4.8 |
| Cerebellum (R) | |  |  | 66 | 14 | | -68 | -16 | 4.78 |
| Mid Frontal Gyrus (L) | |  |  | 82 | -38 | | 50 | 20 | 4.67 |
| OFC (R) | |  |  | 19 | 46 | | 28 | -8 | 4.65 |
| Mid Temporal Gyrus (R) | |  |  | 24 | 80 | | -14 | -14 | 4.61 |
| Temporal Pole (L) | |  |  | 73 | -50 | | 26 | -22 | 4.61 |
| Rolandic Operculum (L) | |  |  | 56 | -40 | | -32 | 22 | 4.55 |
| Amygdala (R) | |  |  | 56 | 26 | | 2 | -10 | 4.47 |
| OFC (L) | |  |  | 49 | -10 | | 68 | -20 | 4.28 |
| Putamen (R) | |  |  |  | 26 | | -2 | 4 | 4.44 |
| Calcarine (R) | |  |  | 42 | 8 | | -92 | 2 | 4.25 |
| Thalamus (R) | |  |  |  | 4 | | -26 | 8 | 4.23 |
| Caudate (L) | |  |  | 13 | -14 | | 4 | 10 | 4.22 |
| Putamen (L) | | 49  53  18  39  32 | | 16 | -22 | | 2 | 8 | 4.18 |
| Amygdala (L) | |  |  | 22 | -26 | | -2 | -14 | 4.15 |
| Lingual Gyrus (R) | |  |  | 13 | 10 | | -80 | -4 | 4.04 |
| Supramarginal Gyrus (L) | |  |  | 14 | -58 | | -44 | 34 | 4 |
| Anterior Cingulum (R) | |  |  | 13 | 2 | | 44 | 4 | 3.95 |
|  | **Sugar: High - Low** | | | | | | | | |
| Cerebellum (R) | | 18  6  4  4  1 | | 21 | 8 | | -76 | -46 | 4.79 |
| Precentral Gyrus (L) | |  |  | 37 | -58 | | -2 | 32 | 4.36 |
| Rolandic Operculum (R) | |  |  | 10 | 52 | | -4 | 22 | 4.29 |
| Postcentral gyrus (L) | |  |  |  | -56 | | -10 | 40 | 4.23 |
| Postcentral Gyrus (R) | |  |  | 21 | 62 | | -14 | 34 | 3.99 |
| Heat - Salt: High - Low | | | | | | | | | |
| Postcentral Gyrus (R) | | 4  40  1  6  6  37  50  48 | | 27 | 32 | | -29 | 59 | 4.95 |
| Supramarginal Gyrus (R) | |  |  | 42 | 59 | | -32 | 29 | 4.75 |
| Rolandic Operculum (R) | |  |  |  | 53 | | -23 | 23 | 3.82 |
| SMA (R) | |  |  | 25 | 5 | | -14 | 62 | 4.57 |
| Mid Cingulum (R) | |  |  |  | 5 | | -14 | 53 | 4.3 |
| Cerebellum (L) | |  |  | 49 | -38 | | -53 | -32 | 4.47 |
| Thalamus (R) | |  |  | 11 | 2 | | -11 | 14 | 4.04 |
| Caudate (R) | |  |  | 11 | 14 | | 26 | 11 | 3.77 |
| Salt - Heat: High - Low | | | | | | | | | |
| Postcentral Gyrus (R) | |  | 4 | 25 | 56 | | -8 | 32 | 4.09 |
| Salt - Sugar: High - Low | | | | | | | | | |
| Rectus (L) | | 10  10  40  40  39  10  6  10  31  23  10 | | 15 | -5 | | 65 | -17 | 4.75 |
| OFC (L) | |  |  |  | -17 | | 68 | -11 | 4.06 |
| Rolandic Operculum (L) | |  |  | 11 | -41 | | -32 | 23 | 4.69 |
| Superior Temporal Gyrus (L) | |  |  |  | -44 | | -29 | 14 | 3.51 |
| Mid Temporal Gyrus (L) | |  |  | 14 | -50 | | -56 | 23 | 4.52 |
| Superior Frontal Gyrus (L) | |  |  | 11 | -17 | | 56 | 29 | 4.5 |
| SMA (L) | |  |  | 11 | -8 | | 17 | 62 | 4.04 |
| Mid Frontal Gyrus (L) | |  |  | 14 | -8 | | 65 | 23 | 4.02 |
| Precuneus (L) | |  |  | 16 | -5 | | -47 | 41 | 3.97 |
| Mid Cingulum (R) | |  |  |  | 2 | | -50 | 35 | 3.6 |
| OFC (L) | |  |  | 10 | -38 | | 83 | -29 | 3.88 |
| Conjunction: Heat (High - Low) & Salt (High - Low) | | | | | | | | | |
| Cerebellum (L) | | 37  24  24  8  18  6  39  44  44  10  44  13 | | 75 | -23 | | -59 | -20 | 5.21 |
| Mid Cingulum (L) | |  |  | 85 | -2 | | -8 | 44 | 4.83 |
| Anterior Cingulum (R) | |  |  |  | 2 | | 20 | 26 | 3.72 |
| SMA (L) | |  |  | 26 | -2 | | 26 | 59 | 4.18 |
| Cerebellum (R) | |  |  | 20 | 17 | | -68 | -17 | 4.09 |
| SMA (R) | |  |  | 10 | 8 | | 5 | 50 | 4 |
| Supramarginal Gyrus (L) | |  |  | 13 | -59 | | -44 | 35 | 4 |
| Temporal Pole (L) | |  |  | 13 | -59 | | 11 | -2 | 3.96 |
| Rolandic Operculum (L) | |  |  |  | -59 | | 5 | 5 | 3.6 |
| Mid Frontal Gyrus (L) | |  |  | 25 | -35 | | 53 | 20 | 3.89 |
| Temporal Pole (R) | |  |  | 16 | 59 | | 8 | 2 | 3.79 |
| Insula (R) | |  |  |  | 41 | | 14 | -2 | 3.47 |
| Conjunction: Salt (High - Low) & Sugar (High - Low) | | | | | | | | | |
| Postcentral Gyrus (L) | | 4 | | 30 | -56 | | -8 | 41 | 4.07 |
| Conjunction: Heat (High - Low) & Salt (High - Low) & Sugar (High - Low) | | | | | | | | | |
| Cerebellum (L) | | 19 | | 2 | -17 | | -59 | -17 | 3.38 |
