## Supplementary material for "Domain-general neural effects of associative learning and expectations on pain and hedonic taste perception": Table S6

**Table S6.** Results of Multivariate Pattern Analysis (MVPA) regarding stimulation level for Heat, Salt, and Sugar in the MNI space. Region names are labeled with the AAL atlas. Results are thresholded with Small Volume Correction (SVC; p < 0.05, FWE correction) in the placebo-pain mask (Atlas & Wager, 2014), or only cluster-wise FWE correction in the whole brain, or p < 0.001 with 10 continuous voxels, uncorrected in the whole brain. Only Heat Group survived SVC with the placebo-pain mask and showed significant activations after clustered-wise FWE correction. BA = Brodmann area, L = left hemisphere, R = right hemisphere.

| Region label | BA | Cluster size | *x* | *y* | *z* | *t*-value |
| --- | --- | --- | --- | --- | --- | --- |
| Heat: High - Low (SVC in the placebo-pain mask) | | | | | | |
| Rolandic_Oper_L | 44 | 1 | -44 | 8 | 16 | 10.69 |
| Frontal_Sup_R | 10 | 1 | 26 | 50 | 10 | 9.8 |
| Frontal_Inf_Tri_L | 44 | 1 | -44 | 22 | 20 | 9.04 |
| Frontal_Inf_Oper_L | 45 | 1 | -46 | 20 | 14 | 8.88 |
| Insula_L | 45 | 2 | -34 | 26 | 8 | 8.34 |
| Frontal_Mid_R | 10 | 1 | 26 | 50 | 2 | 8.3 |
| Frontal_Inf_Orb_R | 47 | 1 | 48 | 44 | -2 | 8 |
| Heat: High - Low (whole-brain activations survives FWE correction in addition to SVC) | | | | | | |
| Frontal_Inf_Oper_R | 44 | 158 | 62 | 14 | 8 | 16.1 |
| Rolandic_Oper_R | 6 |  | 56 | 4 | 10 | 11.49 |
| Frontal_Inf_Tri_R | 9 |  | 44 | 26 | 26 | 9.92 |
| SupraMarginal_R | 40 | 126 | 64 | -32 | 32 | 10.63 |
| Putamen_L | 49 | 1 | -26 | 8 | 4 | 10.59 |
| Frontal_Mid_L | 10 | 67 | -50 | 52 | 22 | 9.92 |
| Temporal_Pole_Sup_R | 22 | 4 | 58 | 4 | -8 | 9.12 |
| Supp_Motor_Area_R | 6 | 1 | 14 | 16 | 62 | 8.69 |
| Temporal_Mid_R | 39 | 1 | 64 | -46 | 10 | 8.58 |
| Caudate_R | 48 | 1 | 22 | 22 | 8 | 8.34 |
| Precentral_R | 6 | 1 | 50 | 4 | 46 | 8.3 |
| Insula_R | 45 | 1 | 26 | 34 | 14 | 8.27 |
| Temporal_Sup_R | 39 | 1 | 70 | -44 | 20 | 8.06 |
| Postcentral_R | 4 | 1 | 70 | -2 | 26 | 8.06 |
| Precentral_L | 6 | 2 | -34 | 2 | 58 | 7.96 |
| Heat: High - Low (whole-brain activations survives 10 voxels uncorrected in addition to FWE) | | | | | | |
| Temporal_Inf_L | 20 | 15 | -50 | -28 | -26 | 5 |
| Cerebelum_Crus1_L | 19 | 31 | -40 | -86 | -20 | 4.75 |
| Cerebelum_4_5_R | 18 | 14 | 8 | -52 | -2 | 4.57 |
| Lingual_R | 19 |  | 14 | -62 | -8 | 3.93 |
| Cerebelum_Crus2_R | 37 | 18 | 44 | -64 | -38 | 4.37 |
| Cerebelum_Crus1_L | 37 | 12 | -58 | -50 | -38 | 4.16 |
| Salt: High - Low (whole-brain activations survives 10 voxels uncorrected) | | | | | | |
| Occipital_Mid_L | 18 | 400 | -34 | -80 | 10 | 7.63 |
| Temporal_Mid_R_L | 39 | 59 | 68 | -56 | 16 | 6.47 |
| Precentral_R | 6 | 40 | 64 | 8 | 28 | 6.13 |
| SupraMarginal_R | 40 | 24 | 76 | -16 | 34 | 6.02 |
| Frontal_Mid_R | 9 | 84 | 52 | 26 | 40 | 5.9 |
| Vermis_6 | 18 | 64 | 2 | -68 | -10 | 5.49 |
| Supp_Motor_Area_R | 6 | 35 | 10 | 16 | 56 | 5.42 |
| Supp_Motor_Area_L | 8 |  | 2 | 14 | 46 | 4.69 |
| Frontal_Mid_Orb_R | 10 | 11 | 50 | 56 | -20 | 5.22 |
| Cuneus_R | 18 | 10 | 16 | -98 | 8 | 5.22 |
| Putamen_R | 49 | 10 | 20 | 20 | -8 | 5.19 |
| Insula_R | 13 |  | 28 | 22 | -4 | 4.95 |
| Frontal_Inf_Orb_L | 10 | 20 | -52 | 52 | -22 | 4.69 |
| Occipital_Sup_L | 19 | 18 | -26 | -98 | 26 | 4.19 |
| Temporal_Pole_Sup_R | 47 | 10 | 58 | 32 | -22 | 4.52 |
| Amygdala_L | 53 | 13 | -26 | -4 | -16 | 4.51 |
| ParaHippocampal_L | 53 |  | -26 | -4 | -26 | 4.24 |
| Frontal_Sup_L | 10 | 12 | -26 | 74 | -2 | 4.2 |
| Sugar: High - Low (whole-brain activations survives 10 voxels uncorrected) | | | | | | |
| Frontal_Inf_Oper_R | 44 | 59 | 44 | 10 | 26 | 6.61 |
| Precentral_R | 6 |  | 50 | 4 | 28 | 5.83 |
| Frontal_Inf_Tri_R | 44 |  | 40 | 22 | 22 | 4.69 |
| Temporal_Mid_R | 37 | 12 | 52 | -50 | 10 | 5.83 |
| ParaHippocampal_L | 36 | 10 | -2 | -4 | -34 | 4.75 |
| Postcentral_L | 1 | 10 | -52 | -20 | 20 | 4.75 |
| Rolandic_Oper_L | 1 |  | -46 | -16 | 26 | 4.75 |
