## Supplementary material for "Domain-general neural effects of associative learning and expectations on pain and hedonic taste perception": Table S7

**Table S7.** Single-trial mediation and moderation results on subjective intensity and valence. Each table represents results from one mediation/moderation model. Significant activations were presented for each path. For complete transparency, we have included results that passed both False Discovery Rate (FDR) corrections or survived an uncorrected threshold of *p* < 0.001 across a minimum of three contiguous voxels. Clusters were defined by contiguity with additional voxels at thresholds of *p* < 0.005 and P < 0.01. See FDR-corrected results for models without moderators for intensity and valence in Table 3. OFC: Orbital Frontal Cortex; IFG: Inferior Frontal Gyrus; DMPFC: Dorsomedial Prefrontal Cortex; RSC: Retrosplenial Cortex; DLPFC: Dorsolateral Prefrontal Cortex; M1: Primary Motor Cortex; IPL: Inferior Parietal Lobule; S1: Primary Somatosensory Cortex; PAG: Periaqueductal Gray; dACC: Dorsal Anterior Cingulate Cortex; SPL: Superior Parietal Lobule; dpIns: Dorsal Posterior Insula; TPJ: Temporoparietal junction; SMA: Supplementary Motor Area; VLPFC: Ventrolateral Prefrontal Cortex; PFC: Prefrontal Cortex.

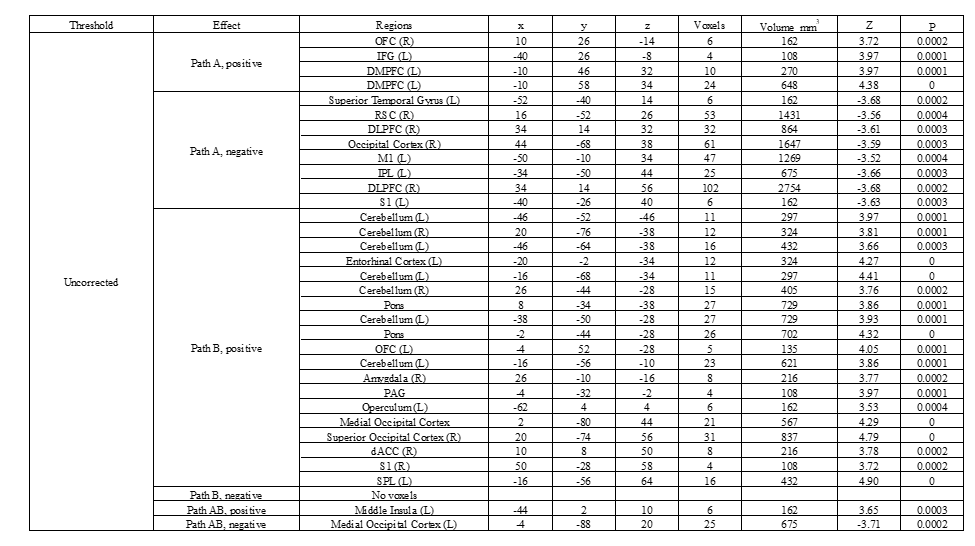
**Table S7-1**. Intensity Mediation Analysis without Moderators

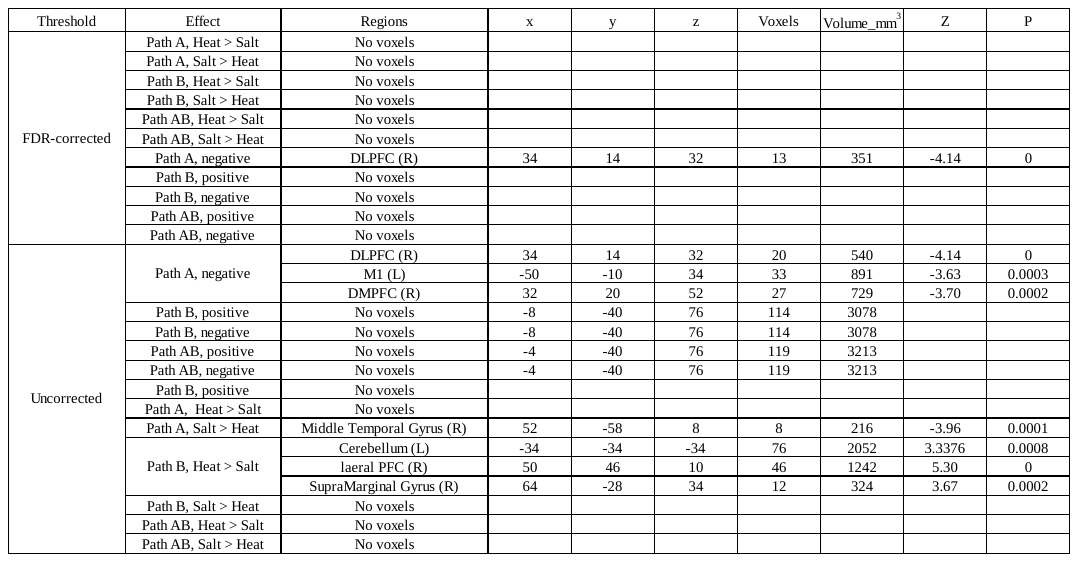
**Table S7-2**. Intensity Mediation Analysis with Heat vs. Salt as a Moderator

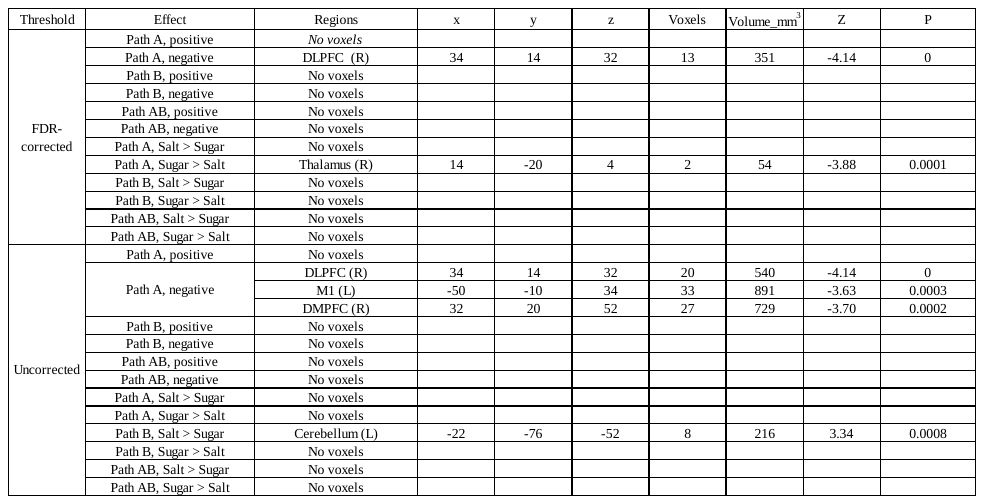
**Table S7-3**. Intensity Mediation Analysis with Salt vs. Sugar as a Moderator

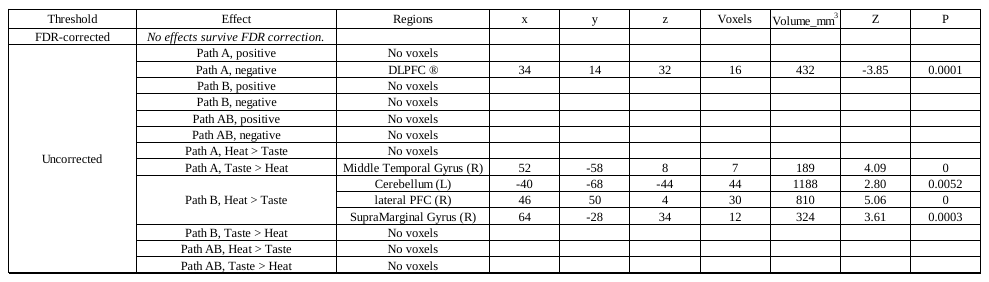
**Table S7-4**. Intensity Mediation Analysis with Modality (Heat vs. Taste) as a Moderator

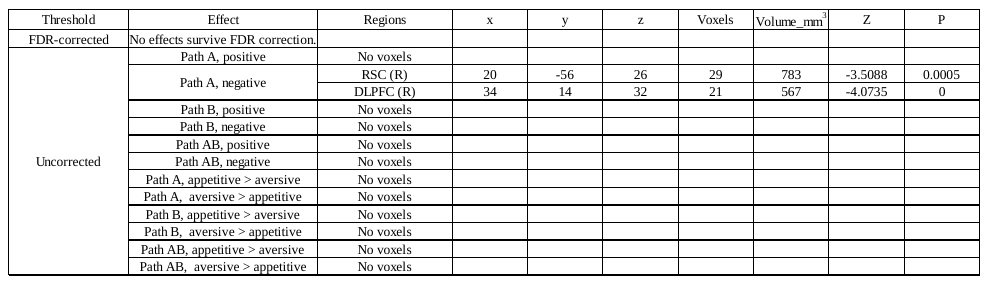
**Table S7-5**. Intensity Mediation Analysis with Aversiveness (Appetitive vs. Aversive) as a Moderator

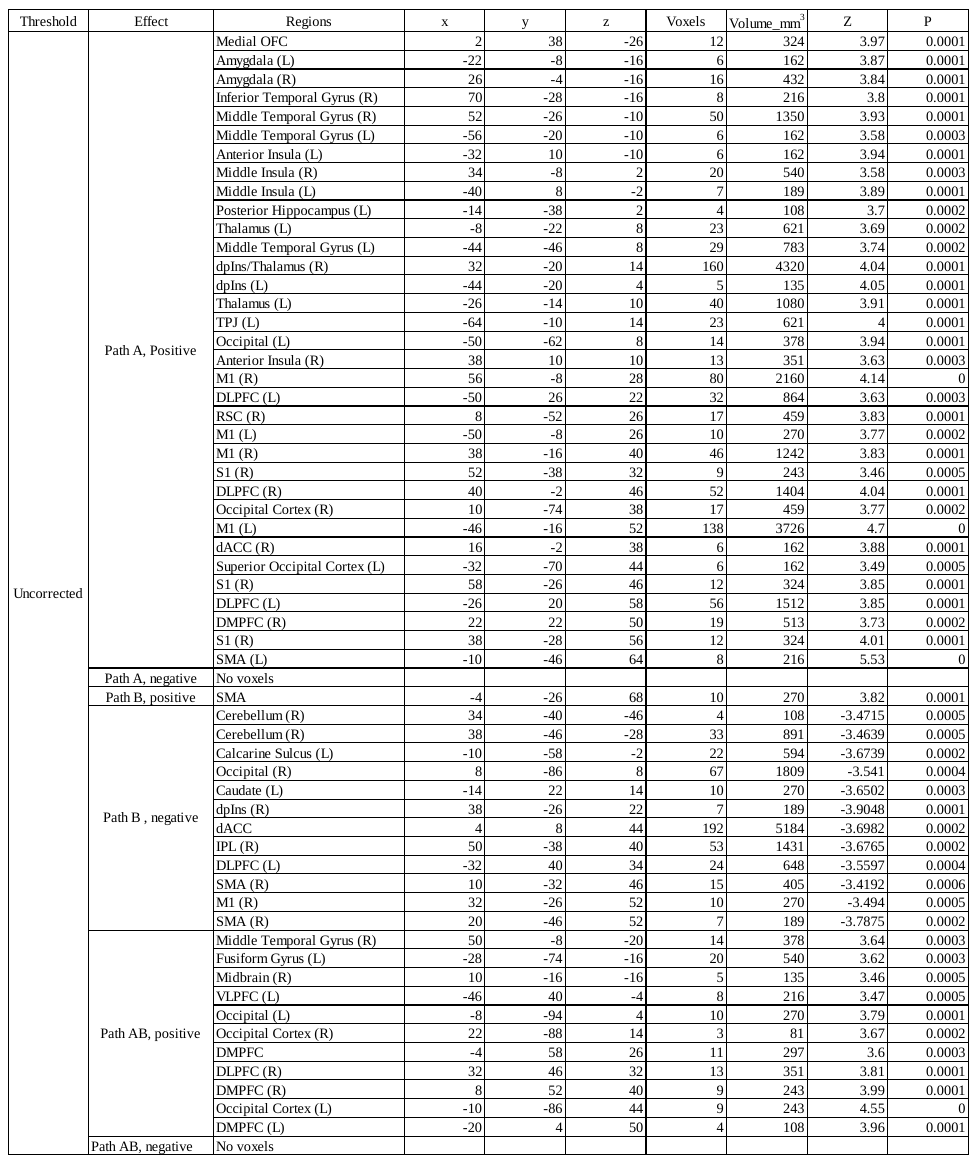
 **Table S7-6**. Valence Mediation Analysis without moderators

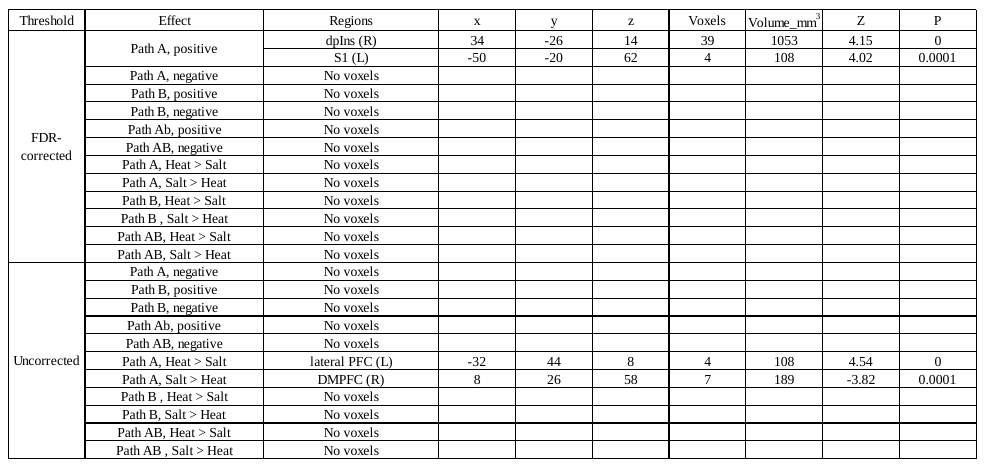
**Table S7-7**. Valence Mediation Analysis with Heat vs. Salt as a Modulator

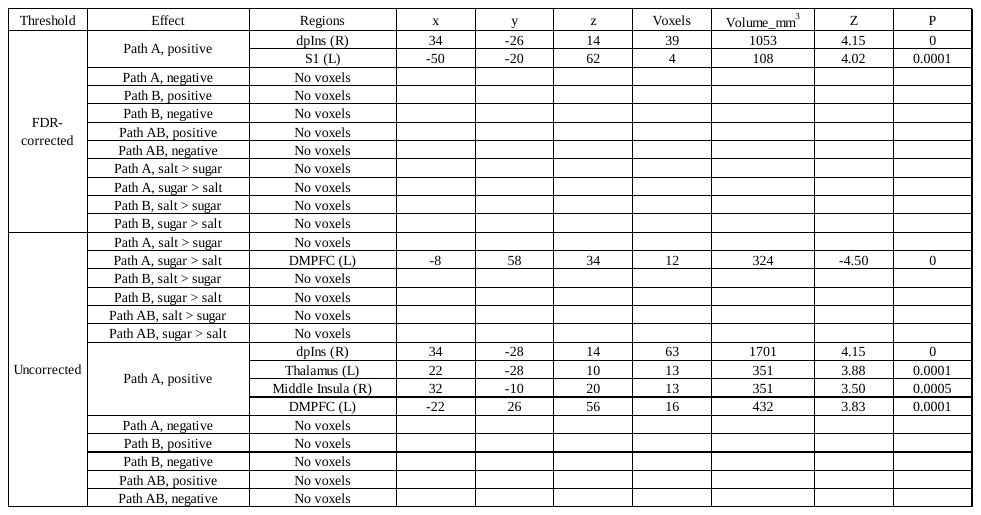
**Table S7-8**. Valence Mediation Analysis with Salt vs. Sugar as a Moderator

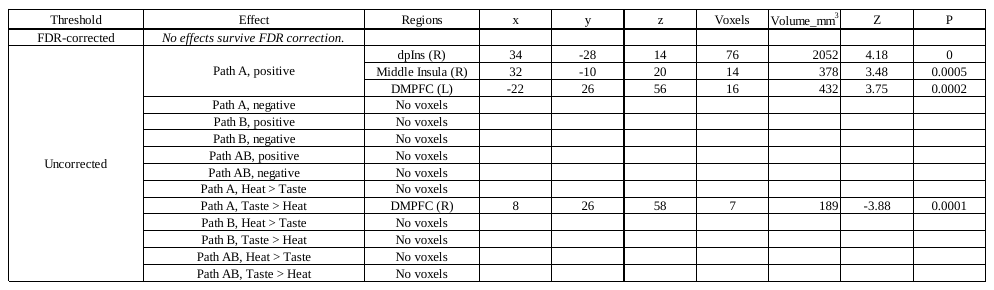
**Table S7-9**. Valence Mediation Analysis with Modality (Heat vs. Taste) as a Moderator

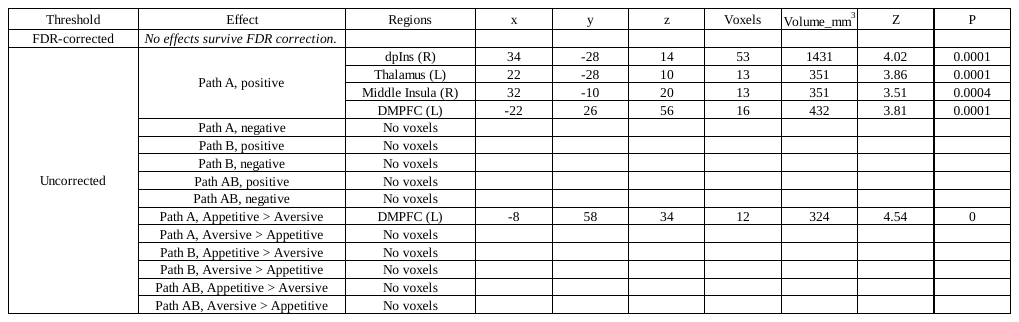
**Table S7-10**. Valence Mediation Analysis with Aversiveness (Appetitive vs. Aversive) as a Moderator
