## Supplementary material for "Domain-general neural effects of associative learning and expectations on pain and hedonic taste perception": Figure S1

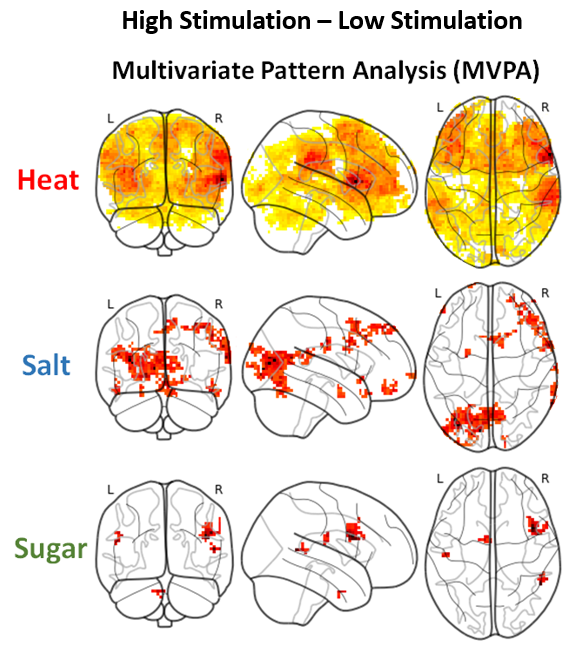

**Figure S1**. MVPA neural activations in response to Heat, Salt, and Sugar stimuli. The Heat group exhibited the strongest and most widespread activations, particularly in the bilateral inferior frontal gyrus (IFG), right Rolandic operculum, right supramarginal gyrus, left putamen, bilateral cerebellum, and left temporal gyrus. The Salt group showed prominent activations in the bilateral middle and superior frontal gyri, right precentral gyrus, left amygdala, right putamen, right supramarginal gyrus, bilateral cerebellum, and left visual cortex. The Sugar group demonstrated activations in the bilateral IFG, right precentral gyrus, right temporal cortex, left postcentral gyrus, right middle frontal gyrus, left cerebellum, and left visual cortex.
