## Supplementary material for "Domain-general neural effects of associative learning and expectations on pain and hedonic taste perception": Figure S2

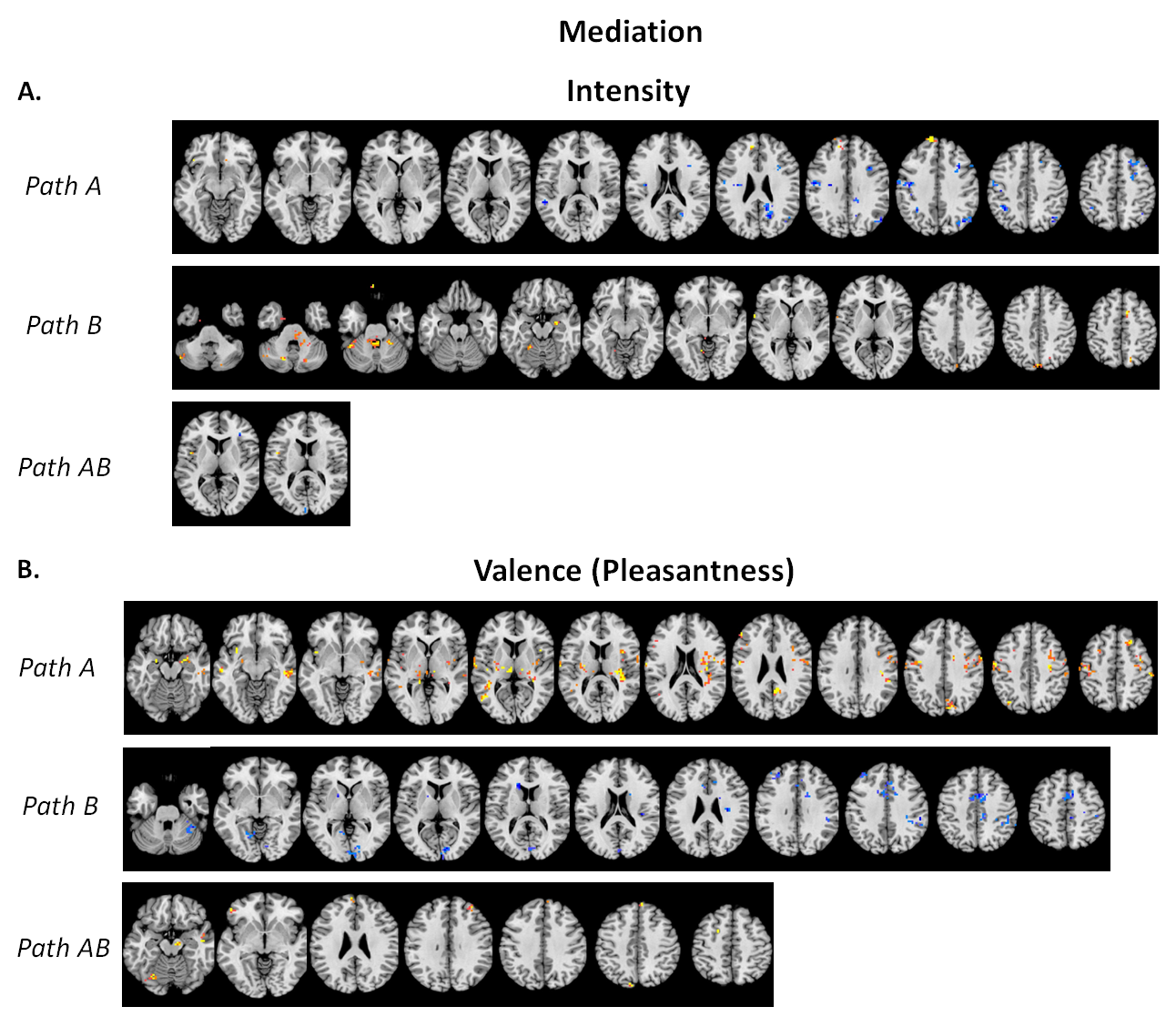

**Figure S2.** Single-Trial Mediation Effects of each path for Intensity and Valence. (A) Intensity. For Path A (top-left panel), significant activations were observed in the orbitofrontal cortex (OFC), inferior frontal gyrus (IFG), DMPFC, dorsolateral prefrontal cortex (DLPFC), postcentral gyrus, and inferior parietal lobule (IPL) in response to High vs. Low levels of intensity across all groups. In Path B (mid-left panel), after controlling for cues, individual intensity ratings were associated with significant activations in the cerebellum, right hippocampus/amygdala, brainstem, left Rolandic operculum, right supplementary motor area (SMA), and left visual cortex. For Path AB (bottom-left panel), we observed positive activations in the left insula and negative activations in the left visual cortex. (B) Valence (Pleasantness). For Path A (top-left panel), significant activations were observed in the OFC, amygdala, insula, DLPFC, and DMPFC in response to High vs. Low levels of pleasantness. These activations reflected responses to Low vs. High cues for aversive stimuli (Heat and Salt), and High vs. Low cues for Sugar. In Path B (mid-left panel), after controlling for cues, individual valence ratings were associated with significant activations in the cerebellum, striatum (left pallidum and right caudate), Rolandic operculum, MCC, DLPFC, and visual cortex. For Path AB (bottom-left panel), we observed positive activations in the left frontal cortex (e.g., OFC) and the brainstem (VTA).
