## Supplement Methods for "Domain-general neural effects of associative learning and expectations on pain and hedonic taste perception"

**Day 1: Calibration Visit**

**Questionnaires**

**The questionnaires participants completed included**: The Beck Depression Inventory (Beck et al., 1996), the Behavioral Inhibition Scale/ Behavioral Activation Scale (Carver & White, 1994), the Big Five Inventory (John et al., 1991), the Ego-Resilience Questionnaire (Block & Kremen, 1996), the Emotion Regulation Questionnaire (Gross & John, 2003), the Fear of Pain Questionnaire (McNeil & Rainwater, 1998), the Intolerance of Uncertainty Scale (Freeston et al., 1994; Buhr & Dugas, 2002), the Mindful Attention Awareness Scale, Trait Version (Brown & Ryan, 2003), the Pain Catastrophizing Scale (Sullivan et al., 1995), the Revised Life Orientation Test (Segerstrom et al., 2011), the Positive and Negative Affect Scale (Watson et al., 1988), and the State-Trait Anxiety Inventory (Spielberger et al., 1971).

**Day 2: fMRI Visit**

**Eligible participants from the Calibration Visit**

Participants were subsequently randomized to a group for the fMRI Visit, with the following constraints: Participants whose ratings did not exhibit a linear correlation with stimulus temperature/concentration (R² < 0.4) in a single modality were deemed ineligible to experience that modality (n = 9: 4 ineligible for the Heat group; 3 ineligible for the Sugar group; 1 ineligible for the Salt group; 1 ineligible for the Salt and Sugar groups); participants who did not predominantly rate sucrose solutions as pleasant and saline solutions as unpleasant were deemed ineligible to be assigned to Sugar or Salt groups, respectively (n = 7: 5 ineligible for the Sugar group; 1 ineligible for the Salt group; 1 ineligible for both groups); and participants whose tolerance was too high (>50 degrees Celsius for heat, >2M for salt and sugar), too low (e.g., finding most stimuli intolerable), or whose range between low and high was too small (<4 degrees for heat) were excluded from those modalities due to an inability to deliver experimental stimuli within safe and appropriate ranges (n = 13: 3 ineligible for the Heat group; 5 ineligible for the Salt group; 2 ineligible for the Salt and Heat groups; 1 ineligible for the Salt and Sugar groups; 2 ineligible for all groups).

**Calibration during the fMRI Visit**

Participants in the Heat Group then underwent a repeat adaptive staircase calibration (Amir et al., 2021) to identify temperatures and skin sites for use during scanning. Participants in the Salt and Sugar Groups underwent an abbreviated calibration procedure using only water (equivalent to 0% concentration) and the prepared concentrations of salt and sucrose that were associated with low, medium, and high intensity ratings on Day 1 (with the caveat that sucrose solutions did not exceed 1.0M). Salt and Sugar Group participants provided ratings of both salt and sucrose, and were not informed which solution would be delivered during the scan. In contrast to the Calibration Visit, which separately measured pleasantness and unpleasantness, we asked participants to rate valence using a continuous scale ranging from -10 (extremely unpleasant) to 10 (extremely pleasant). Intensity ratings were the same as the Calibration Visit.

**Post-tasks**

After the fMRI scan, participants completed the debriefing interview. Participants who received thermal stimulation completed the McGill Pain Questionnaire (Melzack, 1987) and had the stimulation site inspected by a physician to determine that no skin damage had occurred. Participants who received saline or sucrose solutions were informed how much salt or sugar they had consumed during the procedures. In addition, they were advised on the recommended maximum daily intake of salt or sugar and advised to limit their salt or sugar intake for the rest of the day.

**HRF determination**

Because we were applying different stimuli at longer durations than a standard event-related design, we wanted to select a hemodynamic response function (HRF) that would fit responses to thermal and gustatory stimuli in relevant brain regions. To determine which HRF best fit this fMRI dataset, we extracted fMRI signal time series from three regions of interest (ROIs) for each participant (18 Heat Pain; 17 Salt; 17 Sugar): right middle insula, left middle insula, and right dorsal posterior insula (see Figure S1A). Middle insula ROIs were selected based on identification of gustatory cortex in a prior study that used liquid tastants while the right dorsal posterior insula ROI was selected based on a mega-analysis of previous studies that compared high and low intensity stimulation (Atlas & Wager, 2012peak coordinate: x = 33.32; y = -16.20; z = 11.70). We used 3dDeconvolve to convolve onsets corresponding to heat or taste delivery in each condition (low intensity; medium intensity + low cue; medium intensity + high cue; high intensity) with 5 different HRFs available in AFNI (WAV, BLOCK4, BLOCK5, SPMG1, MION). As shown in Figure S1A, the general linear model also included an intercept and linear drift for each run, as well as regressors for the following events with their respective HRFs: cue onsets (BLOCK(2,1)); swallow (GAM); rating (BLOCK(10,1)); rinse (BLOCK(4,1)). We included the flag “input1D” in the call to 3dDeconvolve to evaluate fits to extracted data for each ROI and each participant. For each HRF, group level analysis was performed using one sample t-tests on the values output from 3dDeconvolve for the following coefficient or contrast values: a) High Heat; b) High Heat > Low Heat; c) Average response across conditions. Subject-level results for the contrast High Heat > Low Heat are depicted in Figure S1B. As these contrasts are independent of our main contrast of interest (High Medium > Low Medium), this analysis does not bias our main findings. Based on visual inspection of the single subject results and group results, we decided to use the MION HRF in voxelwise general linear models and single trial analyses implemented in the main manuscript.

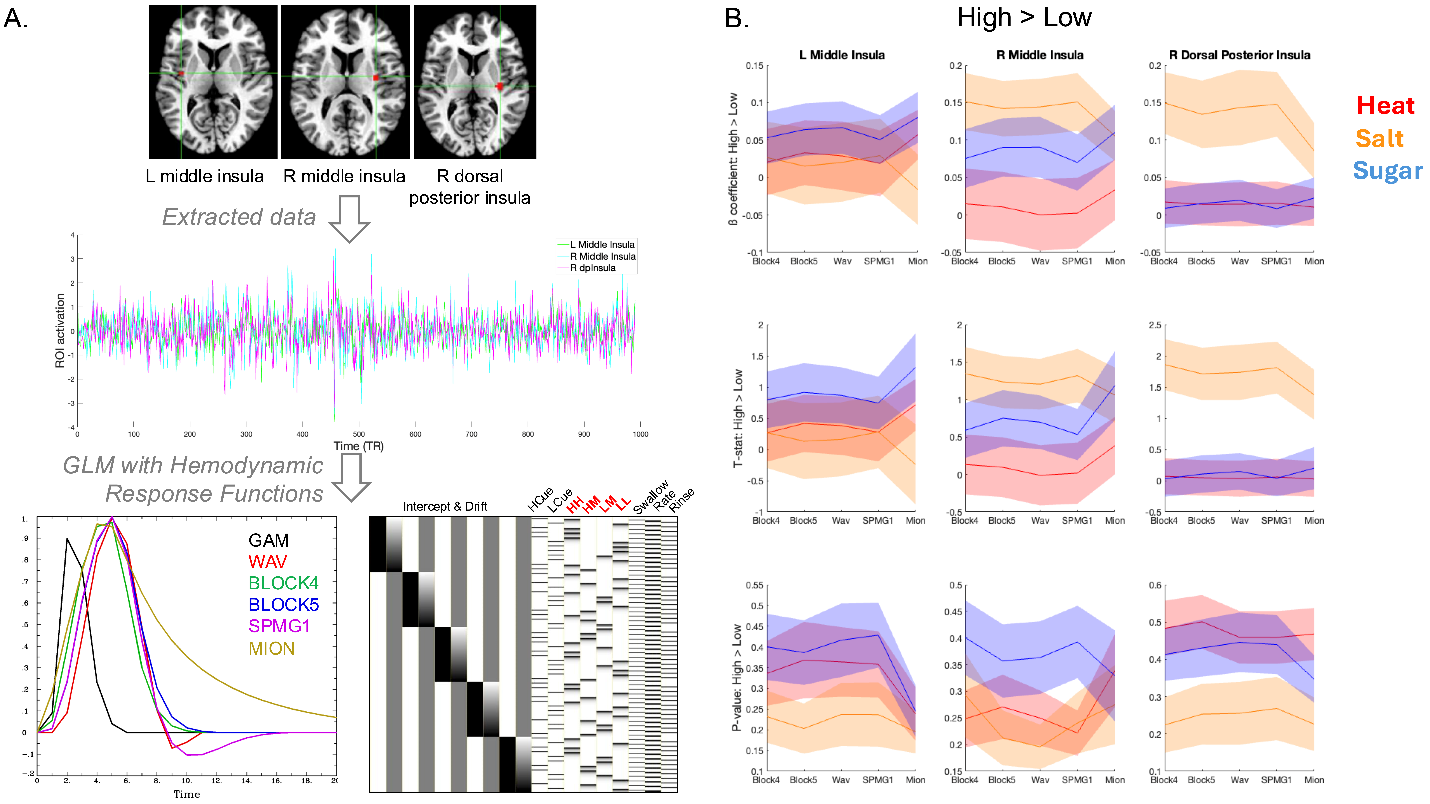

**Figure S1**. **HRF results across regions of interest.** A) *Top:* We extracted data from three regions of interest (ROIs) that have been previously identified in studies of taste (left middle insula), pain (right dorsal posterior insula), or taste and pain (right middle insula). *Middle:* For each participant, we concatenated all runs and extracted data from each ROI. Data for one participant is depicted here. *Bottom:* For each hemodynamic response function (HRF; left), we fit a general linear model (GLM) that convolved the HRF with stimulus onsets for heat or taste stimuli (red). Cue onset, swallow, and ratings were also modeled in the same GLM using impulse functions. An example GLM using MION for the stimulus onsets is presented for one participant. B) Across participants, we evaluated the impact of HRF and ROI on activation across all trials, on high intensity trials, and for the contrast [High > Low]. Here, we depict results for the [High > Low] as a function of Group (Red = Heat; Orange = Salt; Blue = Sugar), ROI (left = left middle insula; middle = right middle insula; right = right dorsal posterior insula), and HRF (x-axis). Beta-coefficients are depicted in the top row, t-statistics are presented in the middle row, and p-values are presented in the bottom row. Shaded error reflects standard error of the mean. Results were similar for other contrasts.

***Analyses of subjective intensity and valence as a function of outcome magnitude.*** We used linear mixed models to measure how changes in Stimulus Level (Low, Medium, High) impacted intensity and valence ratings. Model comparisons revealed that the best-fitting model for both intensity and valence ratings as a function of Group included all interactions among Group, Cue, Stimulus Level, and Trial as fixed effects, with the interaction of Cue and Stimulus Level per Subject as random effects. The optimal models for Aversiveness and Modality also included these interactions as fixed effects, with the intercepts of Cue and Stimulus Level per subject as random effects. All comparisons were adjusted using Bonferroni correction. See details of all model comparisons in Table S1-1.

***Analyses of cue effects on medium trials.*** Analyses of medium outcomes included the same factors as analyses across all outcomes with the exception of the Stimulus Level Factor. For intensity ratings, the wining models for the models including Group or Modality both included Cue and the interaction of Group/Modality and Trial as fixed effects, with the intercept of Cue and Trial per Subject as random effects. For the aversiveness models, the optimal model only included Cue as a fixed effect and the slope of Cue per Subject as random effects. For valence ratings, the optimal Group model mirrored the intensity model in terms of fixed and random effects. The Aversiveness model included Aversiveness, Cue, and Trial as fixed effects, with interactions of Cue and Trial per Subject as random effects. The Modality model only had Cue and Trial as fixed effects, with the interaction of Cue and Trial per Subject as random effects. See details of all model comparisons in Table S1-2.

***Expectancy ratings for High and Low cues.*** To investigate whether conscious expectations were influenced by learned cues, we analyzed separate multilevel models for expected intensity and expected valence. These included the same factors as analyses of outcome ratings, i.e., Cue, Group/Aversiveness/Modality, and time (Block). Model comparison showed that the optimal model for expected intensity contained the interaction of Group/Aversiveness/Modality and Cue, and the main effect of Block. For expected valence, the best model included the interaction of Group/Aversiveness, Cue, and Block. For model contained Modality, the best model included the interaction of Modality and Cue, and the main effect of Block. See model comparison results in Table S1-3.

**Subjective-level Multivariate pattern analysis (MVPA)**

Per preregistration, we used multivoxel pattern analysis (MVPA) techniques to assess whether the brain patterns associated with stimulus outcome levels were robust and differed across modalities. We utilized the Decoding Toolbox (TDT, https://sites.google.com/site/tdtdecodingtoolbox/) employing a support vector machine (SVM)-based searchlight method with a 5-voxel radius . Our analysis incorporated a leave-one-run-out cross-validation scheme paired with a permutation test. For this approach, the classifier was trained on data from three randomly selected runs (ensuring at least three trials per condition per run) and then tested on the remaining run. We excluded the first run from our analysis due to its distinct conditioning/learning nature, which differed from the testing-focused nature of the subsequent four runs. Due to having fewer than three trials per condition per run, two participants from the pain group were excluded from this analysis. The searchlight procedure was conducted in the MNI space for each participant without prior normalizing and smoothing. After individual-level decoding, we normalized and smoothed the results to prepare for group-level analysis.
