## Supplement Results for "Domain-general neural effects of associative learning and expectations on pain and hedonic taste perception"

**Psychophysical response functions across modalities**

We observed high correlations between temperature/concentration and subjective intensity in all conditions (see Figure 1A). A one-way ANOVA on correlation coefficients revealed a main effect of Condition (F(2,70) = 6.22, *p* = .003), driven by lower stimulus-response correlations in the Sugar condition than the Heat condition (Heat – Sugar: *p* _adjusted_ = .001). Associations between stimulus temperature/concentration and subjective intensity are reported in the main manuscript. We also observed a main effect of Condition when we examined with unpleasantness, we found that an increase of one SD temperature or concentration was associated with 1.56 units higher subjective unpleasantness (B = 1.56, *p* < .001). We also observed a main effect of Condition (F(2, 4725.5) = 1260.8, *p* < .001) and a Stimulus Level x Condition interaction (F(2,4725) = 545.8, *p* < .001), such that saline solution was rated most unpleasant (Intercept _Salt_ = 4.63; Intercept _Heat_ = 3.58; Intercept _Sugar_ = 1.08; all *p’s* < .001), but effects of stimulus level on subjective unpleasantness were strongest with heat stimuli (B _heat_ = 2.45; B _Salt_ = 1.99; B _Salt_ = 0.18; all *p’s* < .001), and pairwise differences between conditions were all significant (*p* < .001). When we examined associations with pleasantness, we found that an increase of one SD temperature or correlation was associated with a 0.08 reduction in pleasantness (B= -0.08, *p* < .01). We also observed a main effect of Condition (F(2,4722.2) = 739.78, *p* < .001) and a significant Stimulus Level x Condition Interaction (F(2, 4722) = 425.60, *p* < .001), such that participants rated sucrose as more pleasant than heat or saline (Intercept _Sugar_ = 3.46; Intercept _Salt_ = 1.00; Intercept _Heat_ = 1.93; all *p’s* < 0.001) and there were positive associations with sucrose solution intensity (B = 0.95, *p* < .001) but negative associations between changes in stimulus intensity and subjective pleasantness for heat (B = -0.93, *p* < .001) and saline solution (B = -0.19, *p* < .001).

**Moderated mediation on intensity: Path b (whole brain uncorrected)**

Path b effects varied across groups in right lateral PFC, driven by higher associations in the Heat Group than the Salt Group (Mean Heat = 0.6497, P < 0.000; Mean Salt = 0.0989, P = 0.2940; t Heat vs Salt =5.6427, P < 0.001). Follow up analyses revealed that the Heat Group showed stronger Path B effects in the right lateral PFC than the Sugar Group (Mean Sugar = 0.2349, P = 0.0536; t Heat vs Sugar = 6.0042, P < 0.001). Consistent with this, we observed moderation of Path b by Modality in the right Supramarginal Gyrus, as well as the left cerebellum. There were no voxels whose Path b effects were moderated by Salt vs Sugar, or by Aversiveness (Supplementary Tables 8). We did not observe any Path b effects when controlling for Group, Aversiveness, or Modality.
