## Supplementary material for "Domain-general neural effects of associative learning and expectations on pain and hedonic taste perception": Table S1

**Table S1-1.** Model comparison of linear regression models for all trials. For each outcome (intensity or valence ratings), our main models included the following mean-centered predictors: Group (dummy coded: Pain vs salt (“group_PainvSalt”; 1 = pain, -1 = salt) and Salt vs sugar (“group_SaltvSugar”, 1 = salt, -1 = sugar), Cue (high cue = 1, low cue = -1), Stimuli (high intensity = 1, medium intensity =0, low intensity = -1), and Trial. We also included random intercepts and slopes per Subject. In addition, we also report model comparison results for models of Aversiveness (Group: aversive versus appetitive) and Modality (Group: pain versus tastes). Factors involved were the same for all three types of models, only with different Group factor. For all models, Akaike information criterion (AIC) and Bayesian information criterion (BIC) were computed to compare models. Smaller AIC/BIC indicates better model fit, adjusting for model complexity. The models highlighted in red were the best fitting models for each type and for each rating. Note that, we only list the models that make sense and do not fail to converge.

| Regression Model | | AIC | BIC |
| --- | --- | --- | --- |
| *Three groups: Heat, Salt, and Sugar* | | | |
| *Intensity ratings* | | | |
| M1: group_PainvSalt + group_SaltvSugar + (1 + 1\|subject) | 14351 | | 14381 |
| M2: group_PainvSalt + group_SaltvSugar + cue + (1 + 1\|subject) | | 12901 | 12938 |
| M3: group_PainvSalt + group_SaltvSugar + cue + stimuli + (1 + 1\|subject) | | 11885 | 11927 |
| M4: group_PainvSalt * stimuli * cue + group_SaltvSugar * stimuli * cue + (1 + 1 \|subject) | | 11500 | 11584 |
| M5: group_PainvSalt + group_SaltvSugar + cue + trial + (1 +1\|subject) | | 12896 | 12938 |
| M6: group_PainvSalt * cue + group_SaltvSugar * cue + stimuli + trial + (1 +1\|subject) | | 11768 | 11828 |
| M7: group_PainvSalt* stimuli + group_SaltvSugar * stimuli + cue + trial + (1 + 1\|subject) | | 11711 | 11772 |
| M8: group_PainvSalt + group_SaltvSugar + cue * stim + trial + (1 + 1\|subject) | | 11863 | 11918 |
| M9: group_PainvSalt * trial + group_SaltvSugar * trial + stimuli + cue + (1 + 1 \|subject) | | 11802 | 11862 |
| M10: group_PainvSalt + group_SaltvSugar + trial * stim + cue + (1 + 1 \|subject) | | 11858 | 11913 |
| M11: group_PainvSalt + group_SaltvSugar + stimuli + cue * trial + (1 + 1 \|subject) | | 11865 | 11920 |
| M12: group_PainvSalt * cue * stimuli + group_SaltvSugar * cue * stimuli + trial + (1 + 1\|subject) | | 11487 | 11578 |
| M13: group_PainvSalt * cue * stimuli * trial + group_SaltvSugar * cue * stim* trial + (1 + 1\|subject) | | 11458 | 11615 |
| M14: group_PainvSalt * cue * stimuli * trial + group_SaltvSugar * cue * stim* trial + (1 + stimuli\|subject) | | 10925 | 11094 |
| M15: group_PainvSalt * cue * stimuli * trial + group_SaltvSugar * cue * stim* trial + (1 + trial\|subject) | | 11410 | 11580 |
| M16: group_PainvSalt * cue * stimuli * trial + group_SaltvSugar * cue * stim* trial + (1 + cue\|subject) | | 11050 | 11219 |
| M17: group_PainvSalt * cue * stimuli * trial + group_SaltvSugar * cue * stim* trial + (1 + cue * stimuli\|subject) | | 10636 | 10848 |
| M18: group_PainvSalt * cue * stimuli * trial + group_SaltvSugar * cue * stim* trial + (1 + stimuli *trial\|subject) | | 10834 | 11046 |
| M19: group_PainvSalt * cue * stimuli * trial + group_SaltvSugar * cue * stim* trial + (1 + trial + cue + stimuli\|subject) | | 10782 | 10994 |
| M20: group_PainvSalt * cue * stimuli * trial + group_SaltvSugar * cue * stim* trial +(1 + trial + stimuli\|subject) | | 10833 | 11021 |
| M21: group_PainvSalt * cue * stimuli * trial + group_SaltvSugar * cue * stim* trial + (1 + trial + stimuli\|subject) (1 + cue+stimuli\|subject) | | 10876 | 11087 |
| *Valence ratings* | | | |
| M1: group_PainvSalt + group_SaltvSugar + (1 + 1\|subject) | | 16219 | 16249 |
| M2: group_PainvSalt + group_SaltvSugar + cue + (1 + 1\|subject) | | 15790 | 15827 |
| M3: group_PainvSalt + group_SaltvSugar + cue + stimuli + (1 + 1\|subject) | | 15550 | 15592 |
| M4: group_PainvSalt * stimuli * cue + group_SaltvSugar * stimuli * cue + (1 + 1 \|subject) | | 15437 | 15491 |
| M5: group_PainvSalt + group_SaltvSugar + cue + trial + (1 +1\|subject) | | 15739 | 15781 |
| M6: group_PainvSalt * cue + group_SaltvSugar * cue + stimuli + trial + (1 +1\|subject) | | 15068 | 15128 |
| M7: group_PainvSalt* stimuli + group_SaltvSugar * stimuli + cue + trial + (1 + 1\|subject) | | 14776 | 14836 |
| M8: group_PainvSalt + group_SaltvSugar + cue * stim + trial + (1 + 1\|subject) | | 15437 | 15491 |
| M9: group_PainvSalt * trial + group_SaltvSugar * trial + stimuli + cue + (1 + 1 \|subject) | | 15412 | 15472 |
| M10: group_PainvSalt + group_SaltvSugar + trial * stim + cue + (1 + 1 \|subject) | | 15480 | 15534 |
| M11: group_PainvSalt + group_SaltvSugar + stimuli + cue * trial + (1 + 1 \|subject) | | 15483 | 15537 |
| M12: group_PainvSalt * cue * stimuli + group_SaltvSugar * cue * stimuli + trial + (1 + 1\|subject) | | 14648 | 14739 |
| M13: group_PainvSalt * cue * stimuli * trial + group_SaltvSugar * cue * stim* trial + (1 + 1\|subject) | | 14580 | 14737 |
| M14: group_PainvSalt * cue * stimuli * trial + group_SaltvSugar * cue * stim* trial + (1 + stimuli\|subject) | | 14092 | 14216 |
| M15: group_PainvSalt * cue * stimuli * trial + group_SaltvSugar * cue * stim* trial + (1 + trial\|subject) | | 14429 | 14598 |
| M16: group_PainvSalt * cue * stimuli * trial + group_SaltvSugar * cue * stim* trial + (1 + cue\|subject) | | 14273 | 14442 |
| M17: group_PainvSalt * cue * stimuli * trial + group_SaltvSugar * cue * stim* trial + (1 + cue * stimuli\|subject) | | 13506 | 13718 |
| M18: group_PainvSalt * cue * stimuli * trial + group_SaltvSugar * cue * stim* trial + (1 + stimuli *trial\|subject) | | 13855 | 14066 |
| M19: group_PainvSalt * cue * stimuli * trial + group_SaltvSugar * cue * stim* trial + (1 + trial + cue + stimuli\|subject) | | 13874 | 14086 |
| M20: group_PainvSalt * cue * stimuli * trial + group_SaltvSugar * cue * stim* trial +(1 + trial + stimuli\|subject) | | 13882 | 14070 |
| M21: group_PainvSalt * cue * stimuli * trial + group_SaltvSugar * cue * stim* trial + (1 + cue+stimuli\|subject) | | 14088 | 14299 |
| Regression Model | | **AIC** | **BIC** |
| *Aversiveness: Aversive (heat and salt) and Appetitive (Sugar)* | | | |
| *Intensity ratings* | | | |
| M1: Aversiveness + cue + stimuli + trial + (1 + 1\|subject) | 11875 | | 11917 |
| M2: Aversiveness + cue * stimuli + trial + (1 + 1\|subject) | | 11871 | 11919 |
| M3: Aversiveness * cue + stimuli + trial + (1 +1\|subject) | | 11790 | 11839 |
| M4: Aversiveness * trial + stimuli + cue + (1 + 1 \|subject) | | 11875 | 11923 |
| M5: Aversiveness* stimuli + cue + trial + (1 + 1\|subject) | | 11733 | 11782 |
| M6: Aversiveness + stimuli + cue * trial + (1 + 1 \|subject) | | 11872 | 11921 |
| M7: Aversiveness + trial * stimuli + cue + (1 + 1 \|subject) | | 11866 | 11914 |
| M8: Aversiveness * stimuli + cue * trial + (1 + 1 \|subject) | | 11731 | 11786 |
| M9: Aversiveness * cue * stimuli + trial + (1 + 1\|subject) | | 11691 | 11758 |
| M10: Aversiveness * trial + cue * stimuli + (1 + 1\|subject) | | 11871 | 11925 |
| M11: Aversiveness * cue * trial + stimuli + (1 + 1\|subject) | | 11786 | 11852 |
| M12: Aversiveness + cue * stimuli * trial + (1 + cue\|subject) | | 11864 | 11931 |
| M13: Aversiveness * cue * stimuli * trial + (1 + 1\|subject) | | 11689 | 11798 |
| M14: Aversiveness * cue * stimuli * trial + (1 + stimuli \|subject) | | 11191 | 11312 |
| M15: Aversiveness * cue * stimuli * trial + (1 + cue\|subject) | | 11304 | 11424 |
| M16: Aversiveness * cue * stimuli * trial + (1 + trial\|subject) | | 11587 | 11708 |
| M17: Aversiveness * cue * stimuli * trial + (1 + stimuli+ trial \|subject) | | 11040 | 11179 |
| M18: Aversiveness * cue * stimuli * trial + (1 + cue + trial \|subject) | | 11167 | 11306 |
| M19: Aversiveness * cue * stimuli * trial + (1 + stimuli + cue + trial \|subject) | | 10991 | 11154 |
| M20: Aversiveness * cue * stimuli + trial + (1 + Aversiveness \|subject) | | 11693 | 11814 |
| M21: Aversiveness * cue * stimuli + trial + (1 + Aversiveness + stimuli \|subject) | | 11190 | 11329 |
| M22: Aversiveness * cue * stimuli + trial + (1 + Aversiveness + cue \|subject) | | 11305 | 11444 |
| M23: Aversiveness * cue * stimuli + trial + (1 + Aversiveness + cue+ stimuli \|subject) | | 11149 | 11312 |
| M24: Aversiveness * cue * stimuli + trial + (1 + Aversiveness + cue+ stimuli+ trial \|subject) | | 10993 | 11187 |
| *Valence ratings* | | | |
| M1: Aversiveness + cue + stimuli + trial + (1 + 1\|subject) | | 15488 | 15530 |
| M2: Aversiveness + cue * stimuli + trial + (1 + 1\|subject) | | 15440 | 15488 |
| M3: Aversiveness * cue + stimuli + trial + (1 +1\|subject) | | 15088 | 15137 |
| M4: Aversiveness * trial + stimuli + cue + (1 + 1 \|subject) | | 15489 | 15538 |
| M5: Aversiveness* stimuli + cue + trial + (1 + 1\|subject) | | 14749 | 14816 |
| M6: Aversiveness + stimuli + cue * trial + (1 + 1 \|subject) | | 15441 | 15495 |
| M7: Aversiveness + trial * stimuli + cue + (1 + 1 \|subject) | | 15059 | 15125 |
| M8: Aversiveness * stimuli + cue * trial + (1 + 1 \|subject) | | 15483 | 15532 |
| M9: Aversiveness * cue * stimuli + trial + (1 + 1\|subject) | | 14801 | 14850 |
| M10: Aversiveness * trial + cue * stimuli + (1 + 1\|subject) | | 14799 | 14863 |
| M11: Aversiveness * cue * trial + stimuli + (1 + 1\|subject) | | 15483 | 15532 |
| M12: Aversiveness + cue * stimuli * trial + (1 + cue\|subject) | | 15432 | 15498 |
| M13: Aversiveness * cue * stimuli * trial + (1 + 1\|subject) | | 14745 | 14854 |
| M14: Aversiveness * cue * stimuli * trial + (1 + stimuli \|subject) | | 14268 | 14389 |
| M15: Aversiveness * cue * stimuli * trial + (1 + cue\|subject) | | 14445 | 14566 |
| M16: Aversiveness * cue * stimuli * trial + (1 + trial\|subject) | | 14520 | 14641 |
| M17: Aversiveness * cue * stimuli * trial + (1 + stimuli+ trial \|subject) | | 13972 | 14111 |
| M18: Aversiveness * cue * stimuli * trial + (1 + cue + trial \|subject) | | 14172 | 14311 |
| M19: Aversiveness * cue * stimuli * trial + (1 + stimuli + cue + trial \|subject) | | 13964 | 14127 |
| M20: Aversiveness * cue * stimuli + trial + (1 + Aversiveness \|subject) | | 14748 | 14869 |
| M21: Aversiveness * cue * stimuli + trial + (1 + Aversiveness + stimuli \|subject) | | 14264 | 14403 |
| M22: Aversiveness * cue * stimuli + trial + (1 + Aversiveness + cue \|subject) | | 14441 | 14580 |
| M23: Aversiveness * cue * stimuli + trial + (1 + Aversiveness + cue+ stimuli \|subject) | | 14259 | 14422 |
| M24: Aversiveness * cue * stimuli + trial + (1 + Aversiveness + cue+ stimuli+ trial \|subject) | | 13962 | 14156 |
| Regression Model | | **AIC** | **BIC** |
| *Modality: Heat and Tastes (Salt and Sugar)* | | | |
| *Intensity ratings* | | | |
| M1: Modality + cue + stimuli + trial + (1 + 1\|subject) | 11874 | | 11917 |
| M2: Modality + cue * stimuli + trial + (1 + 1\|subject) | | 11870 | 11919 |
| M3: Modality * cue + stimuli + trial + (1 +1\|subject) | | 11814 | 11862 |
| M4: Modality * trial + stimuli + cue + (1 + 1 \|subject) | | 11812 | 11861 |
| M5: Modality* stimuli + cue + trial + (1 + 1\|subject) | | 11575 | 11642 |
| M6: Modality + stimuli + cue * trial + (1 + 1 \|subject) | | 11809 | 11863 |
| M7: Modality + trial * stimuli + cue + (1 + 1 \|subject) | | 11752 | 11818 |
| M8: Modality * stimuli + cue * trial + (1 + 1 \|subject) | | 11795 | 11843 |
| M9: Modality * cue * stimuli + trial + (1 + 1\|subject) | | 11872 | 11920 |
| M10: Modality * trial + cue * stimuli + (1 + 1\|subject) | | 11792 | 11847 |
| M11: Modality * cue * trial + stimuli + (1 + 1\|subject) | | 11865 | 11973 |
| M12: Modality + cue * stimuli * trial + (1 + cue\|subject) | | 11864 | 11930 |
| M13: Modality * cue * stimuli * trial + (1 + 1\|subject) | | 11552 | 11660 |
| M14: Modality * cue * stimuli * trial + (1 + stimuli \|subject) | | 10940 | 11061 |
| M15: Modality * cue * stimuli * trial + (1 + cue\|subject) | | 11109 | 11230 |
| M16: Modality * cue * stimuli * trial + (1 + trial\|subject) | | 11502 | 11623 |
| M17: Modality * cue * stimuli * trial + (1 + stimuli+ trial \|subject) | | 10838 | 10977 |
| M18: Modality * cue * stimuli * trial + (1 + cue + trial \|subject) | | 11026 | 11166 |
| M19: Modality * cue * stimuli * trial + (1 + stimuli + cue + trial \|subject) | | 10789 | 10952 |
| M20: Modality * cue * stimuli + trial + (1 + Modality \|subject) | | 11547 | 11668 |
| M21: Modality * cue * stimuli + trial + (1 + Modality + stimuli \|subject) | | 10935 | 11074 |
| M22: Modality * cue * stimuli + trial + (1 + Modality + cue \|subject) | | 11107 | 11246 |
| M23: Modality * cue * stimuli + trial + (1 + Modality + cue+ stimuli \|subject) | | 10890 | 11054 |
| M24: Modality * cue * stimuli + trial + (1 + Modality + cue+ stimuli+ trial \|subject) | | 10787 | 10981 |
| *Valence ratings* | | | |
| M1: Modality + cue + stimuli + trial + (1 + 1\|subject) | | 15527 | 15569 |
| M2: Modality + cue * stimuli + trial + (1 + 1\|subject) | | 15479 | 15525 |
| M3: Modality * cue + stimuli + trial + (1 +1\|subject) | | 15368 | 15416 |
| M4: Modality * trial + stimuli + cue + (1 + 1 \|subject) | | 15462 | 15510 |
| M5: Modality* stimuli + cue + trial + (1 + 1\|subject) | | 15184 | 15251 |
| M6: Modality + stimuli + cue * trial + (1 + 1 \|subject) | | 15412 | 15466 |
| M7: Modality + trial * stimuli + cue + (1 + 1 \|subject) | | 15288 | 15355 |
| M8: Modality * stimuli + cue * trial + (1 + 1 \|subject) | | 15283 | 15332 |
| M9: Modality * cue * stimuli + trial + (1 + 1\|subject) | | 15525 | 15573 |
| M10: Modality * trial + cue * stimuli + (1 + 1\|subject) | | 15281 | 15335 |
| M11: Modality * cue * trial + stimuli + (1 + 1\|subject) | | 15522 | 15570 |
| M12: Modality + cue * stimuli * trial + (1 + cue\|subject) | | 15470 | 15537 |
| M13: Modality * cue * stimuli * trial + (1 + 1\|subject) | | 15133 | 15242 |
| M14: Modality * cue * stimuli * trial + (1 + stimuli \|subject) | | 14175 | 14296 |
| M15: Modality * cue * stimuli * trial + (1 + cue\|subject) | | 14594 | 14715 |
| M16: Modality * cue * stimuli * trial + (1 + trial\|subject) | | 15014 | 15136 |
| M17: Modality * cue * stimuli * trial + (1 + stimuli+ trial \|subject) | | 13947 | 14086 |
| M18: Modality * cue * stimuli * trial + (1 + cue + trial \|subject) | | 14418 | 14557 |
| M19: Modality * cue * stimuli * trial + (1 + stimuli + cue + trial \|subject) | | 13944 | 14107 |
| M20: Modality * cue * stimuli + trial + (1 + Modality \|subject) | | 15127 | 15248 |
| M21: Modality * cue * stimuli + trial + (1 + Modality + stimuli \|subject) | | 14168 | 14307 |
| M22: Modality * cue * stimuli + trial + (1 + Modality + cue \|subject) | | 14586 | 14725 |
| M23: Modality * cue * stimuli + trial + (1 + Modality + cue+ stimuli \|subject) | | 14166 | 14329 |
| M24: Modality * cue * stimuli + trial + (1 + Modality + cue+ stimuli+ trial \|subject) | | 13938 | 14132 |

**Supplementary Table 1-2.** Model comparison of linear regression models for medium trials. For each outcome (intensity or valence ratings), our main models included the following mean-centered predictors: Group (dummy coded: Pain vs salt (“group_PainvSalt”; 1 = pain, -1 = salt) and Salt vs sugar (“group_SaltvSugar”, 1 = salt, -1 = sugar), Cue (high cue = 1, low cue = -1), and Trial. We also included random intercepts and slopes per Subject. In addition, we also report model comparison results for models of Aversiveness (Group: aversive versus appetitive) and Modality (Group: pain versus tastes). Factors involved were the same for all three types of models, only with different Group factor. For all models, Akaike information criterion (AIC) and Bayesian information criterion (BIC) were computed to compare models. Smaller AIC/BIC indicates better model fit, adjusting for model complexity. The models highlighted in red were the best fitting models for each type and for each rating. Note that, we only list the models that make sense and do not fail to converge.

| Regression Model | | AIC | BIC |
| --- | --- | --- | --- |
| *Three groups: Heat, Salt, and Sugar* | | | |
| *Intensity ratings* | | | |
| M1: group_PainvSalt + group_SaltvSugar + (1 + 1\|subject) | 4955.9 | | 4982.0 |
| M2: group_PainvSalt + group_SaltvSugar + cue + (1 + 1\|subject) | | 4895.1 | 4926.4 |
| M3: group_PainvSalt + group_SaltvSugar + cue + trial+ (1 + 1\|subject) | | 4894.3 | 4930.7 |
| M4: group_PainvSalt * cue + group_SaltvSugar * cue + (1 +1\|subject) | | 4896.3 | 4938.0 |
| M5: group_PainvSalt * cue + group_SaltvSugar * cue + trial + (1 +1\|subject) | | 4895.4 | 4942.3 |
| M6: group_PainvSalt * cue * trial + group_SaltvSugar * cue * trial + (1 +1\|subject) | | 4876.4 | 4949.4 |
| M7: group_PainvSalt * trial + group_SaltvSugar * trial + cue + (1 + 1 \|subject) | | 4874.6 | 4921.5 |
| M8: group_PainvSalt * trial + group_SaltvSugar * trial + cue + (1 + trial\|subject) | | 4846.9 | 4904.2 |
| M9: group_PainvSalt * trial + group_SaltvSugar * trial + cue + (1 + cue \|subject) | | 4843.2 | 4900.5 |
| M10: group_PainvSalt * trial + group_SaltvSugar * trial + cue + (1 + cue + trial \|subject) | | 4810.5 | 4883.4 |
| M11: group_PainvSalt * trial + group_SaltvSugar * trial + cue + (1 + cue * trial \|subject) | | 4816.8 | 4910.5 |
| *Valence ratings* | | | |
| M1: group_PainvSalt + group_SaltvSugar + (1 + 1\|subject) | | 6119.4 | 6145.4 |
| M2: group_PainvSalt + group_SaltvSugar + cue + (1 + 1\|subject) | | 6108.3 | 6139.6 |
| M3: group_PainvSalt + group_SaltvSugar + cue + trial+ (1 + 1\|subject) | | 6010.0 | 6057.5 |
| M4: group_PainvSalt * cue + group_SaltvSugar * cue + (1 +1\|subject) | | 6109.4 | 6151.1 |
| M5: group_PainvSalt * cue + group_SaltvSugar * cue + trial + (1 +1\|subject) | | 6010.7 | 6057.5 |
| M6: group_PainvSalt * cue * trial + group_SaltvSugar * cue * trial + (1 +1\|subject) | | 5975.6 | 6048.6 |
| M7: group_PainvSalt * trial + group_SaltvSugar * trial + cue + (1 + 1 \|subject) | | 5970.8 | 6017.7 |
| M8: group_PainvSalt * trial + group_SaltvSugar * trial + cue + (1 + trial\|subject) | | 5808.7 | 5866.0 |
| M9: group_PainvSalt * trial + group_SaltvSugar * trial + cue + (1 + cue \|subject) | | 5969.8 | 6027.1 |
| M10: group_PainvSalt * trial + group_SaltvSugar * trial + cue + (1 + cue + trial \|subject) | | 5802.3 | 5875.2 |
| M11: group_PainvSalt * trial + group_SaltvSugar * trial + cue + (1 + cue * trial \|subject) | | 5806.4 | 5900.1 |
| Regression Model | | **AIC** | **BIC** |
| *Aversiveness: Aversive (heat and salt) and Appetitive (Sugar)* | | | |
| *Intensity ratings* | | | |
| M1: cue + (1 + 1\|subject) | 4908.7 | | 4929.5 |
| M2: trial + (1 + 1\|subject) | | 4968.3 | 4989.1 |
| M3: Aversiveness + (1 +1\|subject) | | 4971.4 | 4992.2 |
| M4: Aversiveness + cue + (1 + 1 \|subject) | | 4910.6 | 4936.6 |
| M5: Aversiveness + trial + (1 + 1\|subject) | | 4970.2 | 4996.2 |
| M6: cue + trial + (1 + 1 \|subject) | | 4907.8 | 4933.9 |
| M7: Aversiveness + trial + cue + (1 + 1 \|subject) | | 4909.7 | 4941.0 |
| M8: Aversiveness * cue + trial + (1 + 1 \|subject) | | 4911.4 | 4947.9 |
| M9: Aversiveness + cue * trial + (1 + 1\|subject) | | 4910.2 | 4946.6 |
| M10: Aversiveness * trial + cue + (1 + 1\|subject) | | 4911.4 | 4947.9 |
| M11: Aversiveness * cue * trial + (1 + 1\|subject) | | 4912.8 | 4964.9 |
| M12: cue + (1 + cue \|subject) | | 4878.9 | 4910.1 |
| M13: cue + (1 + trial \|subject)) | | 4857.4 | 4888.7 |
| M14: cue + (1 + cue + trial \|subject)) | | 4821.3 | 4868.2 |
| M15: cue + (1 + cue * trial \|subject)) | | 4826.2 | 4894.0 |
| *Valence ratings* | | | |
| M1: cue + (1 + 1\|subject) | | 6140.9 | 6161.8 |
| M2: trial + (1 + 1\|subject) | | 6053.3 | 6074.1 |
| M3: Aversiveness + (1 +1\|subject) | | 6124.8 | 6145.6 |
| M4: Aversiveness + cue + (1 + 1 \|subject) | | 6113.7 | 6139.8 |
| M5: Aversiveness + trial + (1 + 1\|subject) | | 6026.1 | 6052.1 |
| M6: cue + trial + (1 + 1 \|subject) | | 6042.6 | 6068.6 |
| M7: Aversiveness + trial + cue + (1 + 1 \|subject) | | 6015.3 | 6046.6 |
| M8: Aversiveness * cue + trial + (1 + 1 \|subject) | | 6015.7 | 6052.2 |
| M9: Aversiveness + cue * trial + (1 + 1\|subject) | | 6017.3 | 6053.8 |
| M10: Aversiveness * trial + cue + (1 + 1\|subject) | | 6017.3 | 6053.8 |
| M11: Aversiveness * cue * trial + (1 + 1\|subject) | | 6020.4 | 6072.5 |
| M12: Aversiveness + trial + cue + (1 + cue \|subject) | | 6015.3 | 6056.9 |
| M13: Aversiveness + trial + cue + (1 + trial \|subject)) | | 5823.5 | 5865.2 |
| M14: Aversiveness + trial + cue + (1 + cue * trial \|subject) | | 5821.7 | 5899.9 |
| M15: Aversiveness + trial + cue + (1 + cue + trial \|subject) | | 5818.0 | 5875.3 |
| M16: Aversiveness + trial + cue + (1 + Aversiveness \|subject) | | 6015.4 | 6057.1 |
| M17: Aversiveness + trial + cue + (1 + Aversiveness + trial \|subject) | | 5825.6 | 5882.9 |
| M18: Aversiveness + trial + cue + (1 + Aversiveness + cue \|subject) | | 6016.5 | 6073.8 |
| M19: Aversiveness + trial + cue + (1 + Aversiveness + trial +cue \|subject) | | 5821.1 | 5899.2 |
| M20: Aversiveness + trial + cue + (1 + Aversiveness * cue \|subject) | | 6022.7 | 6100.9 |
| Regression Model | | **AIC** | **BIC** |
| *Modality: Heat and Tastes (Salt and Sugar)* | | | |
| *Intensity ratings* | | | |
| M1: cue + (1 + 1\|subject) | 4908.7 | | 4929.5 |
| M2: trial + (1 + 1\|subject) | | 4968.3 | 4989.1 |
| M3: Modality + (1 +1\|subject) | | 4959.6 | 4980.5 |
| M4: Modality + cue + (1 + 1 \|subject) | | 4898.9 | 4924.9 |
| M5: Modality + trial + (1 + 1\|subject) | | 4958.4 | 4984.5 |
| M6: cue + trial + (1 + 1 \|subject) | | 4907.8 | 4933.9 |
| M7: Modality + trial + cue + (1 + 1 \|subject) | | 4898.0 | 4929.3 |
| M8: Modality * cue + trial + (1 + 1 \|subject) | | 4897.2 | 4933.7 |
| M9: Modality + cue * trial + (1 + 1\|subject) | | 4898.5 | 4934.9 |
| M10: Modality * trial + cue + (1 + 1\|subject) | | 4883.4 | 4919.8 |
| M11: Modality * cue * trial + (1 + 1\|subject) | | 4884.7 | 4936.8 |
| M12: Modality + trial + cue + (1 + cue \|subject) | | 4852.3 | 4899.2 |
| M13: Modality + trial + cue + (1 + trial \|subject)) | | 4849.6 | 4896.5 |
| M14: Modality + trial + cue + (1 + cue * trial \|subject) | | 4879.7 | 4926.6 |
| M15: Modality + trial + cue + (1 + cue + trial \|subject) | | 4813.3 | 4875.8 |
| M16: Modality + trial + cue + (1 + Modality \|subject) | | 4849.0 | 4911.6 |
| M17: Modality + trial + cue + (1 + Modality + trial \|subject) | | 4850.5 | 4913.0 |
| M18: Modality + trial + cue + (1 + Modality + cue \|subject) | | 4814.5 | 4897.9 |
| M19: Modality + trial + cue + (1 + Modality + trial +cue \|subject) | | 4817.7 | 4901.1 |
| M20: Modality + trial + cue + (1 + Modality * cue \|subject) | | 4859.7 | 4943.1 |
| *Valence ratings* | | | |
| M1: cue + (1 + 1\|subject) | | 6140.9 | 6161.8 |
| M2: trial + (1 + 1\|subject) | | 6053.3 | 6074.1 |
| M3: Modality + (1 +1\|subject) | | 6153.7 | 6174.6 |
| M4: Modality + cue + (1 + 1 \|subject) | | 6142.7 | 6168.7 |
| M5: Modality + trial + (1 + 1\|subject) | | 6055.0 | 6081.1 |
| M6: cue + trial + (1 + 1 \|subject) | | 6042.6 | 6068.6 |
| M7: Modality + trial + cue + (1 + 1 \|subject) | | 6044.3 | 6075.6 |
| M8: Modality * cue + trial + (1 + 1 \|subject) | | 6043.3 | 6079.8 |
| M9: Modality + cue * trial + (1 + 1\|subject) | | 6046.3 | 6082.8 |
| M10: Modality * trial + cue + (1 + 1\|subject) | | 6011.2 | 6047.6 |
| M11: Modality * cue * trial + (1 + 1\|subject) | | 6013.7 | 6065.8 |
| M12: trial + cue + (1 + cue \|subject) | | 6042.0 | 6078.5 |
| M13: trial + cue + (1 + trial \|subject)) | | 5851.0 | 5887.5 |
| M14: trial + cue + (1 + cue + trial \|subject) | | 5844.9 | 5897.0 |
| M15: trial + cue + (1 + cue * trial \|subject) | | 5846.9 | 5919.8 |

**Table S1-3.** Model comparison of linear regression models for expectation. For each outcome (expected intensity or expected valence ratings), our main models included the following mean-centered predictors: Group (dummy coded: Pain vs salt (“group_PainvSalt”; 1 = pain, -1 = salt) and Salt vs sugar (“group_SaltvSugar”, 1 = salt, -1 = sugar), Cue (high cue = 1, low cue = -1), and Block. We also included random intercepts and slopes per Subject. We also included random intercepts and slopes per Subject. In addition, we also report model comparison results for models of Aversiveness (Group: aversive versus appetitive) and Modality (Group: pain versus tastes). Factors involved were the same for all three types of models, only with different Group factor. For all models, Akaike information criterion (AIC) and Bayesian information criterion (BIC) were computed to compare models. Smaller AIC/BIC indicates better model fit, adjusting for model complexity. The models highlighted in red were the best fitting models for each type and for each rating. Note that, we only list the models that make sense and do not fail to converge.

| Regression Model | | AIC | BIC |
| --- | --- | --- | --- |
| *Three groups: Heat, Salt, and Sugar* | | | |
| *Expected Intensity ratings* | | | |
| M1: group_PainvSalt + group_SaltvSugar + cue + block + (1 + 1\|subject) | 2225.9 | | 2256.2 |
| M2: group_PainvSalt + group_SaltvSugar + cue * block + (1 + 1\|subject) | | 2226.3 | 2261.0 |
| M3: group_PainvSalt * cue + group_SaltvSugar * cue + block + (1 + 1\|subject) | | 2183.8 | 2222.7 |
| M4: group_PainvSalt * block + group_SaltvSugar * block + cue + (1 + 1\|subject) | | 2223.0 | 2261.9 |
| M5: group_PainvSalt * cue * block + group_SaltvSugar * cue * block + (1 + 1\|subject) | | 2183.9 | 2244.5 |
| M6: group_PainvSalt * cue * block + group_SaltvSugar * cue + (1 + 1\|subject) | | 2184.8 | 2219.4 |
| *Expected Valence ratings* | | | |
| M1: group_PainvSalt + group_SaltvSugar + cue + block + (1 + 1\|subject) | | 2848.9 | 2879.2 |
| M2: group_PainvSalt + group_SaltvSugar + cue * block + (1 + 1\|subject) | | 2849.7 | 2884.3 |
| M3: group_PainvSalt * cue + group_SaltvSugar * cue + block + (1 + 1\|subject) | | 2726.6 | 2765.6 |
| M4: group_PainvSalt * block + group_SaltvSugar * block + cue + (1 + 1\|subject) | | 2836.3 | 2875.2 |
| M5: group_PainvSalt * cue * block + group_SaltvSugar * cue * block + (1 + 1\|subject) | | 2712.7 | 2773.3 |
| M6: group_PainvSalt * cue * block + group_SaltvSugar * cue + (1 + 1\|subject) | | 2742.9 | 2777.5 |
| Regression Model | | **AIC** | **BIC** |
| *Aversiveness: Aversive (heat and salt) and Appetitive (Sugar)* | | | |
| *Expected Intensity ratings* | | | |
| M1: Aversiveness+ cue + block + (1 + 1\|subject) | 2237.7 | | 2263.6 |
| M2: Aversiveness+ cue * block + (1 + 1\|subject) | | 2238.1 | 2268.3 |
| M3: Aversiveness* cue + block + (1 + 1\|subject) | | 2199.5 | 2229.8 |
| M4: Aversiveness * block + cue + (1 + 1\|subject) | | 2237.7 | 2267.9 |
| M5: Aversiveness* cue * block + (1 + 1\|subject) | | 2101.1 | 2244.4 |
| M6: cue * block + (1 + 1\|subject) | | 2100.4 | 2226.4 |
| *Expected Valence ratings* | | | |
| M1: Aversiveness+ cue + block + (1 + 1\|subject) | | 2895.8 | 2921.8 |
| M2: Aversiveness+ cue * block + (1 + 1\|subject) | | 2896.5 | 2926.8 |
| M3: Aversiveness* cue + block + (1 + 1\|subject) | | 2849.6 | 2879.8 |
| M4: Aversiveness * block + cue + (1 + 1\|subject) | | 2888.7 | 2919.0 |
| M5: Aversiveness* cue * block + (1 + 1\|subject) | | 2843.7 | 2886.9 |
| M6: cue * block + (1 + 1\|subject) | | 2863.0 | 2889.0 |
| Regression Model | | **AIC** | **BIC** |
| *Modality: Heat and Tastes (Salt and Sugar)* | | | |
| *Expected Intensity ratings* | | | |
| M1: Modality + cue + block + (1 + 1\|subject) | 2234.7 | | 2260.7 |
| M2: Modality + cue * block + (1 + 1\|subject) | | 2235.1 | 2265.4 |
| M3: Modality * cue + block + (1 + 1\|subject) | | 2210.5 | 2240.8 |
| M4: Modality * block + cue + (1 + 1\|subject) | | 2235.2 | 2265.5 |
| M5: Modality * cue * block + (1 + 1\|subject) | | 2213.2 | 2256.4 |
| M6: Modality * block + (1 + 1\|subject) | | 2111.3 | 2237.3 |
| *Expected Valence ratings* | | | |
| M1: Modality + cue + block + (1 + 1\|subject) | | 2851.7 | 2877.6 |
| M2: Modality + cue * block + (1 + 1\|subject) | | 2852.4 | 2882.7 |
| M3: Modality * cue + block + (1 + 1\|subject) | | 2732.7 | 2763.0 |
| M4: Modality * block + cue + (1 + 1\|subject) | | 2852.8 | 2883.1 |
| M5: Modality * cue * block + (1 + 1\|subject) | | 2735.8 | 2779.1 |
| M6: Modality * block + (1 + 1\|subject) | | 2748.6 | 2774.6 |
